# “Generation of morphologically distinct astrocyte subtypes from human iPSCs reveals a link between glial shape, function and reactivity”

**DOI:** 10.1101/2024.11.11.622998

**Authors:** Kelly O’Toole, Ludovica Guetta, Paulina Urbanaviciute, Pacharaporn Suklai, Chandler Wright, Theodore Fisher, Shane Liddelow, Raphaelle Luisier, Sarah J Marzi, Andrea Serio

## Abstract

Astrocyte morphology *in vivo* is heterogeneous across different subtypes and dynamically changes in response to various stimuli. However, several questions on the mechanistic links between shape and function remain unanswered. Here, we developed an efficient protocol to generate pure populations of morphologically distinct human astrocytes *in vitro*, which we used for a systematic analysis of shape-function relationships. We performed a structural, molecular, and functional characterization of these populations and highlighted how their distinct morphologies mirror distinct functional and transcriptional patterns at the population level. We were also able to both correlate gene expression profiles of these morphologically distinct astrocyte subtypes with *in vivo* astrocytes in the human brain, and to validate our findings with primary isolated murine astrocytes in vitro. Moreover, we show that the observed morphological differences are correlated with changes in key cytoskeletal proteins, which offers a potential link to the observed functional differences. Finally, we demonstrated that different morphological subtypes of astrocytes have distinct reactivity responses to a common stimulus. This study offers a glimpse into the shape-function dynamics of human astrocytes, highlighting potential mechanistic links between cytoskeletal usage and astrocyte function, while also providing tools and datasets that will be useful for further studies into human glial biology in health and disease.

## Introduction

Every cell is configured to acquire an optimal shape which is tailored to its specific function and tasks within an organism or tissue. This, in turn, can influence cellular metabolism and homeostasis. This link between shape and function is a fundamental aspect of cellular biology^1–3^, and has profound implications in our understanding of pathological mechanisms^4^. Moreover, this is a dynamic relationship, as morphology can change not only across cell types, but also over space and time; different subtypes of the same cell populations can vary widely depending on their regional specification, like neurons and muscle cells, and their shape can change significantly in response to different stimuli and conditions^5,6^.

Astrocytes^7,8^ have a highly ramified morphology, that changes across each regional and functional subtype^7–10^. These glial cells play a crucial role in the development and function of the central nervous system (CNS)^11^, and their diversity of form allows them to perform a plethora of key functions: tiling the brain and spinal cord in vast astrocytic networks, contacting other cell types, sensing and responding to their local microenvironment, and even acting as master regulators of metabolism for the brain and spinal cord^9,10^. The complexity of astrocyte morphology dynamically responds to injury, inflammation, and various neurological disorders^12,13^. The observed correlations between altered astrocyte shape and pathological processes and the mechanisms underlying astrocyte shape changes in neurodegenerative disorders remain poorly understood.

Recent advances in induced pluripotent stem cell (iPSC) reprogramming^14^, *in vitro* differentiation and screening ^15^, sequencing^16^, and organoids^17^ have provided a plethora of ways to more accurately model human astrocytes in health and disease. These improvements have been critical for understanding the pathogenesis of various neurodegenerative disorders^18–20^. However, lots of unanswered questions remain on how astrocyte morphology is determined, as *in vitro* astrocytes often lack the distinctive complexity of their *in vivo* counterparts^21^, and in some protocols the cultures present high morphological variability^22^. Some of these limitations are partly due to the low efficiency of astrocyte differentiation protocols, which often use undefined factors (e.g. serum), or mixes of cytokines and growth factors with diverging effects (e.g. epidermal growth factor [EGF], fibroblastic growth factor-2 [FGF2], bone morphogenic [BMP4], ciliary neurotrophic factor [CNTF]).

In response to these challenges, we developed a highly efficient, chemically defined protocol to generate pure populations of functional astrocytes from human iPSC cultures, in a reliable and scalable manner. Our approach uses a minimal set of cytokines, optimised to generate distinct morphological subtypes of astrocytes *in vitro*. This system allowed us to obtain pure populations of human astrocytes with distinct morphologies, which we characterised at the population, cellular, and subcellular level, demonstrating that morphology correlates with subtle differences in functionality, as well as distinct transcriptional profiles.

These insights will be key to both enhance our understanding of mechanisms underlying astrocyte functionality and better model disease *in vitro*.

## Results

### BMP4 and CNTF based differentiation produces human astrocytes with distinct morphologies

To investigate the links between morphology and function in human astrocytes we first sought to establish a reproducible protocol that would generate astrocytes with distinct morphologies in a controlled fashion, starting from common glial progenitors.

We adapted a previously developed protocol to derive progenitor cells committed to the astrocytic fate (astrocyte progenitor cells, APCs) starting from iPSC populations^19^. (**Figure 1A**). These APCs are derived in completely defined conditions and can be propagated as homogeneous monolayers for several passages in a combination of FGF2 and EGF (**Figure 1B**), offering an ideal system for large scale studies. These astrocyte progenitors express *NES* (Nestin) and *NF1A*^23,24^ (**Figure 1B**), and importantly, once expanded, can be differentiated into high purity (>95%) astrocytes that retain important astrocyte functions (e.g. glutamate uptake, ability to support synaptogenesis). These and have already been employed to assess different disease phenotypes^19^, demonstrating that they can mimic several aspects of astrocyte biology. Using this system, we initially aimed to test whether exposing the same population of iPSC-derived APCs to known astrogenic factors might produce morphologically distinct terminally differentiated astrocytes. Based on previous evidence from both *ex vivo*^25,26^ and mouse in vitro models^16^, we selected BMP4 and CNTF as potential “polarizing” factors for astrocyte morphology.

**Figure 1-.**
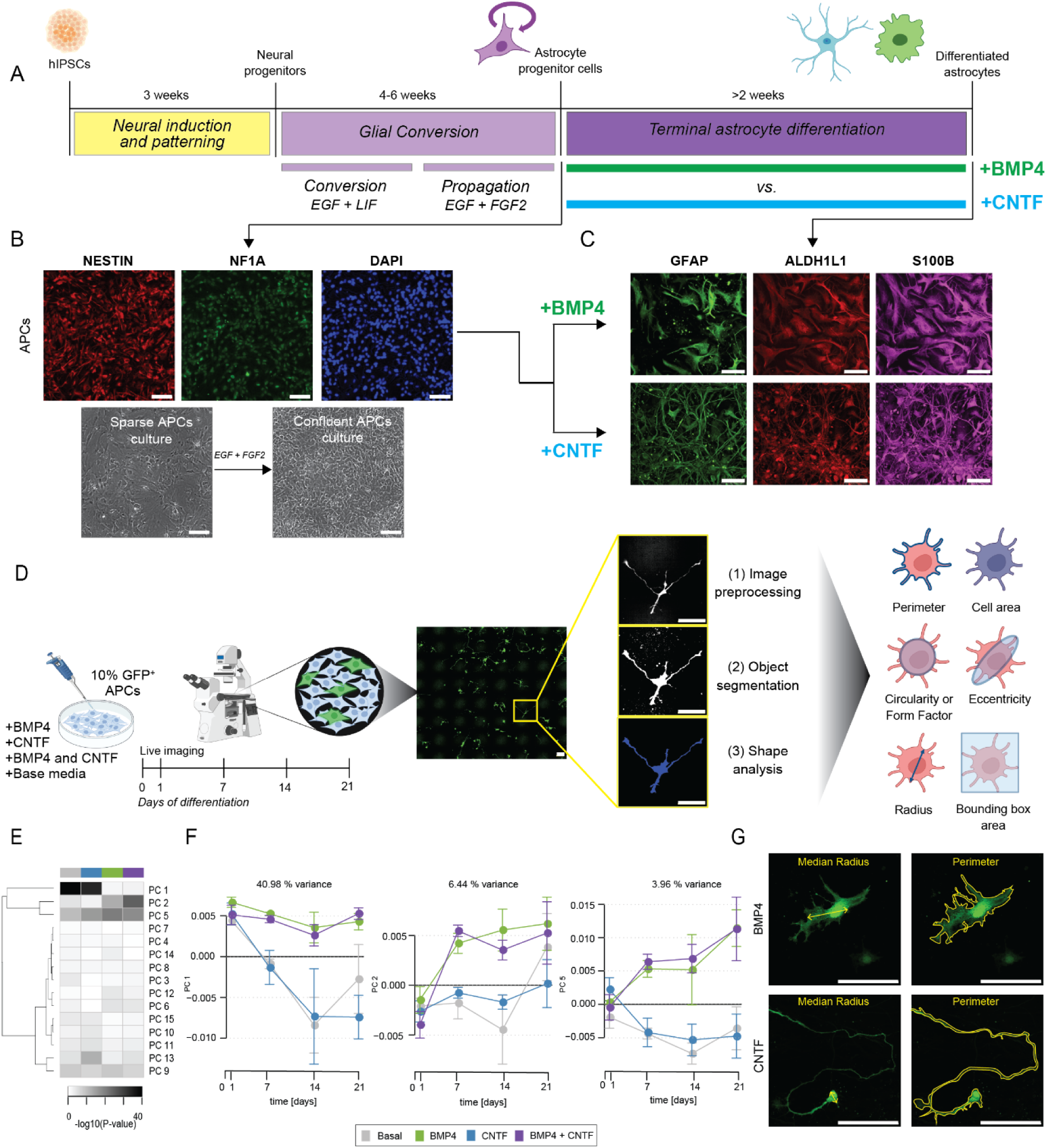
BMP4 and CNTF produce human astrocyte subtypes with distinct morphologies. A) Schematic overview of the protocols for the generation of human iPSC-derived astrocytes terminally differentiated with BMP4 or CNTF. Astrocyte progenitors are generated and propagated as previously described (Serio et al, 2013) and then terminally differentiated for two weeks in the presence of BMP4 or CNTF. B) Representative images of astrocyte progenitors stained for NESTIN, NF1A and nuclei (DAPI) and brightfield pictures showing sparse and confluent progenitor cultures. Scale bar: 100 μm. C) Representative images of differentiated BMP4 and CNTF astrocytes stained for astrocytic markers GFAP, ALDH1L1 and S100β. Scale bar: 100 μm. D) Pipeline for the analysis of astrocyte morphological characteristics. Astrocyte progenitor cells (APCs) are sparsely transfected with green fluorescent protein (GFP) under the EEF1A1 promoter, differentiated with BMP4, CNTF, base media or a combination of cytokines and then imaged live over differentiation. The images acquired are pre-processed, followed by object segmentation using pixel classification. Morphometric analysis is then performed on these objects to obtain key morphometric parameters. Scale bar: 50 μm. E) Principal component analysis of astrocyte morphological variations. The principal components cluster with unsupervised hierarchal clustering according to cytokine groups. F) Representation of the top 3 principal components (PC1, PC2, PC5) which explain over 50% of the variance and fluctuate over the days of differentiation for the different cytokine groups. Data represents averages of wells from 1 control line in 1 experimental block, with 7 technical replicates (n=7 wells). G) Graphical representations (yellow) of shape descriptors, median radius, and perimeter, superimposed onto BMP4 and CNTF GFP^+^ astrocytes. Scale bar: 100 μm.Astrocyte shape is committed during the initial phases of differentiation

We adapted the established protocol^19^ to include a differentiation with medium supplemented with either BMP4 or CNTF for two weeks (**Figure 1A**). We found that both protocols produce non-proliferative (**Figure S1A and S1B**) cells that are immunoreactive for key astrocytic markers, such as Aldh1l1 and S100β^27,28^ (**Figure 1C**). However, the resultant populations of astrocytes have distinct morphologies which are homogeneous within each population. Specifically, BMP4-differentiated astrocytes display larger and more spread-out cell bodies with a rounder shape compared to CNTF-differentiated astrocytes, which are characterized by small cell bodies with long thin processes (**Figure 1C**).

To better quantify these morphological differences and define the timescale of development of astrocyte morphology, we performed high content live imaging of astrocyte progenitors sparsely transfected with GFP under the *EEF1A1* promoter at different timepoints (Days 0, 1, 7, 14, and 21) during the differentiation process. We compared astrocytes differentiated with BMP4, CNTF, both cytokines together, or base medium devoid of any cytokine (**Figure 1D**). To assess the morphological heterogeneity across these conditions in an unbiased manner, we established a computational pipeline for semi-automated single-cell profiling based on discrete morphological features over the time-course of terminal differentiation. We first applied singular value decomposition (SVD) to the matrix of single-cell measurements (52 measurements across 5,376 cells) to identify major orthogonal temporal dynamics of morphological changes associated with terminal differentiation of astrocytes exposed to different cytokines^29^. Our analysis reveals that the first 20 components capture 90% of the variance in the data (**Figure 1E**). Using linear mixed modelling, we found significant associations between principal component (PC) 1 (41% of variance), PC2 (6.4% of variance), and PC5 (4% of variance) with the different cytokine combinations over time (**Figure 1F**). Notably, our results indicate that exposure to BMP4, irrespective of CNTF treatment, leads to diverging differentiation trajectories. In contrast, astrocytes cultured with CNTF or basal medium show similar progressive changes over time, with the exception of a notable difference observed at day 21. Further analysis of the morphological parameters most contributing and correlating with PC1, PC2, and PC5 revealed that measurements describing rounder and more extended morphologies (e.g., median radius and extent) most positively contribute to PC1, whereas measurements such as cell perimeter and compactness, which describe more complex, ramified morphologies, most contribute negatively to PC1 (**Figure S1C**). This indicates that the terminal differentiation of astrocytes in the absence of BMP4 leads to a progressive reduction in roundness and extent, along with a progressive increase in ramification (**Figure 1G**).

To evaluate the difference in morphology between BMP4 and CNTF, we selected median radius and perimeter (**Figure 1G and S1D**) which show opposite direction of change. The cells treated with BMP4 or with the combination of both cytokines show a gradual increase in median radius over three weeks of differentiation, but no change in perimeter – suggesting an overall simplification of morphology with time. Conversely, CNTF and basal medium astrocytes show an increase in their perimeter as they differentiate but no appreciable in median radius – suggesting a switch to a more complex cell morphology (**Figure S1D**).

Overall, this demonstrates that BMP4 and CNTF specifically influence human astrocyte morphology *in vitro* producing homogeneous populations with opposing cell shapes. In addition, we show that the effect of BMP4 can overpower that of CNTF suggesting it is a stronger morphogen (**Figure 1E-F and S1D**), while addition of CNTF maintains a base non-divergent morphological differentiation as astrocytes supplemented with CNTF mimic those grown in basal differentiation medium alone, with the exception of slight differences in cell body shape (**Figure S1E-F**).

We next sought to investigate whether the distinct morphological phenotypes obtained with BMP4 and CNTF are the result of the commitment to a specific developmental program, and therefore stable once acquired, or plastic and changing based on a constant supply of cytokines.

We therefore performed medium switch experiments (**Figure 2A**), in which we first swapped 2-week differentiated astrocytes to the opposite cytokine regimen for an additional week of differentiation, to test whether this would alter their final morphology to resemble that of the opposite subtype. We found that cytokine switching did not correlate with a switch of morphologies between the two subtypes, with BMP4 astrocytes showing no significant alteration in any of the shape descriptors we evaluated. While CNTF astrocytes showed minimal changes in median radius and compactness compared to their non-switched counterpart, this did not alter the overall architecture of the cells, nor induce them to adopt the same morphology as BMP4 astrocytes (**Figure 2B**). These results suggest that astrocytes in our model commit to a specific morphology by two weeks of differentiation, albeit with the potential of morphological malleability at earlier timepoints through exposure to different cytokines.

**Figure 2-.**
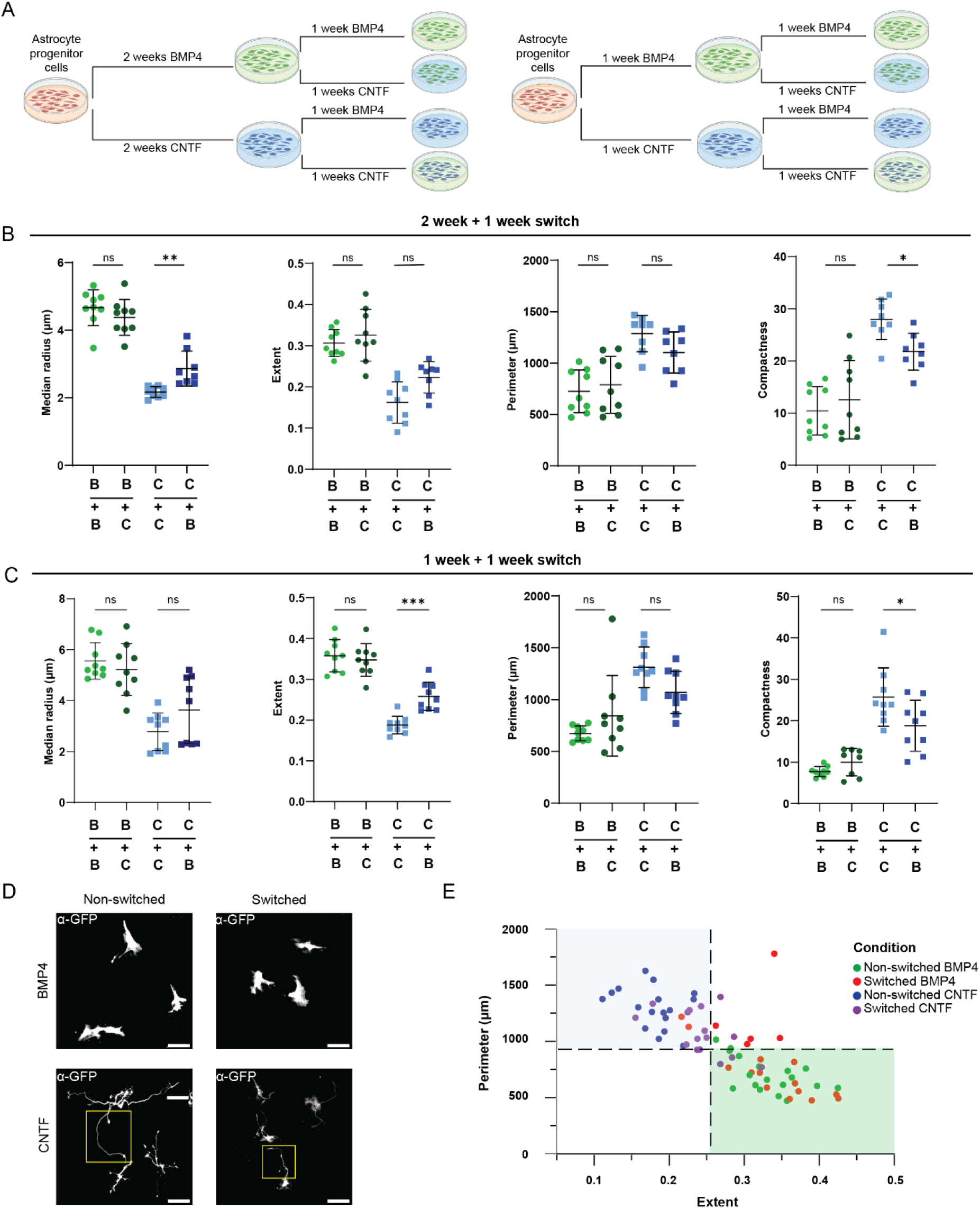
Astrocyte morphology is largely committed during the first week of differentiation. A) Schematic overview of the experimental set up for cytokine switching experiments, with astrocytes differentiating for 1 or 2 weeks with BMP4 switched to CNTF and astrocytes differentiating for 1 or 2 weeks with CNTF switched to BMP4, in all cases for one additional week. B-C) Quantification of morphological differences between BMP4 and CNTF GFP^+^ astrocytes differentiated for either 2 weeks (B) or 1 week (C) with one cytokine and then ‘switched’ (B+C or C+B) or ‘non-switched’ (B+B or C+C). One-way ANOVA or Kruskall-Wallis test with Dunn’s multiple comparisons to test for significance between switched and non-switched pairs. Data represents technical replicate well averages of astrocytes from 2 control lines from 2 independent experiment blocks (n=7-9 wells) mean ± SD. ****p<0.0001, ***p<0.001, **p<0.01, *p<0.05, ns: non-significant. B+B: BMP4 + BMP4; B+C: BMP4 + CNTF; C+C: CNTF+CNTF; C+B: CNTF+BMP4. D) Representative fluorescent images (20x) of non-switched BMP4 (B+B) and switched BMP4 (B+C) astrocytes, and non-switched CNTF (C+C) and switched CNTF (C+B) astrocytes. Inset in CNTF panels highlights the process length differences between non-switched and switched CNTF astrocytes. Scale bar: 100μm. E) 2D scatter plot of perimeter versus extent to visualise the correlation between these two shape descriptors, with the data points representing well averages of BMP4 and CNTF non-switched versus switched astrocytes. The dotted line on the y-axis is the median of control non-switched CNTF astrocytes and the dotted line on the x-axis is the median of control non-switched BMP4 astrocytes. The top left quadrant contains cells with a ‘CNTF’ phenotype, and the bottom right quadrant contains cells with a ‘BMP4’ phenotype. Data points represent well averages of 2 control lines across 4 experimental blocks, from both switching experiments.

To test this hypothesis, we repeated the medium-switching experiment after one week of differentiation. BMP4 astrocytes switched to CNTF medium showed no significant differences from non-switched BMP4 controls, while CNTF astrocytes switched to BMP4 exhibited increased extent and reduced compactness (**Figure 2C**). Qualitatively, it seems that Qualitative assessment suggests that these differences primarily result from to the reduced process length observed in comparison to non-switched CNTF (**Figure 2D**). These alterations impact overall perimeter and associated descriptors without altering the global architecture, defined by a smaller cell body and elongated, thin processes.

These experiments indicate that the contrasting morphologies of BMP4 and CNTF astrocytes, characterised by opposing extent and perimeter (i.e. high extent/low perimeter for BMP4 and low extent/high perimeter for CNTF), are largely committed established after 1-week of differentiation. While minor adjustments can be made, each subtype broadly maintains its morphology irrespective of culture conditions (**Figure 2E**). Notably, BMP4 appears to exert a dominant effect, consistent with observations from astrocytes differentiating in the presence of both cytokines (**Figure 1E-F and S1D**).

### BMP4 and CNTF astrocytes show distinct homeostatic capabilities

We next sought to determine whether the distinct morphologies of CNTF and BMP4 astrocytes also translate into distinct functional properties.

Calcium signalling via intercellular calcium waves (ICWs) is key in regulating astrocytic network-level communication^30^, as well as modulating interactions with neurons^31^. As ICWs rely on calcium microdomains within astrocytic processes for propagation^32,33^, we hypothesized that the difference in morphology between BMP4 and CNTF astrocytes may also correlated with different calcium wave dynamics. To test this, we stimulated BMP4 and CNTF astrocyte populations with adenosine triphosphate (ATP; 5mM) and used Fluo-4AM calcium dye to visualize the calcium signalling response over a period of 160 seconds (**Figure S3A**). CNTF astrocytes responded more swiftly to ATP administration compared to BMP4, during imaging (**Figure 3A**), indicating different kinetics associated with the two subtypes, which we confirmed by peak analysis (**Figure 3B**). Following ATP stimulation, CNTF astrocytes exhibited a significantly shorter time to peak, whereas BMP4 astrocytes displayed significantly higher and broader peak responses (**Figure 3C**). This indicates that, while BMP4 astrocytes are slower in responding, they show a higher and sustained concentration of calcium within the cell during the spiking activity. Interestingly, this difference in ICW kinetics does appears independent of Connexin-43 protein levels, which are comparable between the two astrocyte populations (**Figure 3D** and **S3B**). However, one possible explanation that we cannot exclude is that CNTF astrocytes might simply respond faster to the bath application of ATP, independently of connexin-based propagation.

**Figure 3-.**
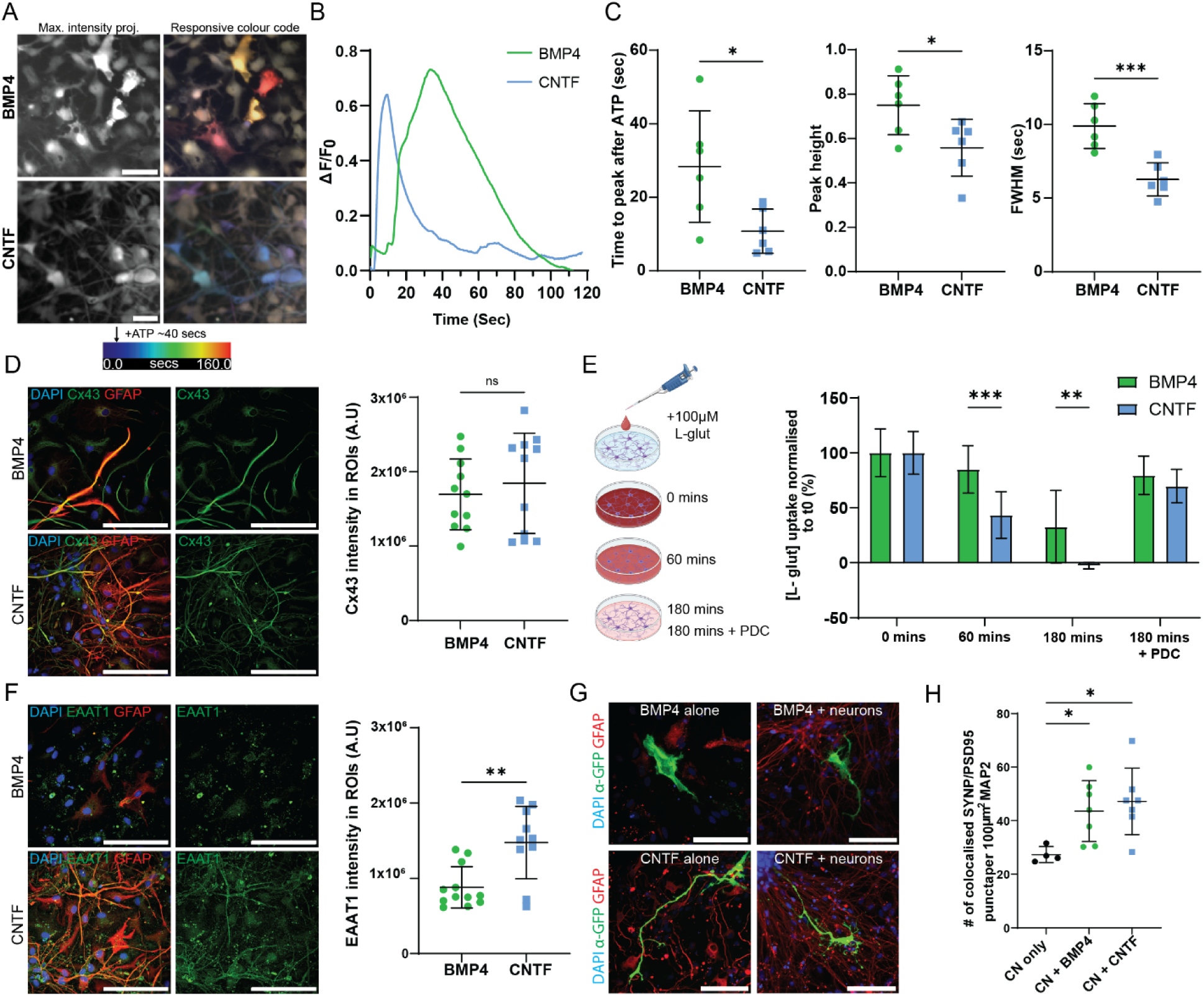
BMP4 and CNTF astrocytes display different homeostatic functional profiles. A-C) BMP4 and CNTF astrocytes have different calcium wave kinetics. A) Temporal colour-coding analysis. Time lapse images of astrocytes loaded with calcium-sensitive indicator Fluo4-AM and stimulated with ATP to initiate ATP-responsive calcium waves were taken. Stacks were processed with temporal color-coding to show at what frame after ATP application (frame 50) the highest peak in intensity occurs for each cell. White indicates cells that did not change intensity, whilst cells coloured in blue spiked earlier than those coloured in yellow-red. Scale bar: 100 μm. B-C) Quantification of calcium spike characteristics (FWHM: full width half maximum). Data represents well averages of 2 control lines from 2 experimental blocks with 6 technical replicates total, median ± IQR. Unpaired t-tests or Mann-Whitney tests were used to test for significance. ****p<0.0001, ***p<0.001, *p<0.05. D) Representative images of BMP4 and CNTF astrocytes stained for connexin-43 (CX43), nucleus (DAPI) and intermediate filaments (GFAP) and quantification of CX43 fluorescence intensity within regions of interest (ROIs). Data points represent well averages of 2 control lines from 2 experimental blocks with 10 technical replicates (n=10-11 wells), mean ± SD. Unpaired t-test was used to test for significance. **p<0.01, ns: non-significant. Scale bar: 100 μm. E-F) CNTF astrocytes have faster glutamate uptake dynamics. E) In vitro fluorometric glutamate (L-glut) uptake assay protocol was performed using inhibitor L-trans-pyrrolidine-2,4-dicarboxylate (PDC) to block glutamate flux through glutamate transporters in the 180-minute incubation. Quantification of fluorescence corresponding to known concentrations of glutamate was performed and normalised to the number of cells and to baseline (t0) control. Data points represent well averages of 2 control lines from 2 experimental blocks with 7-8 technical replicates (n=7-8 wells), mean ± SD. Two-way ANOVA with Šídák’s multiple comparisons to test for significance between BMP4 versus CNTF astrocytes at different timepoints. F) Representative images of BMP4 and CNTF astrocytes stained for glutamate transporter (EAAT1), nucleus (DAPI) and intermediate filaments (GFAP) and quantification of EAAT1 fluorescence intensity within regions of interest (ROIs). Data points represent well averages of 2 control lines from 2 experimental blocks with 10 technical replicates (n=10-11 wells), mean ± SD. Unpaired t-test was used to test for significance. **p<0.01, ns: non-significant. Scale bar: 100μm. G) Representative fluorescent images of GFP^+^ BMP4 and CNTF astrocytes cultured alone or with human stem-cell-derived motor neurons (BMP4 + neurons, CNTF + neurons) and stained with α-GFP, nuclear staining (DAPI) and neuronal tubulin (β3-Tub) or astrocyte intermediate filaments (GFAP). Scalebar: 100 μm. H) Quantification of the number of colocalised SYN and PSD95 puncta per 100 μm^2^ MAP2 positive dendrite areas in cortical neurons cultured alone (CN only) or in the presence of BMP4 (CN+BMP4 ACs) or CNTF (CN+CNTF ACs) astrocytes. Datapoints represent well averages of 2 control lines (for astrocytes) for 2 experimental blocks with 4-8 technical replicates (n=4-8 wells). One-way ANOVA with Dunn’s multiple comparisons to test for significance, mean ± SD, ***p<0.001, *p<0.05.

Glutamate reuptake is another key homeostatic function of astrocytes and is considered a hallmark for the characterization of differentiated astrocytes *in vitro*^34^. To test whether our two astrocyte subtypes differed in their glutamate reuptake ability, we employed a fluorometric glutamate uptake assay that measures the depletion of glutamate from the culture medium over time, starting from a known concentration (100 μM) (**Figure 3E**). We used a selective glutamate transport inhibitor L-trans-pyrrolidine-2,4-dicarboxylate (PDC)^35^ as a negative control. We observed that, while both BMP4 and CNTF astrocytes are capable of glutamate uptake, the rate is twice as high for CNTF astrocytes than for astrocytes differentiated with BMP4 (c.a. 33 μM/hr and 15μM/hr, respectively) (**Figure 3E**). This difference also corresponds to differences in protein levels of the glutamate transporter excitatory amino acid transporter 1 (EAAT1), which are significantly higher in CNTF astrocytes (**Figure 3F**).

### BMP and CNTF differentiated astrocyte morphologies are not altered by neuron contact

Astrocytes have a bidirectional relationship with neurons *in vivo*, which is known to influence the morphology, homeostasis, and function of both cell types^36,37^. For this reason, we investigated the morphologies and functional properties of both BMP4 and CNTF astrocytes when co-cultured with neurons. First, we wanted to test whether their morphology would be altered by the presence of neurons, using our established imaging pipeline with GFP^+^ BMP4 and CNTF astrocytes. When co-cultured with neurons^38^, CNTF and BMP4 astrocytes displayed subtle morphological differences (not reflected in previously defined parameters) but retained their distinct base cell shapes (**Figure S3C, Figure 3G**). Importantly, co-culture with neurons significantly upregulated excitatory amino acid transporter 2 (EAAT2) levels in both BMP- and CNTF-differentiated astrocytes compared to monoculture (**Figure S3D**), suggesting that both these distinct subtypes can respond productively to neurons.

Astrocytes also have a key role in promoting neuronal synapse formation^39,40^. We therefore tested whether BMP4 and CNTF astrocytes differ in their ability to stimulate synaptogenesis when co-cultured with neurons. To do this we differentiated cortical neuron progenitors^38^, either alone or in the presence of BMP4 or CNTF differentiated astrocytes. To assess the co-localization of synapses, we stained for re- and post-synaptic proteins synaptophysin (SYN) and post-synaptic density-95 (PSD95), along with MAP2 to demarcate dendrites (**Figure S3E**), and quantified the number of SYN^+^/PSD95^+^ puncta per 100μm^2^ of MAP2 dendrites. Our results indicate that BMP4 and CNTF astrocytes both significantly promote synaptogenesis to a similar extent within our system (**Figure 3H**).

### BMP4 and CNTF astrocytes constitute transcriptionally distinct subtypes

Having established that BMP4 and CNTF astrocytes are functionally and morphologically distinct subtypes, we investigated their transcriptomic profiles to elucidate the molecular basis of these differences.

We then performed bulk RNA-sequencing (RNA-seq) on astrocyte progenitors under basal conditions, as well as BMP4- and CNTF astrocytes differentiated for 2 and 3-weeks (**Figure 4A**) to compare their transcriptional profiles. Our analysis revealed minimal differences between 2- and 3-weeks differentiated astrocytes, for both BMP4 and CNTF (**Figure S4**). Therefore, we focused our analysis on 3-week differentiated astrocytes (**Figure 4A**). Principal component analysis (PCA) highlighted that both astrocytes subtypes are transcriptionally distinct from progenitors and from each other (**Figure 4B**), with 2162 differentially expressed genes (DEGs) between BMP4 and CNTF astrocytes (**Figure 4C**). Importantly, comparing these DEGs, we were able to confirm at the gene expression level (**Figure S5**) what we previously observed at the protein level (**Figure 3D and 3F**) for *Slc1a3* (EAAT1) and *Cx43* (Connexin-43). When we compared the differentially expressed genes (DEGs) between progenitors and each subtype, we found that around a half of all DEGs are shared between both differentiation protocols, while the other half is uniquely altered between astrocytes differentiated with either BMP4 or CNTF cytokines (**Figure 4D**). This suggests that, while the process of differentiation from progenitors involves common mechanisms, BMP4 and CNTF astrocytes also undergo unique changes that contribute to forming their distinct identities and, potentially, their divergent morphologies.

**Figure 4-.**
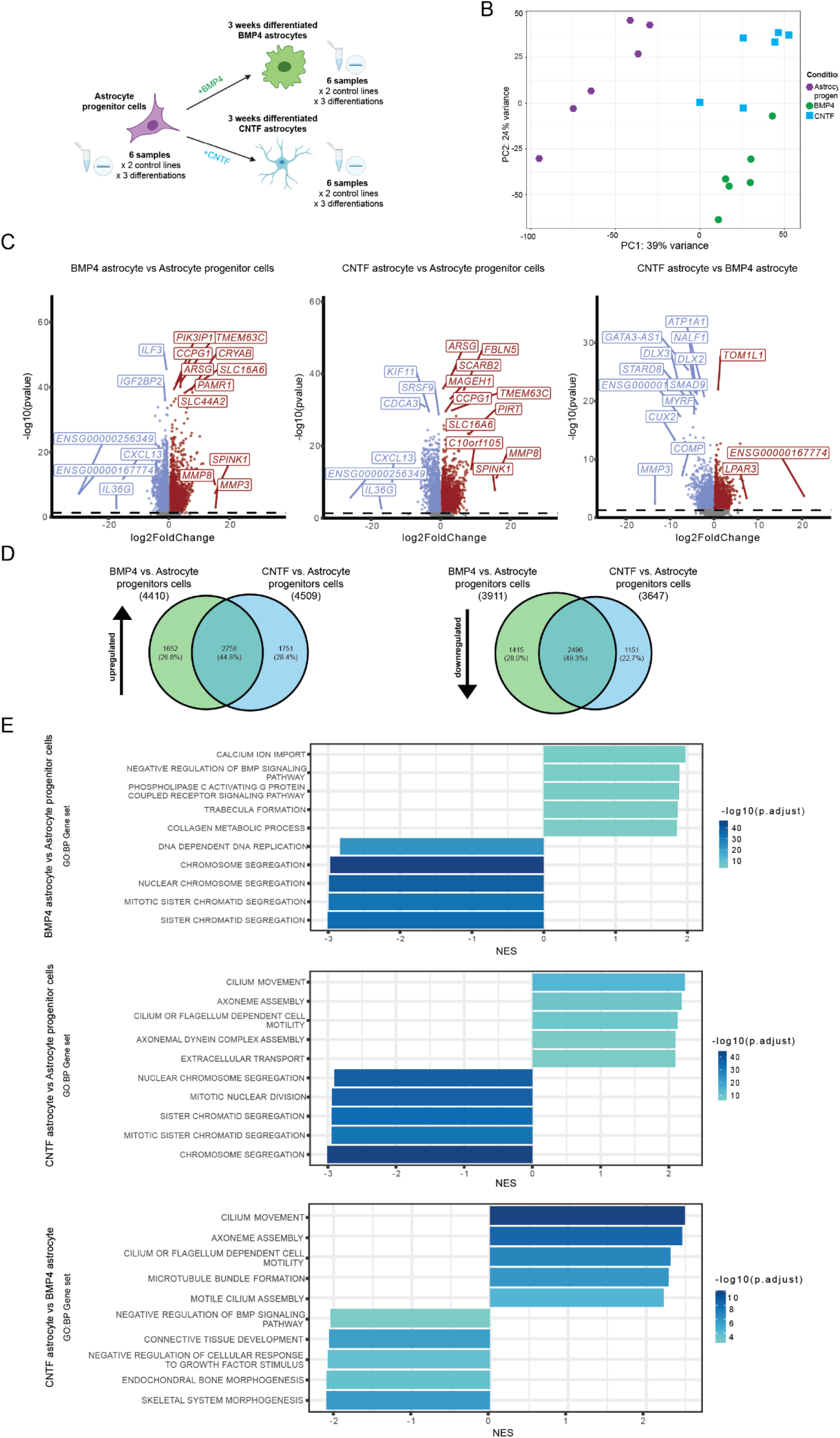
Transcriptional profile differences between BMP4 and CNTF astrocyte subtypes. A) Schematic of the experimental protocol for bulk RNA sequencing of astrocyte progenitors and 3-week differentiated BMP4 and CNTF astrocytes. For each timepoint, 6 samples were analysed, coming from 2 control lines with 3 independent differentiations performed for each line. B) Principal component analysis of transcripts from astrocyte progenitors and BMP4 and CNTF differentiated astrocytes. C) Pairwise comparison volcano plots with top differentially downregulated (blue) and upregulated (red) genes (adjusted p-value<0.05) for BMP4 astrocytes versus Astrocyte progenitors, CNTF astrocytes vs. Astrocyte progenitors, and CNTF astrocytes versus BMP4 astrocytes. The significance (−log_10_ p-value) is plotted on the Y axis while the fold change (log_2_ fold change) is plotted on the X axis. D) Venn diagrams showing the percentages of common and subtype-specific genes that are upregulated or downregulated during the transition between progenitors and differentiated BMP4 and CNTF astrocytes. E) Bar plot showing the top 5 gene sets upregulated and downregulated in BMP4 astrocytes versus astrocyte progenitors, CNTF astrocytes versus astrocyte progenitors, and CNTF astrocytes versus BMP4 astrocytes. Gene sets were derived from the Biological Process (BP) ontology. Gene sets are ordered according to the absolute effect size (NES) and coloured by significance (−log_10_ p-value adjusted).

To determine the biological pathways underlying the specialization of BMP4 and CNTF astrocytes, we performed gene set enrichment analysis (**Figure 4E and S6-S11**).

As expected, compared with astrocyte progenitors, in both BMP4- and CNTF-differentiated astrocytes the main downregulated gene sets are related to cell division and DNA replication, which suggests that both differentiation processes give rise to post-mitotic cells (**Figure 4E**).

Comparing the two differentiated subtypes, we observed that gene sets with the highest normalized enrichment score (NES) for CNTF astrocytes compared to BMP4 astrocytes were overwhelmingly associated with the microtubule cytoskeleton. On the other hand, gene sets enriched in BMP4 differentiated astrocytes were related to development and morphogenesis, which is in accordance with the functions carried out by BMPs in these processes.

Next, we wanted to compare the transcriptional profile of BMP4- and CNTF-differentiated astrocytes generated using our *in vitro* system with recently published single cell RNA profiling datasets of different cell types isolated directly from the human brain^41,42^ and found that both show a strong correlation with human astrocytes (Pearson’s correlation coefficient *r*=0.5 and 0.61 for BMP4- and CNTF-differentiated astrocyte DEGs with adjusted p-value<0.05, respectively) (**Figure S12A-B and S13A-B**). In addition, the comparison with a recently generated dataset comprising single nuclei RNA-seq data from white and grey matter astrocytes^43^ revealed that both subtypes show a more significant correlation with white matter astrocytes, with CNTF astrocytes displaying higher correlation to some of the specific white matter astrocyte clusters identified, namely ciliated astrocytes (**Figure S14A-C**).

### Effect of BMP4 and CNTF on primary isolated mouse astrocytes

After analysing the differential gene expression profiles of our human astrocytes differentiated with BMP4 and CNTF, we evaluated whether these growth factors have similar effects on primary rodent astrocytes. We isolated astrocytes from the neocortex of postnatal day (P) 2-3 mouse pups according to a previously described protocol^44^. After early-stage differentiation, immature astrocytes were treated for 7 days with BMP4 or CNTF. We used the previously validated astrocyte trophic factor heparin-binding EGF-like growth factor (HBEGF)^44,45^, or left cells untreated as controls (**Figure 5A**).

**Figure 5-.**
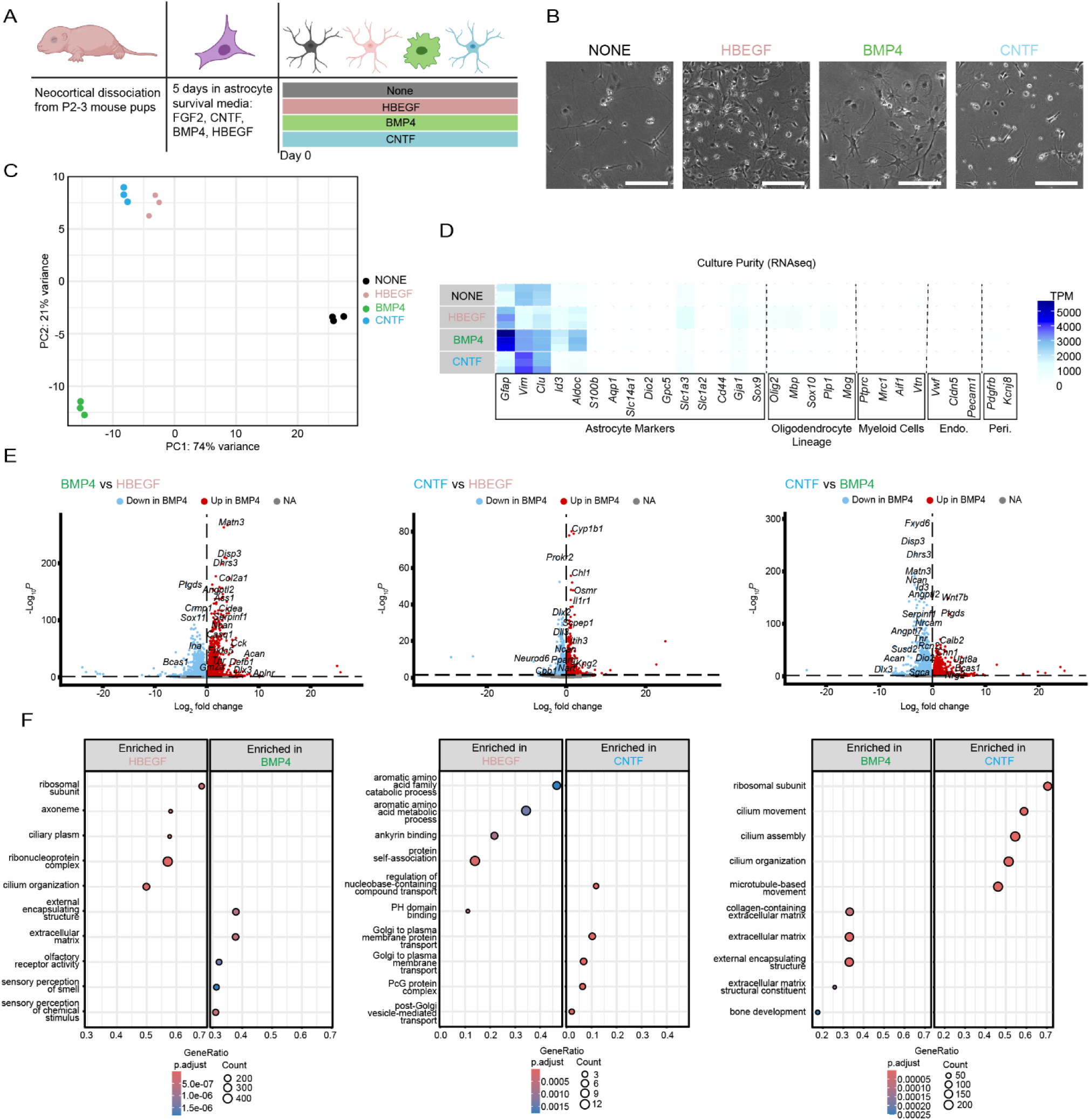
Effects of BMP4 and CNTF on mouse primary astrocytes. A) Schematic for generation of mouse astrocytes. B) Representative brightfield images of astrocytes treated for each condition at 7 days; scale bar is 20 µm. C) Dimensionality reduction by principal components 1 and 2 of sequenced samples. Each dot represents an individual technical replicate and experimental conditions are grouped by color (n=3 technical replicates/condition). D) Heatmap of transcript per million (TPM) expression of various CNS cell type marker genes across experimental conditions. Known cell type markers are grouped along X axis by cell type (n=3 technical replicates/condition). Rows are representative of individual technical replicates. E) Volcano plots of differentially expressed genes between respective treatment conditions. Positive log_2_ fold change is representative of enrichment while negative log_2_ fold change is representative of de-enrichment. F) Dot plotting of GO biological process gene set enrichment. Gene ratio describes the ratio of significantly changed genes to the total genes in each pathway set. Dot size represents the total number of enriched or de-enriched genes while color is representative of adjusted p-value.

Brightfield imaging on Day 7 of treatment revealed similar patterns of morphology to human astrocytes (**Figure 5B**). We found that BMP4-treated mouse astrocytes often had rounder morphologies, with shorter processes extending from a large soma. CNTF-treated astrocytes adopted less round morphologies with longer processes extending from relatively smaller somas. We found that there was more heterogeneity within these mouse astrocytes than the human astrocytes, likely owing to the fact that astrocytes obtained by acute isolation represent a more heterogenous population at baseline than would stem cell derived astrocytes.

We next performed bulk RNA-seq to evaluate transcriptomic changes between BMP4 and CNTF-treated astrocytes. PCA analysis revealed that both BMP4 and CNTF-treated astrocytes were transcriptionally distinct from one another and from untreated astrocytes (**Figure 5C**). Interestingly, CNTF and HBEGF-treated astrocytes were transcriptionally similar. Before performing differential expression analysis, we confirmed the purity of our cultures by evaluating robust expression of various astrocytic marker genes, as well as low-to-no expression of marker genes for other potential contaminating cell types (**Figure 5D**). Importantly, even untreated mouse astrocytes were highly pure, suggesting that they are already committed to an astrocyte fate after purification which parallels human astrocyte progenitors.

Differential gene expression analysis detailed transcriptomic differences between BMP4 and CNTF-treated astrocytes (**Figure 5E**). We find found 2649 DEGs between BMP4 and CNTF-treated astrocytes (log2FoldChange < −0.6; p-value < 0.05). To investigate similarities from between human to mouse astrocytes, we looked at SLC1A3lc1a3 Slc1a3 and GjaJAja1 expression levels across conditions (**Figure S15**). We saw higher expression of SLC1A3lc1a3 Slc1a3 in CNTF-treated astrocytes than BMP4-treated astrocytes, which corroborates the functional glutamate uptake assay, immunostaining and sequencing results in human astrocytes (**Figure 3E-F and S5**). Gja1 expression was similar across treatments, also corroborating the immunostaining and sequencing of human astrocytes (**Figure 3D and Figure S5**).

Finally, to investigate the biological pathways being differentially regulated by BMP4 and CNTF, we performed GO biological pathway gene set enrichment analysis (**Figure 5F**). Like human astrocytes, mouse astrocytes treated with CNTF were enriched for pathways involving microtubule and cilium movement. On the other hand, BMP4-treated mouse astrocytes were enriched for pathways involving the extracellular matrix, possibly pointing to a role for BMP4 astrocytes in establishing the extracellular milieu. In summary, these data show that many of the upregulated gene and pathways that respond to developmental BMP4 and CNTF treatment in mouse astrocytes areis evolutionarily conserved in human. Morphologies between human and mouse astrocytes were similar, and CNTF-treated astrocytes in both humans and mice upregulate transcripts associated with cilium and microtubule movement, which is related to their relatively more process-bearing morphology. in both human and mouse astrocytes.

### The morphology of BMP4 and CNTF astrocytes relies on distinct cytoskeletal components

Based on the transcriptomic analysis results and our previous observations on their morphology, we hypothesised that a different cytoskeletal organization might underpin the distinct phenotypes of our astrocyte subtypes. To test this, we first investigated the role of the actin cytoskeleton in determining astrocyte morphology in our system. We exposed GFP^+^ BMP4- and CNTF-differentiated astrocytes for 2 hours to either latrunculin B^46^ (LAT B; 1 μM) or Rho-associated, coiled-coil containing protein kinase (ROCK) inhibitor Y27632 dihydrochloride^47^ (Y27632; 25 μM), which both interfere directly or indirectly with actin polymerization and contractility, and performed single-cell morphology profiling using our semi-automated pipeline.

We found that BMP4 astrocytes treated with Y27632 displayed a more ramified morphology, with processes emanating from the cell body (**Figure 6A**), although the overall size remained unchanged, while DMSO-treated controls remained unaffected (**Figure S16A**). Conversely, treatment with LAT B caused a decrease in area, as well as the acquisition of a more irregular and ramified shape (**Figure 6A**). The same effects were also evident in the quantification of the key shape descriptors (**Figure 6B and S16B**). Interestingly, CNTF astrocytes treated with either drug remained mostly unchanged, suggesting that actin disruption does not affect their overall architecture (**Figure 6A-B and S16B**). The same effects observed on cultures sparsely transfected with GFP were also verified at the population level, by immunostaining for the tubulin cytoskeleton, which is presumed to be unaffected by these interventions (**Figure S16C**).

**Figure 6-.**
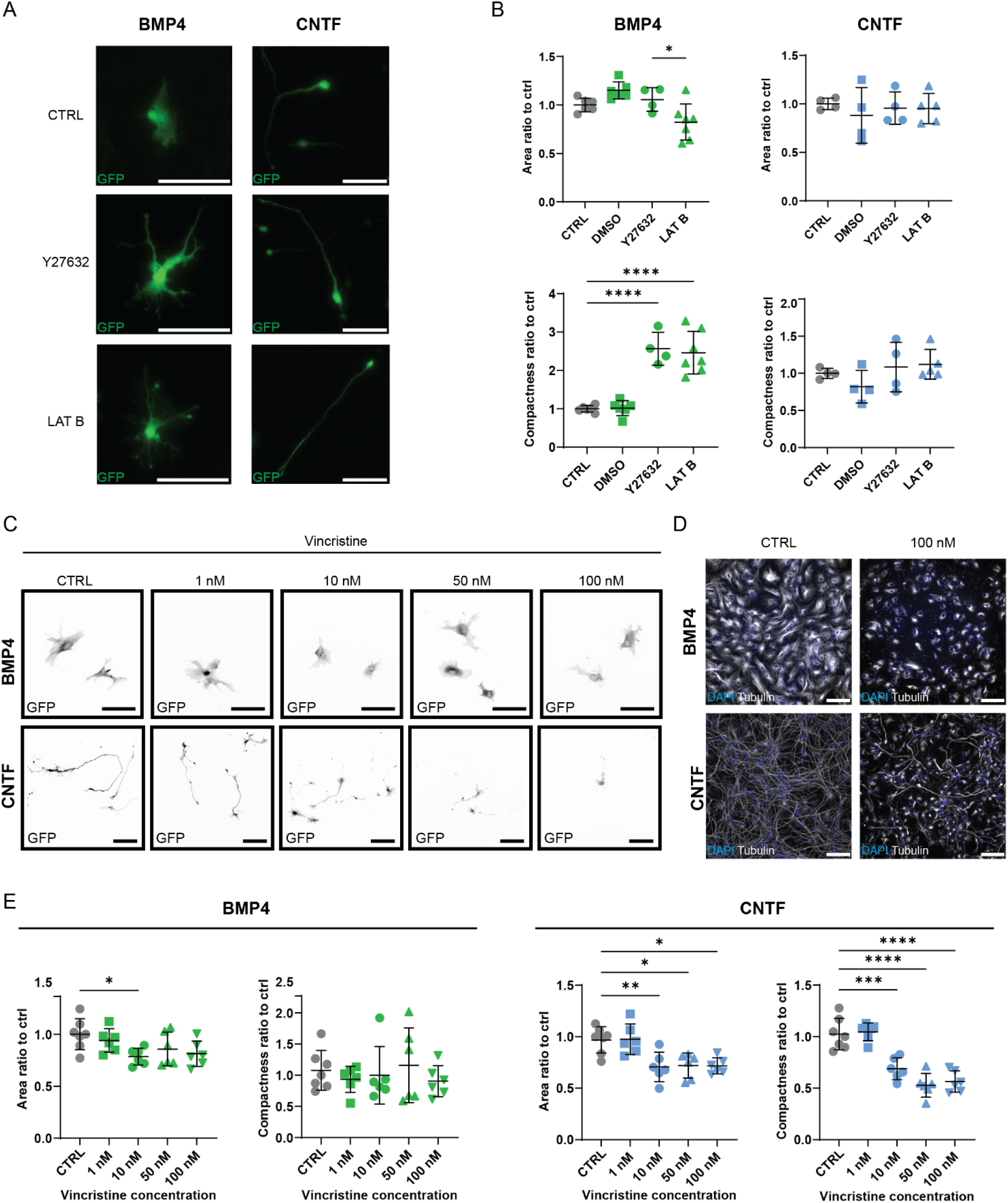
The morphology of BMP4 and CNTF astrocytes relies differently on the actin and microtubule cytoskeleton. A-B) Actin inhibition significantly affects the morphology of BMP4 astrocytes but not that of CNTF astrocytes. A) Representative images of GFP^+^ BMP4 and CNTF astrocytes, either untreated (CTRL) or treated with actin inhibitors Y27632 dihydrochloride (Y27632; 25 μM) or latrunculin B (LAT B; 1 μM) for 2 hours before live imaging. Scale bar: 100 μm. B) Quantification of shape changes in the presence of actin inhibitors analysing area and compactness expressed as a ratio compared to control (ctrl, black). Data represents well averages of technical replicates of 2 control lines from 2 experimental blocks (n=4-5 wells total), mean ± SD. Ordinary one-way ANOVAs with Tukey’s post-hoc was used to test for significance. *p<0.05, **p<0.01, ****p<0.0001 ns or no comparisons shown: non-significant. C-E) Microtubule destabilisation significantly affects the morphology of CNTF astrocytes but not that of BMP4 astrocytes. C) Representative images of GFP^+^ BMP4 and CNTF astrocytes either left untreated (CTRL) or treated with increasing concentrations of vincristine (1, 10, 50 and 100 nM). Scale bar: 100 μm. D) Representative images of BMP4 and CNTF astrocytes, either untreated (ctrl) or treated with 100 nM vincristine, stained with nuclear (DAPI, blue) and microtubule cytoskeletal markers (Tubulin, white). Scale bar: 100 μm. E) Quantification of shape changes in the presence of vincristine analysing area and compactness, expressed as a ratio compared to control (ctrl, black). Data represents well averages of technical replicates of 2 control lines from 2 experimental blocks (n=6-7 wells total), mean ± SD. Ordinary one-way ANOVAs with Tukey’s post-hoc was used to test for significance.

To assess potential differences between the reliance of BMP4 and CNTF astrocyte morphological features on microtubular cytoskeletal components, we exposed differentiated astrocytes to increasing concentrations of the microtubule destabilising drug vincristine^48^ (VIN; 1, 10, 50 and 100 nM). VIN was effective in altering the microtubular architecture for both BMP4 and CNTF astrocyte subtypes (**Figure 6D**), however, only CNTF astrocytes showed a radical alteration in their overall architecture. BMP4 cells were largely unchanged (**Figure 6C top**), with only a decrease in area with concentrations of VIN above 10 nM (**Figure 6E**). As the effect size of this is quite small and none of the other shape descriptors displayed significant differences, we concluded that the overall architecture of BMP4 astrocytes is not overly reliant on the microtubule network, but rather on actin contractility (**Figure 5E and S16D**).

Conversely, we found that in CNTF astrocytes VIN causes a concentration-dependent shortening of the cell processes, which constitute the main morphological hallmark of these cells (**Figure 6C, bottom**). This was then validated by the quantification of morphometric parameters which showed that these cells display a reduction in area, compactness, and perimeter at all VIN concentrations except for 1 nM, while the median radius was only altered in the 50 nM condition (**Figure 6E and S16D**).

Our combined investigation of cytoskeletal components, supported by the RNA-seq data and morphological analysis, indicates that BMP4 astrocytes rely mostly on actin polymerization and contractility for their shape, while CNTF astrocytes likely owe their morphology to a stable polymerized microtubule network.

### The distinct morphology of BMP4 and CNTF astrocytes correlates with different distribution and motility features of their mitochondria

While astrocytes use glycolysis for most energy production^49^, they also heavily rely on mitochondria for calcium storage and signaling, as well as fatty acid and neurotransmitter metabolism^50–52^. Astrocyte mitochondria are located in cytoplasmic microdomains throughout the cell, including in the finer processes^53^, in order to meet local energy demands and upkeep ionic balance^56^.

While the propagation and modulation of astrocytic ICWs is mainly mediated by the endoplasmic reticulum (ER)^54^, mitochondria also actively participate in local Ca^2+^ dynamics both by releasing and sequestering Ca^2+^ ^52^. Given the observed differences in ICW dynamics between the astrocyte subtypes, and the fact that mitochondrial transport is microtubule-bound^55^, we reasoned that the observed differences between the cytoskeletal organization of BMP4 and CNTF astrocytes may indicate a potential difference in distribution and motility of mitochondria between these cells.

To evaluate mitochondrial distribution within single cells, we transfected BMP4 and CNTF astrocytes with both a cytoplasmic RFP and a mitochondrial targeted fluorophore (mito-dendra2)^56^ before fixing them and staining the nuclei with DAPI. We then segmented each cell into 25 μm bins automatically marked radially around the nuclei (**Figure 7A**) and quantified the proportion of fluorescence intensity as a proxy of mitochondrial distribution in the different compartments (**Figure 7B**). While BMP4 cells maintain over 60% of their mitochondria in the perinuclear space (i.e. 0-25 μm from nucleus edge), in CNTF astrocytes less than half of the mitochondria are localized within this perinuclear space. Instead, there seems to be numerous hotspots of Mitodendra2 signal distributed along the cell processes (**Figure 7B&C**). Additionally, BMP4 cells overall show a higher number of mitochondria occupying their cytoplasm per unit area, which may be due to a difference in either content or size of the mitochondria between the two subtypes (**Figure 7D**).

**Figure 7-.**
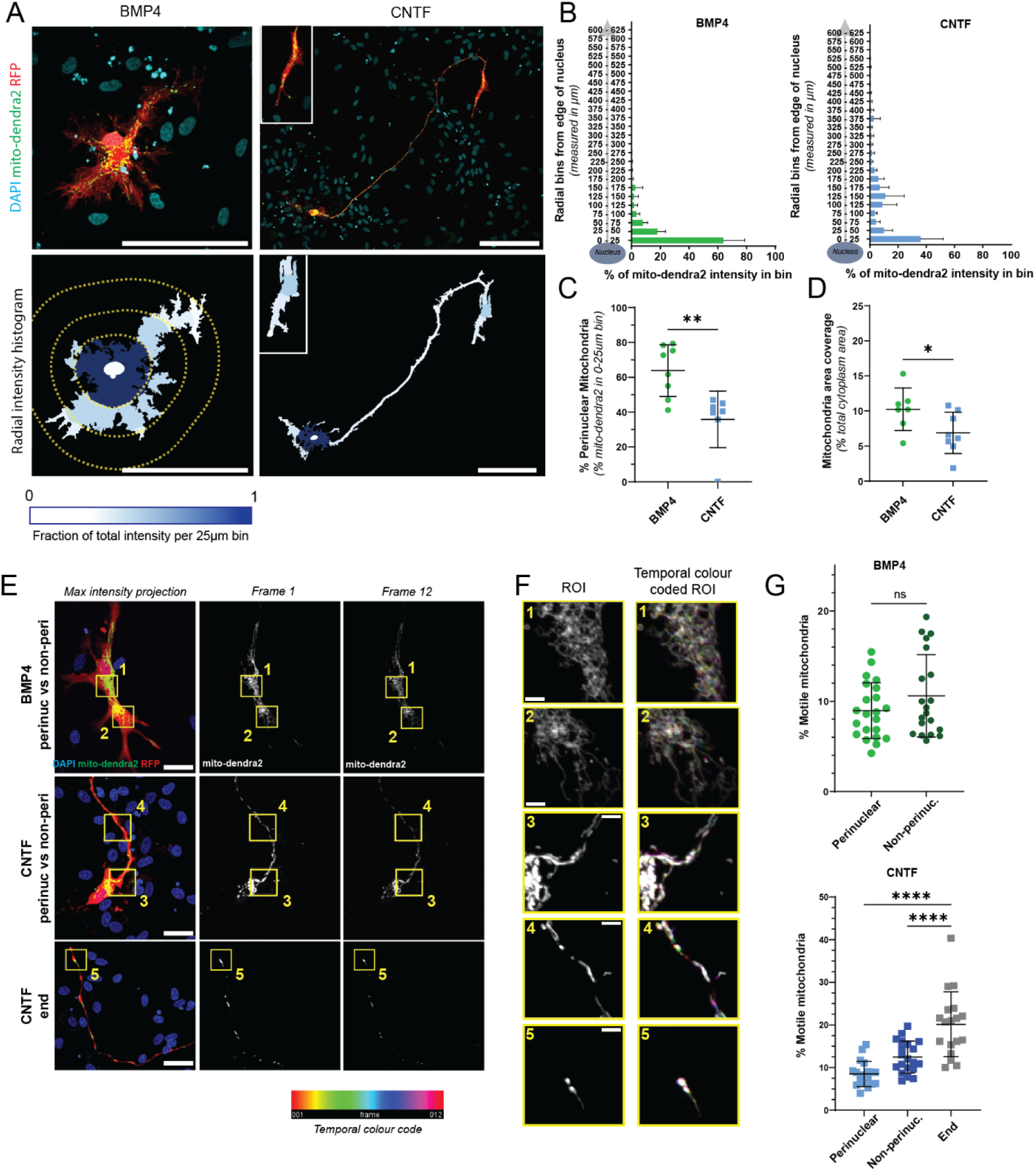
Mitochondrial distribution and motility differ between BMP4 and CNTF astrocytes. A) BMP4 and CNTF 2-weeks differentiated astrocytes were co-transfected with cytoplasmic RFP (red) and mito-dendra2 (green), fixed and immunostained with a nuclear marker (DAPI, blue). The co-transfection shows the distribution of mitochondria within single cells, highlighting the hotspots of mitochondria throughout the processes (zoom inset, CNTF astrocyte). Quantification was performed by measuring mito-dendra2 intensity in 25 μm radial bins around the nucleus (marked with DAPI) within whole single cells, segmented using RFP. The fractional contribution of mito-dendra2 intensity within each bin throughout the entire cell is coloured coded in a gradient from white (0) to dark blue (1). B) Quantification of the fraction of mito-dendra2 intensity per 25 μm radial bin around the nucleus expressed as a percentage in both BMP4 and CNTF astrocytes. C) Quantification of the fraction of mito-dendra2 signal contained within the first radial bin (0-25 μm) as a percentage of the total for BMP4 astrocytes (BMP4 perinuc.) and CNTF astrocytes (CNTF perinuc.). D) Quantification of mitochondria (mito-dendra2) area coverage expressed as percentage of the whole cytoplasm area (RFP) in BMP4 and CNTF astrocytes. Data represents well averages of technical replicates of 2 control lines from 2 experimental blocks (n=6-8 wells), mean ± SD. Unpaired t-tests were used to test for significance. **p<0.01, *p<0.05. E) BMP4 and CNTF astrocytes were co-transfected with free-floating RFP (red) and mito-dendra2 (green) and live dyed with Hoescht 33342 (blue). Maximum intensity projections show regions of BMP4 and CNTF astrocytes, with representative time lapse images of beginning (frame 1) and end (frame 12) of videos for mitochondrial motility analysis over 2 minutes of imaging (mito-dendra2, white). Scale bar: 100 μm. Yellow zoom insets 1 and 2 show perinuclear and non-perinuclear regions, whilst yellow zoom insets 3 for CNTF astrocytes shows the end of the process used as regions of interest (ROIs) for analysis (25 μm x 25 μm ROIs). F) Temporal colour coded ROIs show stationary mitochondria as white, whilst coloured pixels indicate the frame in which there was the most recent change in motility. Scale bar: 5 μm. G) Quantification of the percentage of motile mitochondria within each subcellular compartment for BMP4 astrocytes (perinuclear vs non-perinuclear) and CNTF astrocytes (perinuclear vs non-perinuclear vs end of process). Data represents averages of percent change from each slice for each cell, of 2 control lines from 2 experimental blocks (n=18-23 cells from 7-8 wells), mean ± SD. Mann-Whitney U test was used to test for significance for BMP4 cells and ordinary one-way ANOVA with Tukey’s post-hoc used to test for significance for CNTF cells. ****p<0.0001, ns: non-significant.

We next sought to test whether the distribution of mitochondria in the two subtypes could be linked to their motility within the astrocytes. To do this, we performed live imaging on BMP4 and CNTF astrocytes co-transfected with RFP and mito-dendra2 and stained with Hoechst 33342 to visualize the nucleus (**Figure 7E-F**). Mitochondrial movements were recorded over 2 minutes at 0.1 frame/sec across 25 μm^2^ regions of interest (ROIs) drawn either in the perinuclear region (0-25 μm from the nucleus, **Figure 7E** inset 1&3), in a non-perinuclear area (25-100 μm from the nucleus, **Figure 7E**, inset 2&4), or in the case of CNTF astrocytes, in an area at the end of a process (0-50 μm from the process end, **Figure 7E** inset 5). Analysis of these ROIs across different cells and replicated revealed that, while BMP4 astrocytes show no difference in mitochondrial motility between the perinuclear and non-perinuclear regions, CNTF astrocytes show a two-fold increase in the fraction of motile mitochondria at the end of the processes compared to either their perinuclear or non-perinuclear areas (**Figure 7G**).

Taken together, these results show that distribution and motility of mitochondria differs substantially between the two subtypes, potentially indicating a distinct organisation of the subcellular domains in CNTF astrocytes. In particular, the higher presence of mitochondrial hotspots in the periphery of CNTF astrocytes matches our findings regarding their dependence on a stable microtubule network (**Figure 5D-E**) and it may hint towards one of the mechanisms behind the faster ICW dynamics in these cells (**Figure 3A-B**).

### BMP4 and CNTF astrocytes have distinct transcriptional and functional reactive phenotypes

Astrocytes respond to a wide range of pathophysiological stimuli, inflammation, and injury by undergoing a profound transformation in their morphological, functional and molecular profile which is commonly referred to as ‘reactivity’^57^. As astrocyte reactivity can generate a wide range of phenotypic outcomes^58^, we were interested in understanding whether it would differentially impact the two human astrocyte subtypes. We therefore treated 2-week differentiated BMP4 and CNTF astrocytes for an additional week in the presence or absence of an established cocktail of factors known to induce astrocyte reactivity comprising of TNF, IL-1α, and C1q (TIC)^59^, and performed bulk RNA-seq comparing treated and untreated astrocytes (**Figure 8A**). We found that, in both cases, reactivity induced a significant transcriptomic transformation, with the upregulation of known pro-inflammatory genes (e.g. *IL36*, *CXCL13*, and *CSF3*) in the treated astrocytes compared to their untreated counterpart (**Figure 8B-C**),

**Figure 8-.**
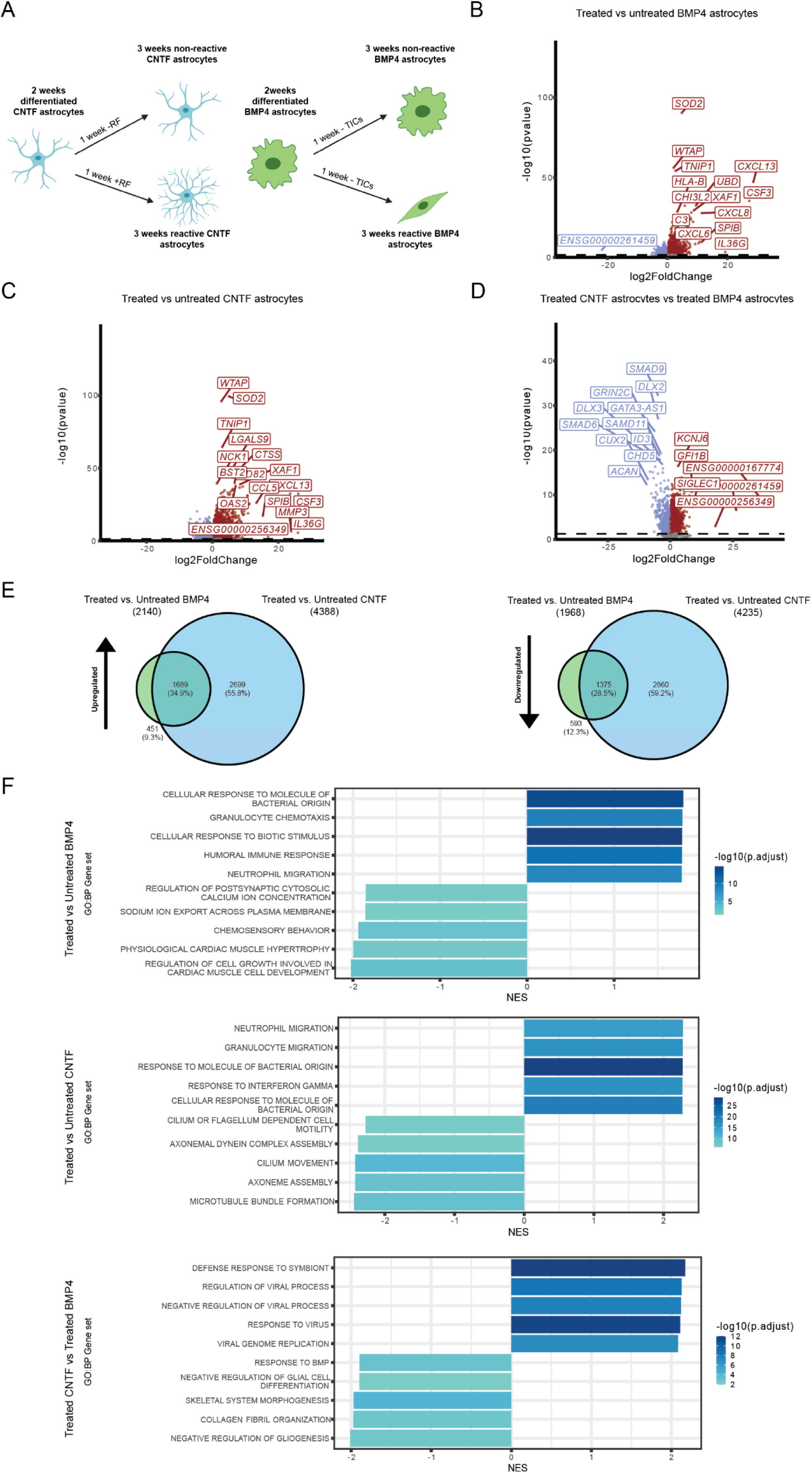
The induction of reactivity causes distinct transcriptomic changes in BMP4 and CNTF astrocytes. A) Schematic of the experimental protocol for bulk RNA-sequencing of astrocytes differentiated for three weeks with BMP4 or CNTF and with or without treatment with known reactivity factors (TNF, IL-1α, C1q; TIC) during the third week of culture. For each timepoint, six samples were analysed, coming from two control lines with three independent differentiations performed for each line. B-D) Pairwise comparison volcano plots for treated BMP4 astrocytes vs. untreated BMP4 astrocytes (B), treated CNTF astrocytes vs. untreated CNTF astrocytes (C), and treated CNTF astrocytes versus treated BMP4 astrocytes (D). The significance (−log_10_ p-value) is plotted on the Y axis while the fold change (log_2_ fold change) is plotted on the X axis. Downregulated genes (log_2_FoldChange<-0.6; p-value<0.05) are coloured in blue, while upregulated genes (log_2_FoldChange>0.6; p-value<0.05) are coloured in red. E) Venn diagrams showing the percentages of common and subtype-specific genes that are upregulated or downregulated in response to the induction of reactivity in BMP4 and CNTF astrocytes. F) Bar plot showing the top five gene sets upregulated and downregulated in treated BMP4 astrocytes vs. untreated BMP4 astrocytes, treated CNTF astrocytes vs. untreated CNTF astrocytes, and treated CNTF astrocytes vs. treated BMP4 astrocytes. Gene sets were derived from the Biological Process (BP) ontology. Gene sets are ordered according to the absolute effect size (NES) and coloured by significance (−log_10_ p-value adjusted).

However, when directly comparing the expression profiles of reactive BMP and CTNF astrocytes (**Figure 8D-E**), we observed that, in addition to a significant number of these common pan-reactive markers, there were also many DEGs that were unique to either astrocyte reactive subtype (**Figure 8E**). For example, we noted differences in the expression of genes such as *MMP3* (upregulated in CNTF, but not in BMP4 reactive astrocytes) and *ACAN* (upregulated in BMP4, but not in CNTF reactive astrocytes), among others (**Figure 8D**). Interestingly, there was a higher number of CNTF-specific DEGs (55.8% of upregulated genes and 59.2% of downregulated genes), while common DEGs represent 34.9% and 28.5% of up- and downregulated genes, respectively. This indicates that the majority of genes that change their expression during reactivity are CNTF-specific, followed by a lower percentage of genes that are shared among both substates, with the lowest percentage constituted of BMP4-specific DEGs (**Figure 8E**).

To better understand which biological pathways might be involved in the respective reactive transition of BMP4 and CNTF astrocytes, we also performed gene set enrichment analysis (**Figure 8F and Figure S10A-C**). This analysis indicated that, as expected, the majority of gene sets upregulated in both cell types upon exposure to TIC is related to immune regulation and, in particular, the modulation of immune cell migration.

When focusing on the comparison between reactive CNTF and reactive BMP4 astrocytes, we observed that gene sets specifically enriched in CNTF reactive astrocytes seem to be associated with antiviral responses which may point towards a specialized function of CNTF astrocytes in this context. On the other hand, gene sets enriched in reactive BMP4 astrocytes compared to reactive CNTF astrocytes were linked with morphogenesis, cell signalling and extracellular matrix (ECM) production. We also observed that, when transitioning to a reactive state, CNTF astrocytes seem to downregulate the same gene sets linked to the tubulin cytoskeleton which were driving their differentiation from progenitors, suggesting a potential morphological change.

Overall, the transcriptomic analysis suggests that BMP4 and CNTF astrocytes, beyond the already shown functional differences in homeostatic conditions, might also acquire distinct reactive states. We therefore wanted to verify whether the observed divergence in gene expression translated into unique morphological and functional reactive profiles.

As the baseline morphology of these two subtypes of astrocytes is very different, we first wanted to test whether the induction of reactivity would make them diverge further or converge to one common “reactive shape”.

Interestingly, when we performed the same morphological characterisation from **Figure 1** with TIC versus control treated GFP^+^ BMP4 and CNTF astrocytes, we found that both subtypes presented morphological alterations. However, while BMP4 astrocytes maintained the same base morphology (i.e. perimeter and extent did not vary significantly at the single-cell level), CNTF reactive astrocytes showed significantly reduced perimeter and increased extent compared to their non-reactive counterparts, which points towards them having larger and rounder cell bodies with shorter processes (**Figure 9Ai and B**). At the population level, we also confirmed that in both subtypes the reactive transformation leads to a significant rearrangement of the actin cytoskeleton (**Figure 9A ii**). This is particularly evident in BMP4 cultures which, when treated, seem to orient in patches around the culture and adopt a “palisading” phenotype^60^.

**Figure 9-.**
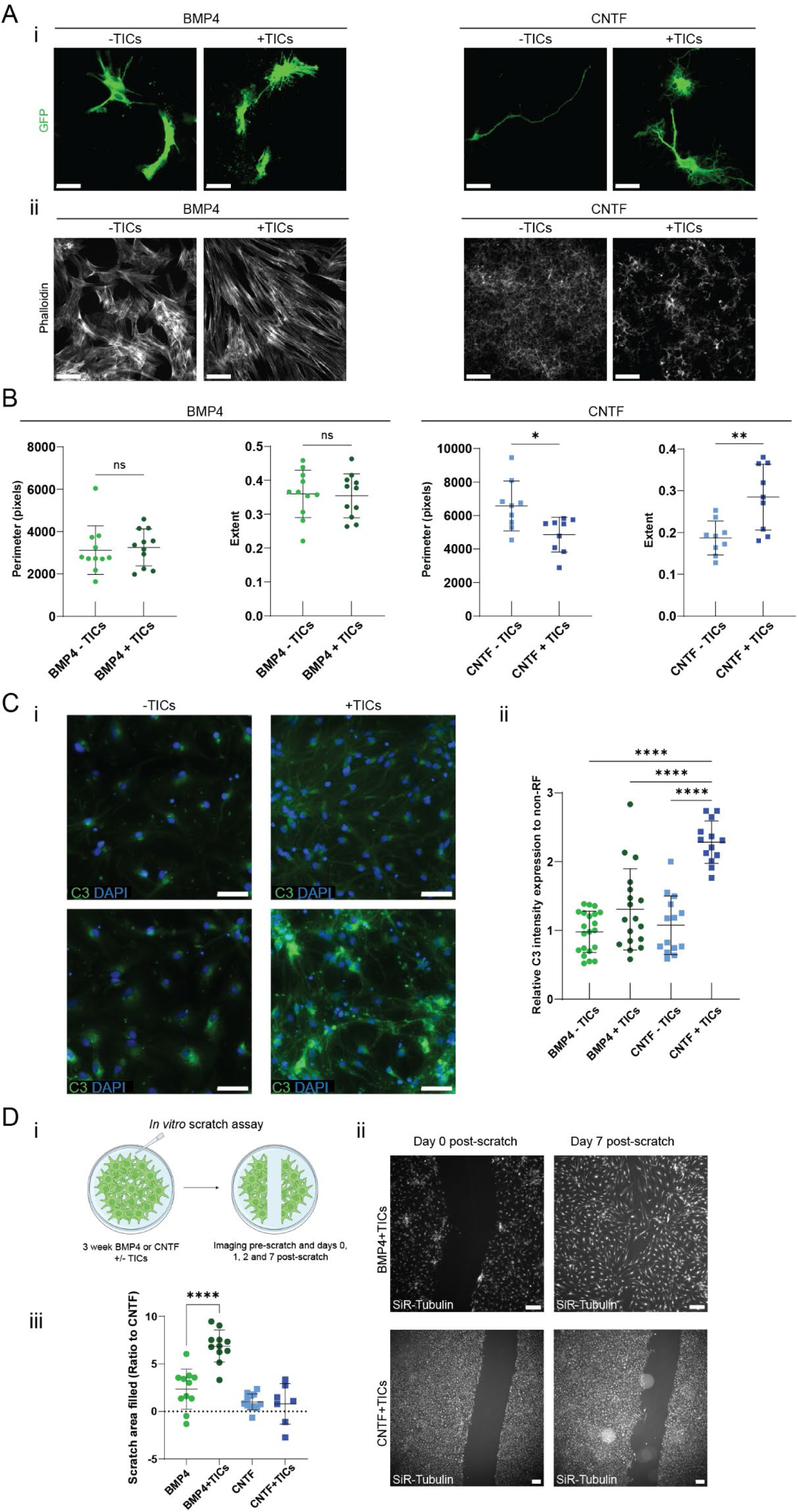
BMP4 and CNTF acquire distinct reactive phenotypes when treated with inflammatory stimuli. A) Representative images of BMP4 and CNTF astrocytes cultured for an additional week in the absence (−TIC) or presence (+TIC) of reactivity inducing factors (TIC; TNF, IL-1α, C1q). GFP^+^ astrocytes (i) (green) were used to visualise a change in overall cell shape, whilst non-transfected astrocytes were immunostained with phalloidin (ii) to visualize actin organization at the population level. Scale bar: 100 μm. B) Quantification of changes in GFP shape in BMP4 and CNTF astrocytes in the presence or absence of reactivity factors (−TIC and +TIC, respectively). Unpaired t-test to test for significance. Data represents technical replicate well averages of astrocytes from two control lines from two independent experiment blocks (n=9-11 wells) mean ± S. **p<0.01, *p<0.05, ns: non-significant. C) (i) Representative images of BMP4 and CNTF astrocytes cultured for an additional week in the absence (−TIC) or presence (+TIC) of reactivity inducing factors (TIC; TNF, IL-1α, C1q) immunostained with nuclear marker (DAPI) and Complement Component 3 (C3) to assess C3 protein changes upon reactivity induction. Scale bar: 100 μm. (ii) Quantification of C3 intensity per field of view normalised to median of non-reactive (non-TIC) control for each cytokine group. Ordinary one-way ANOVA with Šídák’s multiple comparisons to test for significance. Data represents fields of views from two control lines from two independent experiment blocks (n=13-21 f.o.v.’s) mean ± SD. ****p<0.0001. D) (i) Schematic of experimental protocol for in vitro scratch assay of differentiated astrocytes. (ii) Representative images of the scratch area of reactive BMP4 (BMP4+TIC) and reactive CNTF (CNTF+TIC) immediately after scratch and 7 days after scratch. Scale bar: 200 μm. (iii) Quantification of astrocyte wound healing response expressed as percentage of scratch area filled at day 7 compared to day 0 for both the cytokine groups +/− reactivity factors (TICs) and normalized to CNTF. Ordinary one-way ANOVA with Tukey’s multiple comparisons to test for significance. Data represents well averages from two control lines from two independent experiment blocks (n=8 wells/condition) mean ± SD. ****p<0.0001.

In addition to evaluating morphology, we also sought to investigate protein changes upon reactivity induction in BMP4 and CNTF astrocytes. Complement factor 3 protein (C3) has widely been reported as a marker of TIC-induced astrocytes ^59,61,62^ ^63^. We therefore immunostained for C3 in our reactive and non-reactive BMP4 and CNTF astrocytes (**Figure 9C**). Protein levels revealed significant differences between the two subtypes. Specifically, in BMP4 reactive astrocytes, C3 expression was highly heterogeneous across the culture, while reactive CNTF astrocytes display a significant increase in C3 levels compared to both untreated CNTF astrocytes and BMP4 conditions (**Figure 9C**).

This further validated that BMP4 and CNTF reactive astrocytes might represent different reactive substates, even when stimulated with the same TIC cytokine cocktail, and motivated us to assess their behaviour in an *in vitro* injury model. We performed a live imaging-based scratch assay^64^ (**Figure 9D i**) to assess whether BMP4 and CNTF astrocytes, either under basal conditions or treated with TIC to induce reactivity, would respond differently to the scratch and display differences in their ability to heal an *in vitro* ‘wound’.

We observed that reactive BMP4 astrocytes display a higher ability to fill the damaged area of the culture by Day 7 after scratching compared to any other condition we evaluated, including reactive CNTF astrocytes (**Figure 9D ii-iii**). Specifically, CNTF astrocytes would not close the gap in the time-period we analyzed, and the pre-treatment with TICs did not significantly affect their performance. Conversely, BMP4 astrocytes, which already in basal conditions show a slightly higher area filled post-scratch, display a marked and significant increase in repair capabilities in response to the induction of reactivity (**Figure 9D iii**).

This suggests that treating BMP4 astrocytes with TIC may produce a reactive substate that is more responsive to injury and more migratory than the CNTF counterpart; however, whether this response is protective or deleterious to neurons is a question that will require further investigation.

## Discussion

Morphology has long been recognized as a fundamental property of astrocytes, contributing to their ability to contact their targets and carry out their modulatory functions in a co-ordinated manner. Many recent studies have contributed to mapping the vast heterogeneity of forms and functions that different subtypes of astrocytes acquire in the human brain^65–67^, and the use of human astrocytes *in vitro* for neurobiology and neurodegeneration studies has grown steadily^68,69^. However, morphological diversity has been rarely recapitulated in *in vitro* models, which generally have either one fixed shape, or present a great variability of morphologies across different experiment, lines and laboratories.

Here we describe how to obtain morphologically distinct human astrocyte subtypes *in vitro* from iPSCs with defined differentiation protocols and show how these represent an ideal platform to systematically explore the mechanistic links between astrocyte morphology and their function.

We first show how exposure of human glial restricted progenitors to either CNTF or BMP4 based differentiation media leads to differentiation of astrocyte populations that exhibit all the functional characteristics of mature astrocytes (e.g. glutamate uptake, promotion of synapse formation, intercellular calcium waves, response to reactivity induction) but are fundamentally different in their overall cellular morphology. We developed and employed an unbiased image analysis pipeline to perform a morphological analysis of the salient features that define each subtype, which would also be adaptable in the future for a wider study on different cytokine combinations. Importantly, we show that the morphology acquired is not dependent on medium exposure after initial commitment and cannot be easily switched from one subtype to the other. Moreover, we show that the difference in astrocyte shape correlates with distinct functional profiles and – importantly – is attributable to the reliance of the two subtypes on different elements of the cytoskeleton to acquire and retain their shape.

Our results indicate that the distinct organization and usage of the actin and tubulin cytoskeleton not only directly underlies the morphological difference, but also provides a mechanistic link between the structural and functional profiles of astrocytes. For example, we show that CNTF astrocytes have faster and more frequent calcium waves, which travel through their intricate arborised long processes; this would also imply the need for a more structured way of meeting metabolic demands within those processes, which matches our results showing that mitochondria in CNTF astrocytes are more delocalised and actively transported in the processes. In turn, this also correlates with the reliance of CNTF astrocytes on a stable microtubule cytoskeleton, and the upregulation of microtubule related stabilisers and transporters. Correspondingly, for BMP4 astrocytes, we find the morphological data (i.e. multiple dynamic short processes) correlates with the cytoskeleton analysis (i.e. reliance on actin filaments more than microtubules), which is consistent with both the expression profile and – importantly – the cellular function (e.g. migratory behavior in response to reactivity).

One important aspect of this study that further highlights the need to consider shape-function links in the context of *in vitro* modelling of astrocytes, is the striking difference in the reactivity profile of the two astrocyte subtypes. When exposed to reactivity inducing cytokines^63^, both subtypes strongly upregulated reactivity associated gene families, albeit with subtle differences. Both subtypes also changed their morphology, with CNTF acquiring a more ramified morphology with bigger cell bodies, while reactive BMP4 astrocytes featured a polarised, palisading morphology. It is interesting to note that genes coding for matrix metalloproteinase (MMPs) enzymes appear among the most differentially expressed in both transitions from progenitors to differentiated cells (for example *MMP3* for BMP4, *MMP8* for CNTF), as well as in the transition between untreated and treated CNTF astrocytes (*MMP3*), potentially pointing towards an important role of ECM remodelling in driving this process (**Figure 4C-D and Figure 7C**).

The most striking feature we observed was the opposite behavior these reactive astrocytes have in response to injury, with the reactivity response of BMP- and CNTF-differentiated astrocytes to the well-defined TIC neurotoxicity-inducing cytokines being markedly different.

While our results highlight the links between shape, function and reactivity in human astrocyte, our *in vitro* model is by its nature reductionist and more simplified compared to the *in vivo* setting. On the one hand, this allows us to retain a much higher level of control on each separate component that might influence astrocyte shape and function, but on the other, there are important non-cell autonomous and environmental effects that we cannot fully account for. Firstly, our astrocytes are differentiated and grown in isolation, while *in vivo* astrogenic signals and other cues would be derived from the surrounding neuronal populations^70^. This means that our analysis, except in the case of specific co-culture experiments, does not account for the presence of neurons and other glial cells that are known to modulate the activity of astrocytes^71^, so we cannot account for potential regional and subtype specific developmental cues that might be missing from our culture.

Comparison of our *in vitro* BMP- and CNTF-differentiated astrocyte RNA-seq data with other published human adult brain astrocyte datasets^41,42^ indicated that both astrocyte subtypes exhibited a degree of transcriptional similarity with acutely purified human brain astrocytes. Of note, similarities were additionally observed with vascular smooth muscle cells (SMCs) and fibroblasts for BMP4-astrocytes and ependymal cells for CNTF-astrocytes. This may be due to overlap between pathways activated by BMP4 and CNTF that are also upregulated in these cell types, such as ECM component secretion and modelling by fibroblasts^72^ activated by BMP4, and cilium regulation and movement by ciliated ependymal cells^73^ also activated by CNTF. For this comparison, we used single-cell RNA-seq datasets, which may be an additional caveat in a comparative analysis with the bulk RNA-seq generated in this study – as they provide lower sequencing depth. Taken together, this validates the signature of our *in vitro* generated astrocyte subtypes as *bona fide* human astroglia, while further supporting the idea that each subtype displays unique transcriptional features which are consistent with our morphological and functional analyses.

To further scrutinize the *in vitro* validity of our human astrocyte model, we also investigated the effects of BMP4 and CNTF on primary isolated mouse neocortical astrocytes^44^. Based on morphological and transcriptomic analyses, the growth factors had similar effects on the mouse astrocytes. CNTF drove a similar transcriptomic profile in mouse astrocytes as HBEGF, the gold standard for *in vitro* rodent astrocyte culture, while BMP4-treated mouse astrocytes displayed extensive gene expression differences from both CNTF- and HBEGF-treated astrocytes. When comparing enriched gene sets, we also found similarities between astrocytes differentiated from human and mouse. CNTF-treated astrocytes were enriched for pathways associated with cilium and microtubule movement. BMP4-treated astrocytes were interestingly enriched for ECM regulation, a known function of astrocytes (reviewed in Wiese et al^74^). Taken together, our investigation of rodent astrocytes revealed commonalities in BMP4 and CNTF signaling, further validating our human astrocyte *in vitro* model and the differential effect that CNTF and BMP4 have on astrocytes. It is also important to note that other combinations of cytokines have been shown to have different degrees of potency in driving astrogliogenesis *in vitro*^70^, however, to our knowledge there are no systematic reports on their effect on astrocyte morphology.

Another important caveat of our approach is that most of our experiments are performed in 2D and on ECM-coated tissue culture plastic, therefore our cells lack the tri-dimensional structure they would have *in vivo*, and they adhere to a substrate with a much higher stiffness than the physiological range. Future studies should address these limitations and test if and how the observed phenotypes translate in a 3D environment.

In conclusion, we report an important step in demonstrating the key links between shape and function in astrocyte cultures, and how including cell shape dynamics within *in vitro* modelling approaches for astrocytes could be highly beneficial to better recapitulate human astrocyte biology in health and disease. We show that different morphogens drive transcriptomic, morphological, and functional differences in human astrocytes, and provide evidence that they respond differently to the same reactivity inducing factors – a long-standing question in the field of astrocyte biology. Moving forward, it will be compelling to discern what other astrocyte differentiation factors may alter astrocyte morphologies and importantly impact how these subtypes of cells function and respond to stimuli in the human adult brain.

## Supplementary information

**Figure 1 (supplementary)-.**
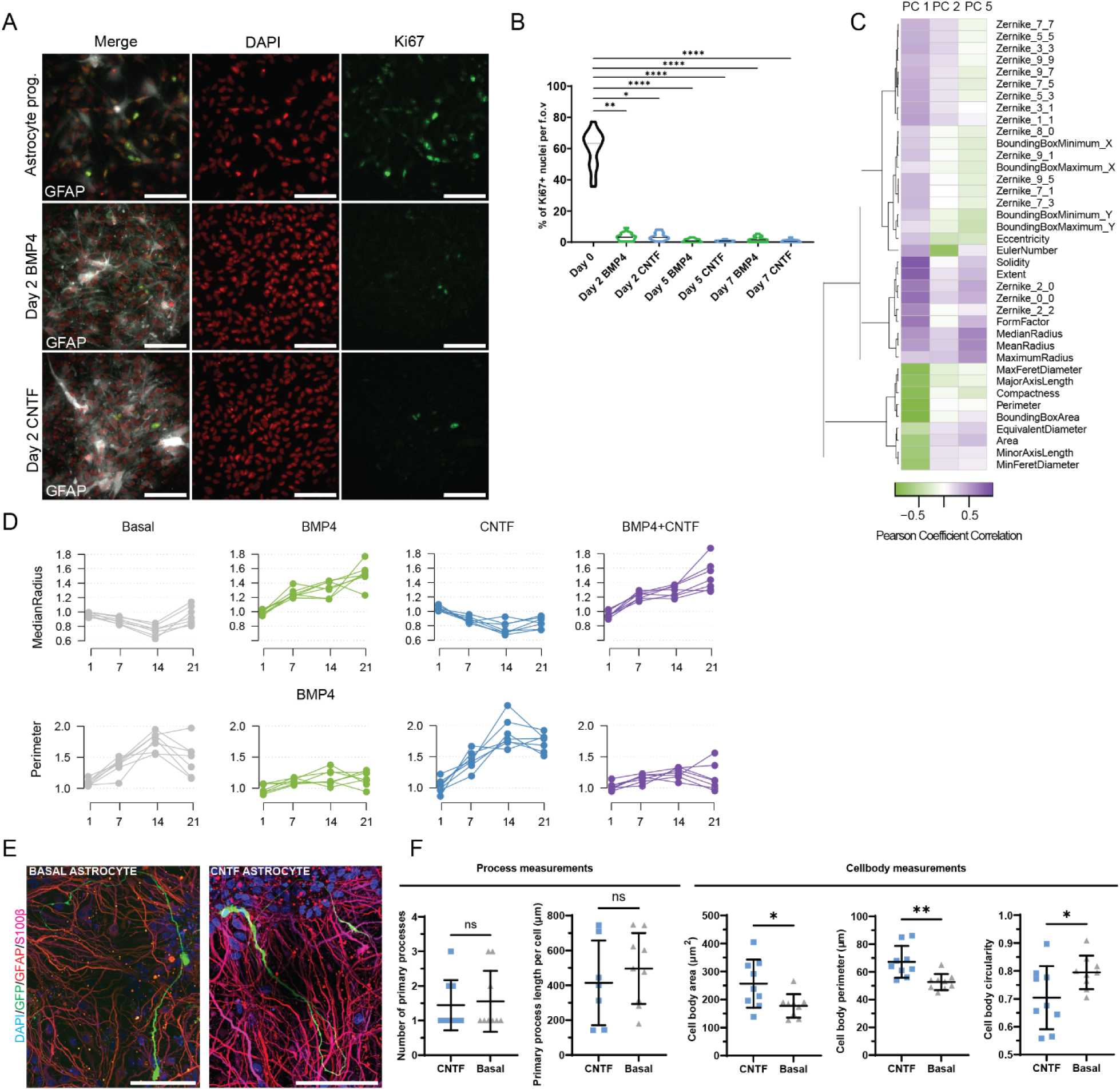
A) Representative images of astrocytes stained at multiple stages of differentiation with DAPI for nuclei detection and Ki67 for proliferating cells. (B) Quantification of DAPI/Ki67+ astrocytes from Day 2 to Day 7 of differentiation. Data from one control line within one experimental block of 6-10 technical well replicates, expressed as median ± IQR. Kruskal-wallis test with Dunn’s multiple comparisons to test significance. ****p<0.0001, ***p<0.001, **p<0.01, *p<0.05. Scale bar: 100 μm. C) Pearson correlation coefficients for principal components clustered according to shape descriptors. Purple indicates positive correlations whilst green indicates negative correlations. D) Plots of key shape descriptors changing over time for the different cytokine groups basal, BMP4, CNTF and BMP4 plus CNTF. Data points are averages of wells from one control line in one experimental block with seven technical replicates (n=7 wells). E) Representative fluorescent images of basal and CNTF differentiated astrocytes labelled with nuclear marker (DAPI), GFP and astrocyte differentiation markers (GFAP, S100β). Scale bar: 100 μm. F) Quantification of process and cell body measurements in basal and CNTF differentiated astrocytes. Data points represent single cells (n=7-9 cells) from one control line in one experimental block from three technical replicates, mean ± SD. Unpaired t-tests and Mann-Whitney test (number of processes quantification) was used to test significance ****P<0.0001, ***P<0T.001, **P<0.01, *P<0.05, ns: non-significant.

**Figure 2 (supplementary)-.**
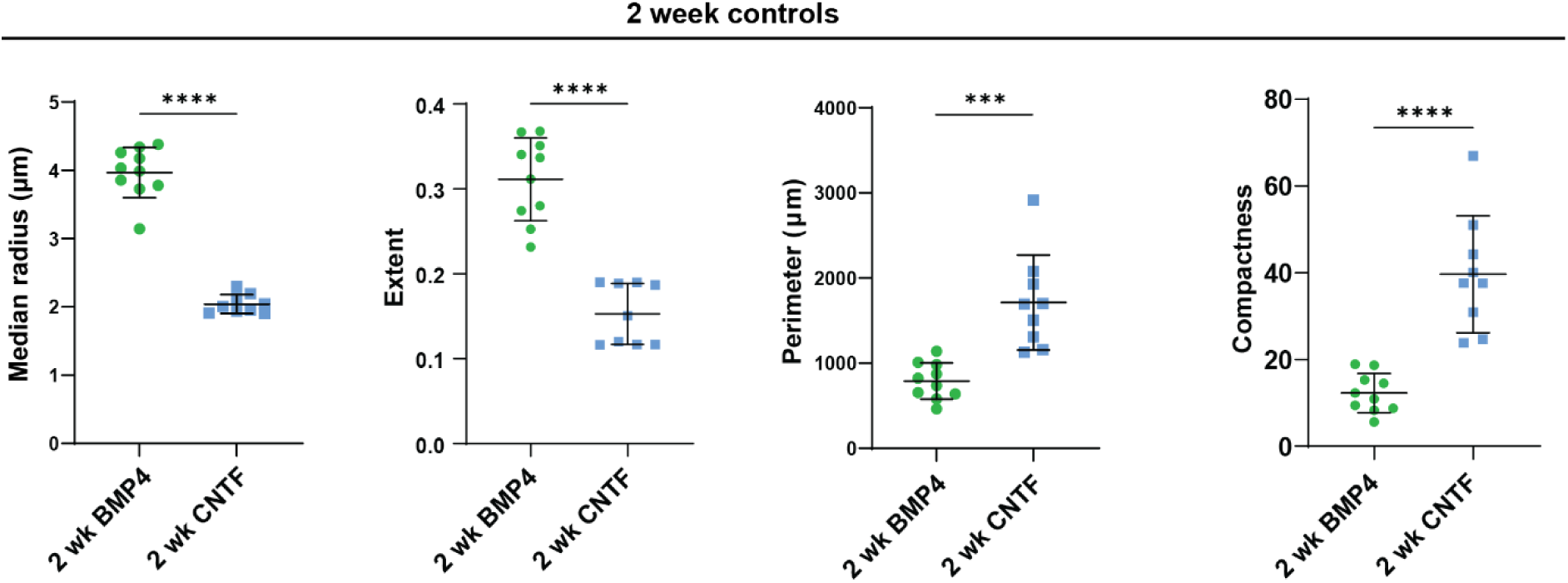
Quantification of key shape descriptors in BMP4 and CNTF differentiated astrocytes, used as a control for cytokine switching experiments. One-way ANOVA or Kruskall-Wallis test with Dunn’s multiple comparisons to test for significance between switched and non-switched pairs. Data represents technical replicate well averages of astrocytes from two control lines from two independent experiment blocks (n=7-9 wells) mean ± SD. ****p<0.0001, ***p<0.001, **p<0.01, *p<0.05, ns: non-significant. B+B: BMP4 + BMP4; B+C: BMP4 + CNTF; C+C: CNTF + CNTF; C+B: CNTF + BMP4.

**Figure 3 (supplementary)-.**
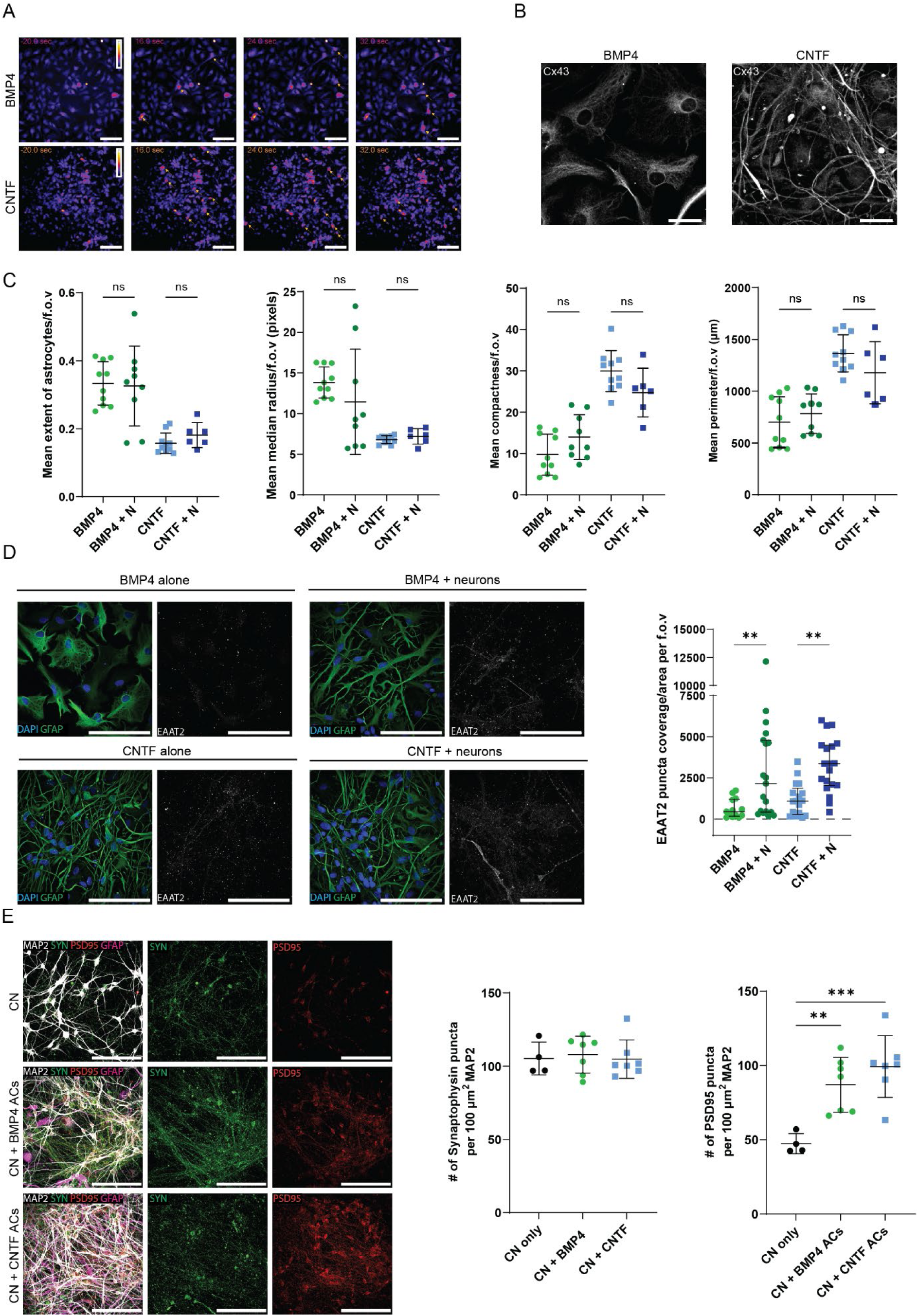
A) Example time lapse images of astrocytes labelled with Fluo4-AM and stimulated with ATP to initiate ATP-responsive calcium waves. Yellow arrows indicate astrocytes which show a peak in their fluorescence intensity during the imaging period, indicating calcium spiking behaviour. B) Representative images of BMP4 and CNTF astrocytes immunostained for connexin 43 (CX43). Scale bar: 25 μm. C) Quantification of astrocyte morphology with and without neuronal co-culture (BMP4+N: BMP4 plus neurons; CNTF+N: CNTF plus neurons) from GFP^+^ astrocytes. One way ANOVA or Kruskal-Wallis with Šídák’s multiple comparisons to test for significance between BMP4 astrocytes +/− neurons and CNTF astrocytes +/− neurons. Data points represent well averages of two control lines from two experimental blocks with 6-9 technical replicates (n = 6-9 wells), mean ± SD, ns: non-significant. Scale bar: 100 μm. D) Representative fluorescent images of BMP4 and CNTF astrocytes either cultured alone or in the presence of differentiated human stem-cell derived motor neurons and labelled with nuclear (DAPI), intermediate filament (GFAP) and glutamate transporter markers (EAAT2). Scale bar: 100 μm. Quantification of EAAT2 puncta coverage per unit area in the absence or presence of motor neurons was performed. Data points represent field of views from two control lines from two experimental blocks with six technical replicates (n = 15-18 fields of view (f.o.v)). One-way ANOVA with Šídák’s multiple comparisons to test for significance between BMP4 astrocytes +/− neurons and CNTF astrocytes +/− neurons, mean ± SD, **p<0.01. E) images of cortical neurons cultured alone (CN or in the presence of BMP4 (CN+BMP4 ACs) or CNTF (CN+CNTF ACs) astrocytes. Cortical neurons were stained for dendrites (MAP2) and pre- and post-synaptic elements with synaptophysin (SYN) and postsynaptic density-95 (PSD95), respectively. BMP4 and CNTF astrocytes were stained for their intermediate filaments (GFAP). Quantification of the number of SYN and PSD95 puncta per 100 μm^2^ MAP2 positive dendrite areas was performed comparing cortical neurons cultured alone (CN only) or in the presence of BMP4 (CN+BMP4 ACs) or CNTF (CN+CNTF ACs) astrocytes. Datapoints represent well averages of two control lines (for astrocytes) for two experimental blocks with 4-8 technical replicates (n=4-8 wells). One-way ANOVA with Dunn’s multiple comparisons to test for significance, mean ± SD, ***p<0.001, *p<0.05.

**Figure 4 (supplementary)-.**
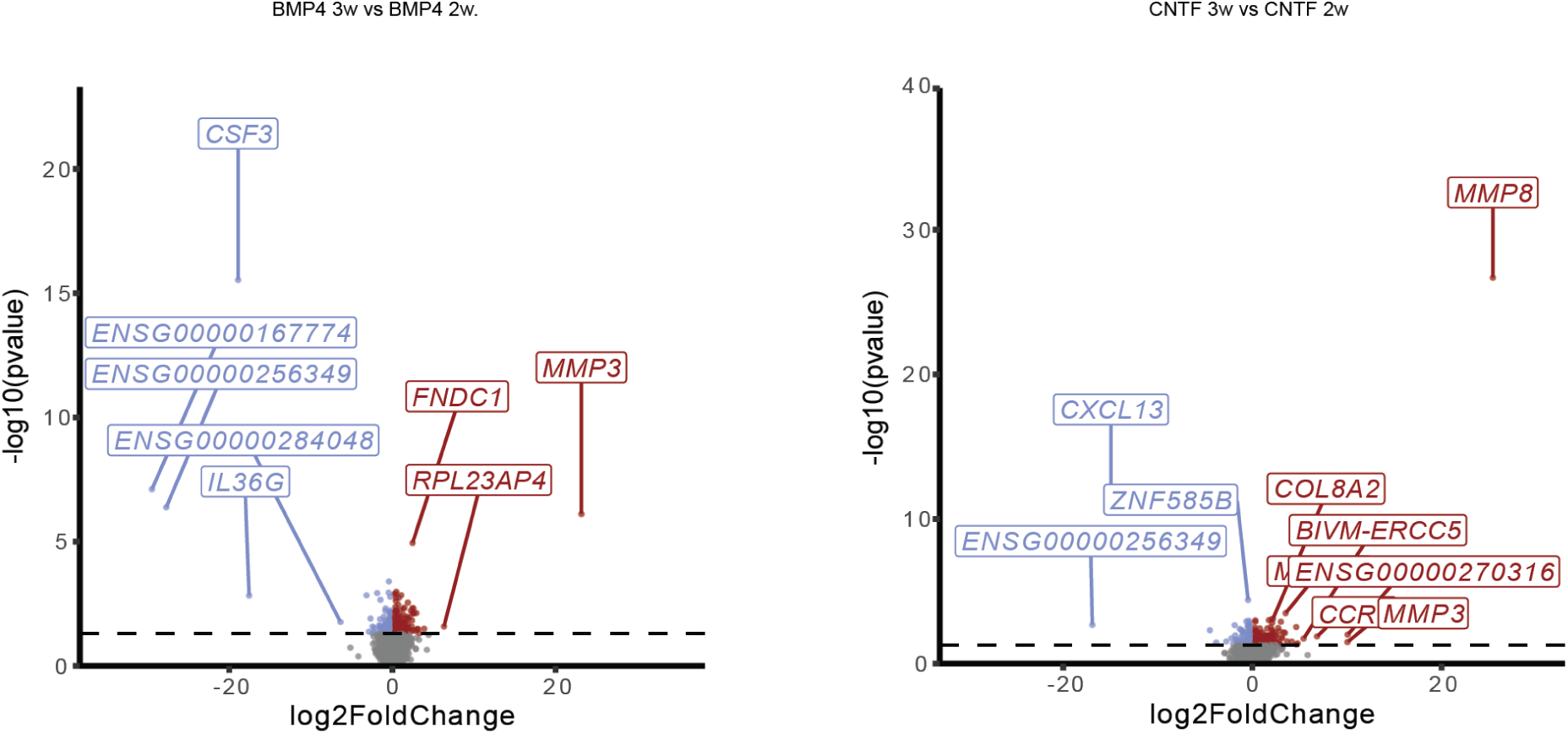
Pairwise comparison volcano plots for 3-week differentiated BMP4 astrocytes versus 2-week differentiated BMP4 astrocytes and 3-week differentiated CNTF astrocytes versus 2-week differentiated CNTF astrocytes.

**Figure 5 (supplementary)-.**
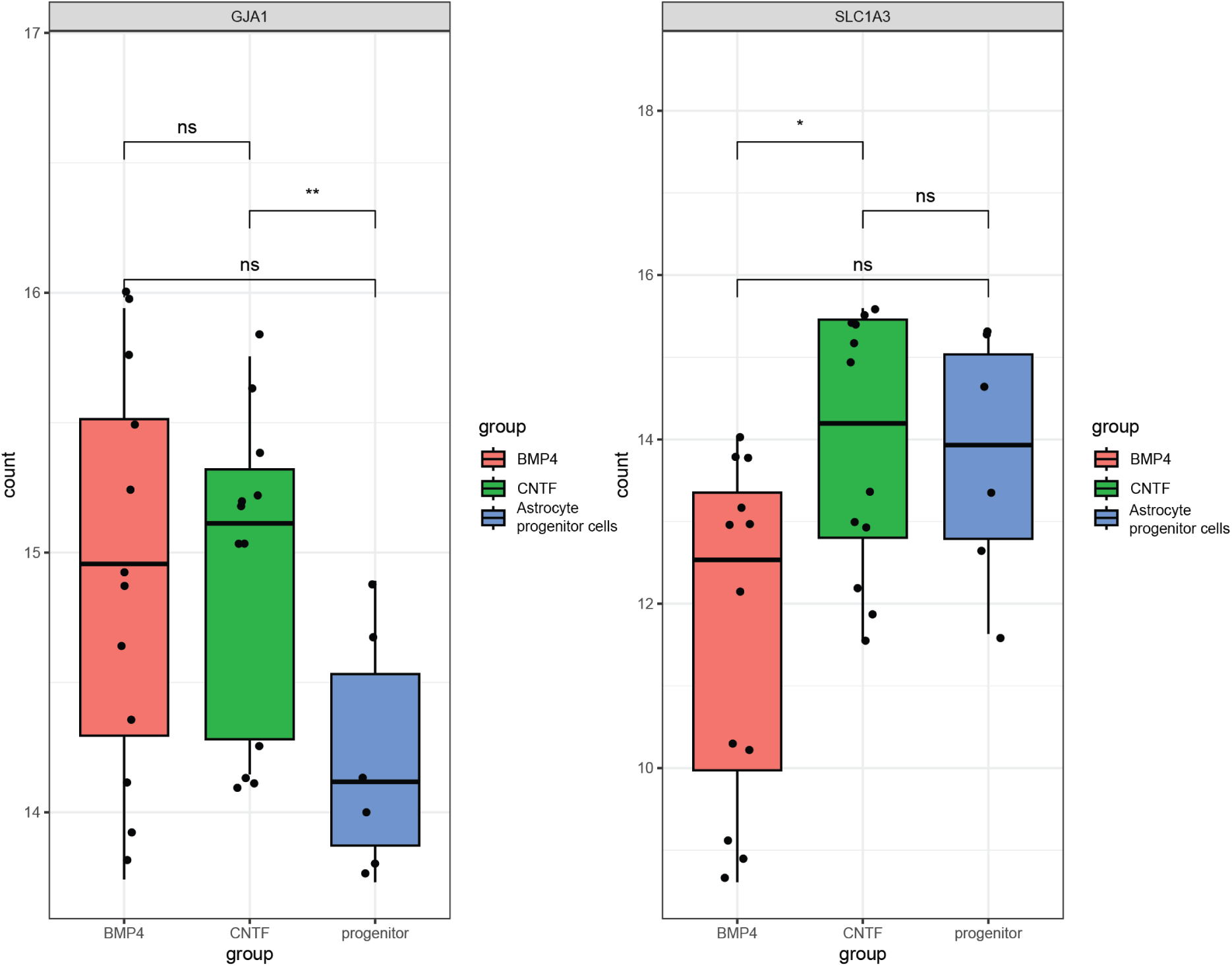
Box plots comparing the transformed expression counts of GJA1 (which encodes for CX43) and SLC1A3 (which encodes for EAAT1) between BMP4 differentiated astrocytes, CNTF differentiated astrocytes and astrocyte progenitor cells. Statistical significance for pairwise comparisons derived using Wilcoxon test (**p<0.01, *p<0.05, ns: non-significant)

**Figure 6 (supplementary)-.**
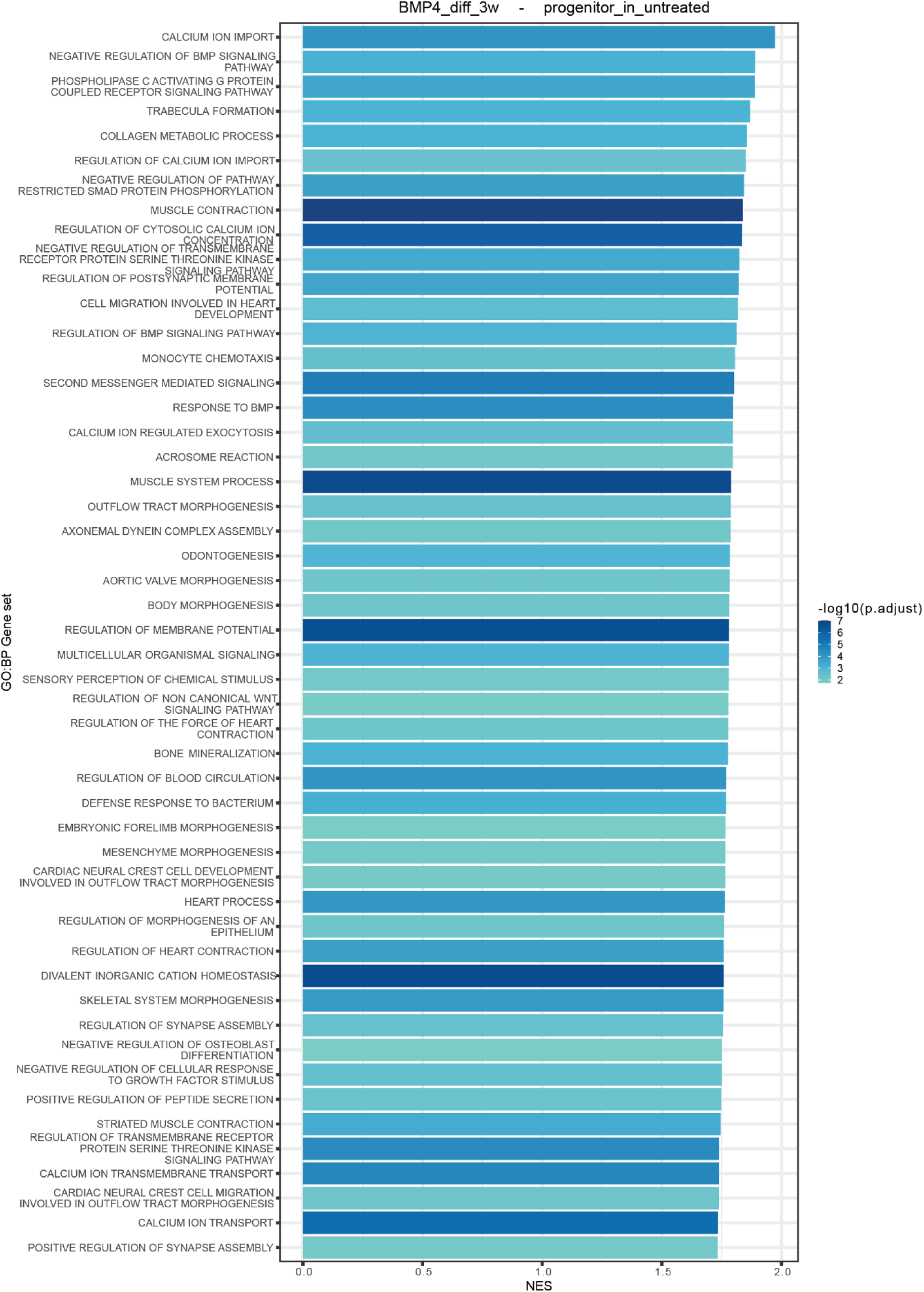
Bar plot showing top 50 gene sets upregulated in BMP4 astrocytes versus astrocyte progenitor cells. Gene sets were derived from the Biological Process (BP) ontology. Gene sets are ordered according to the absolute effect size (NES) and colored by significance (−log_10_ p-value adjusted).

**Figure 7 (supplementary)-.**
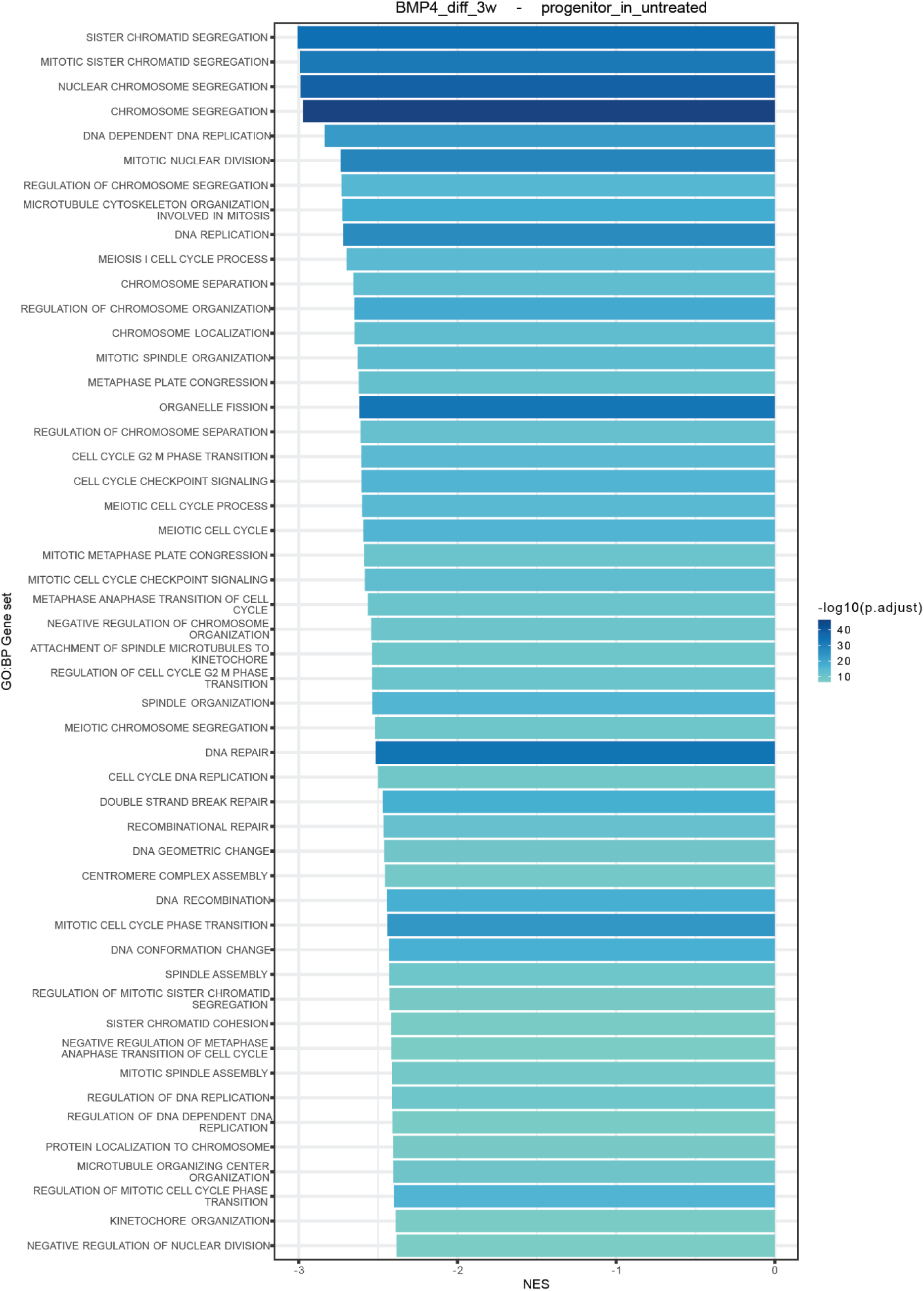
Bar plot showing top 50 gene sets downregulated in BMP4 astrocytes versus astrocyte progenitor cells. Gene sets were derived from the Biological Process (BP) ontology. Gene sets are ordered according to the absolute effect size (NES) and coloured by significance (−log_10_ p-value adjusted).

**Figure 8 (supplementary)-.**
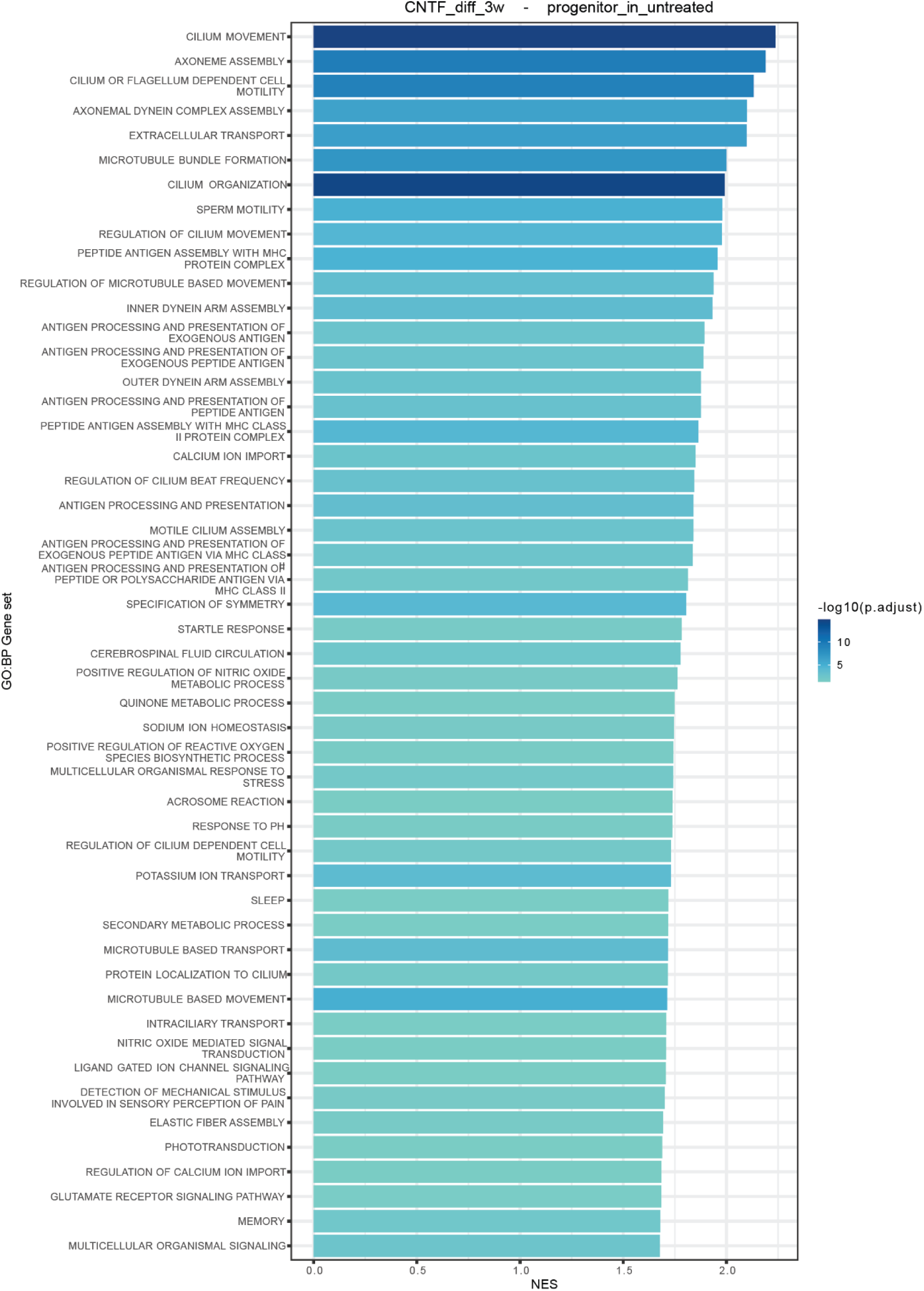
Bar plot showing top 50 gene sets upregulated in CNTF astrocytes versus astrocyte progenitors. Gene sets were derived from the Biological Process (BP) ontology. Gene sets are ordered according to the absolute effect size (NES) and colored by significance (−log_10_ p-value adjusted).

**Figure 9 (supplementary)-.**
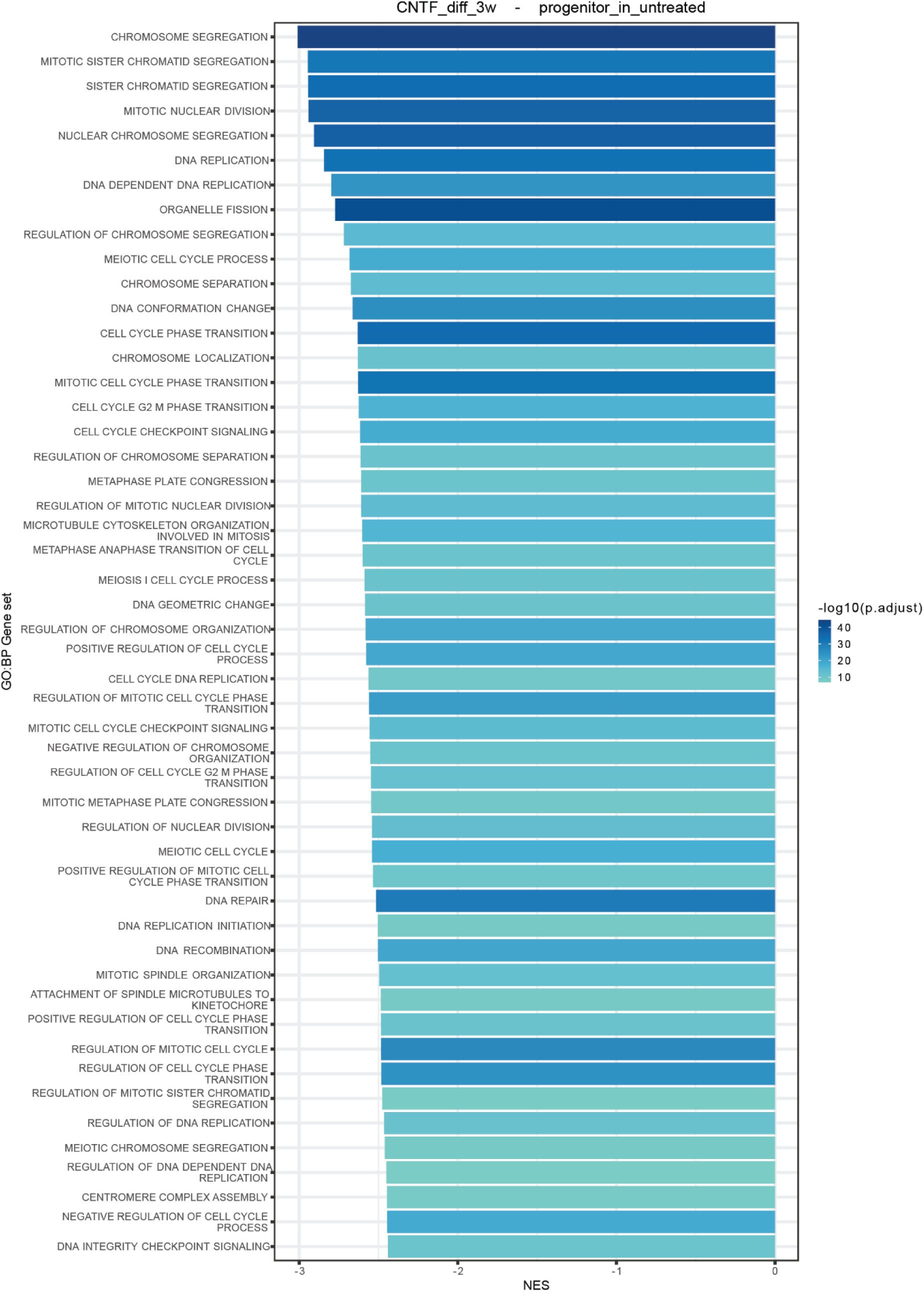
Bar plot showing top 50 gene sets downregulated in CNTF astrocytes versus astrocyte progenitors. Gene sets were derived from the Biological Process (BP) ontology. Gene sets are ordered according to the absolute effect size (NES) and colored by significance (−log_10_ p-value adjusted).

**Figure 10 (supplementary)-.**
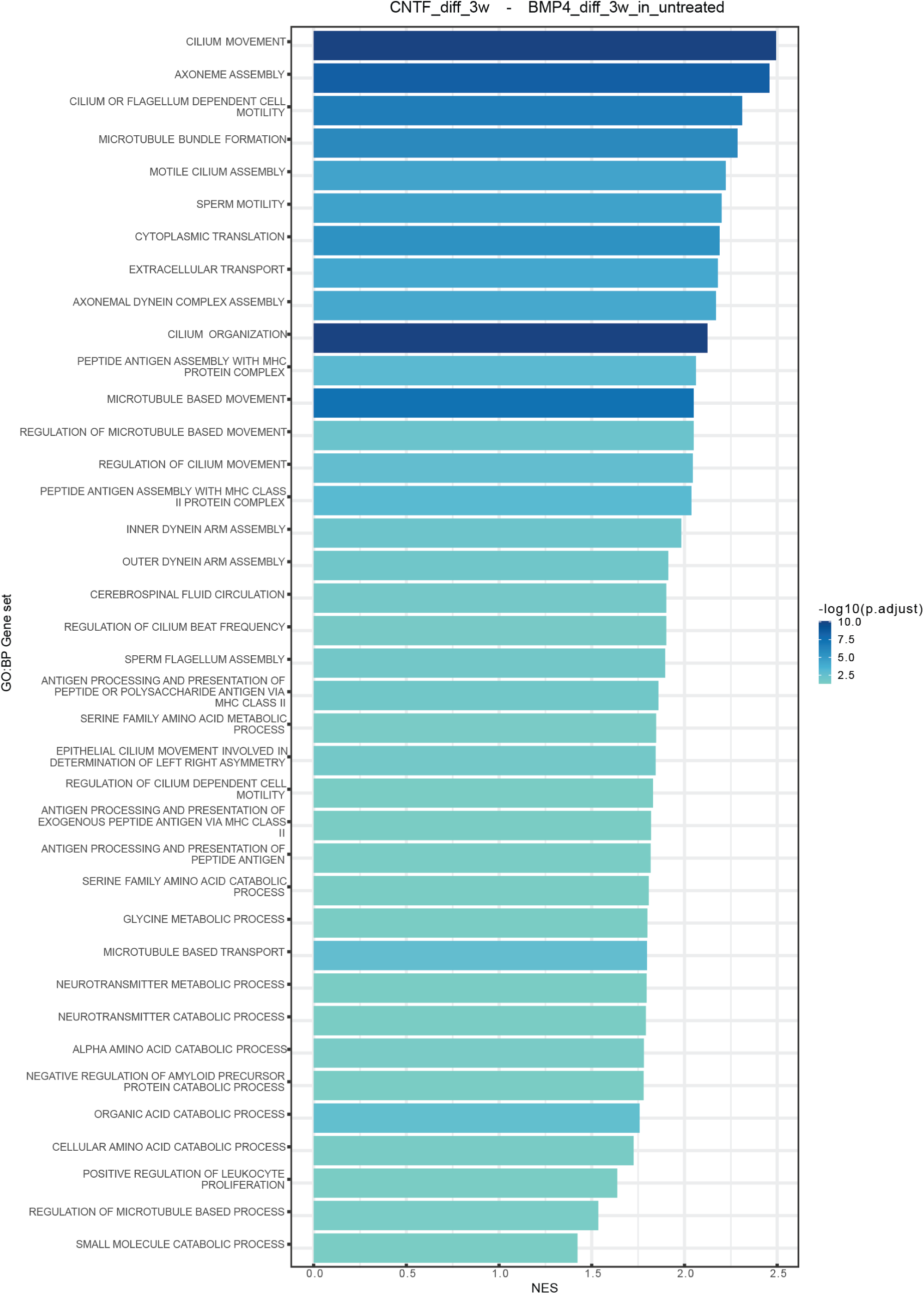
Bar plot showing top 50 gene sets upregulated in CNTF astrocytes versus BMP4 astrocytes. Gene sets were derived from the Biological Process (BP) ontology. Gene sets are ordered according to the absolute effect size (NES) and colored by significance (−log_10_ p-value adjusted).

**Figure 11 (supplementary)-.**
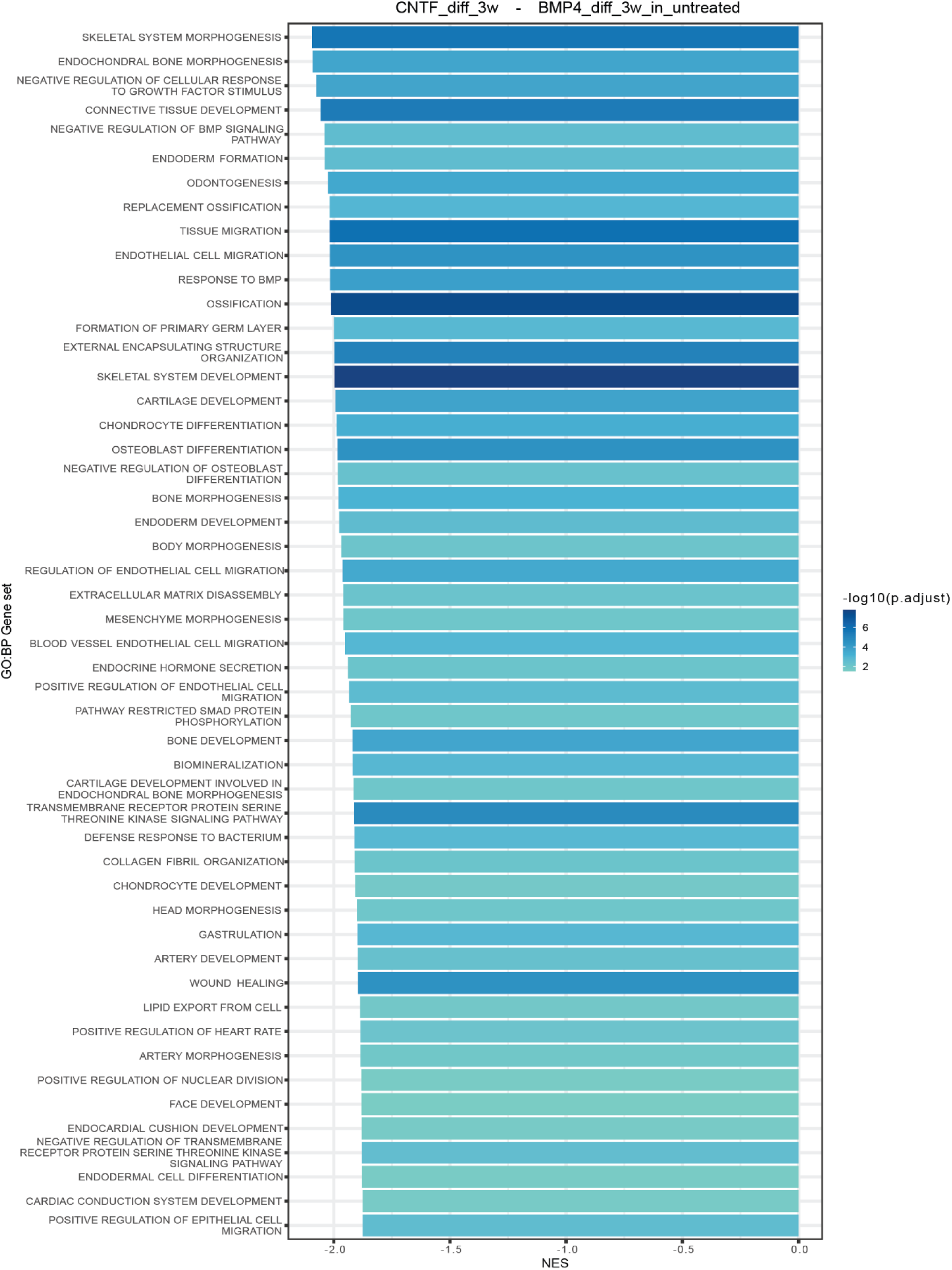
Bar plot showing top 50 gene sets downregulated in CNTF astrocytes versus BMP4 astrocytes. Gene sets were derived from the Biological Process (BP) ontology. Gene sets are ordered according to the absolute effect size (NES) and colored by significance (−log_10_ p-value adjusted).

**Figure 12 (supplementary)-.**
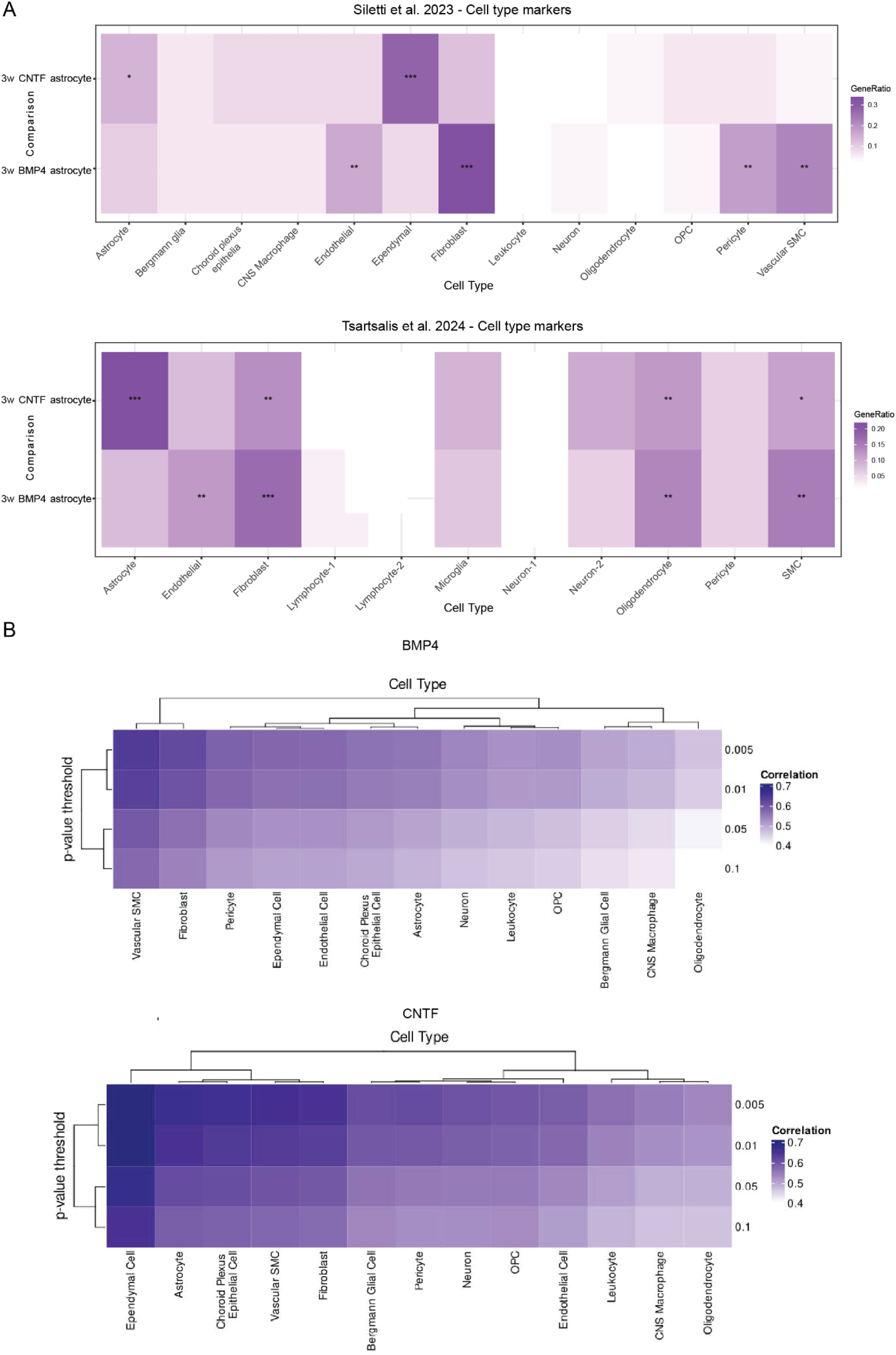
A) Comparison between the expression patterns of differentiated BMP4 and CNTF astrocytes and cell type markers from Silletti et al, 2023 and Tsartsalis et al, 2024. B) Clustered heatmap showing the correlation coefficients (Pearson’s r) between the effect sizes (log_2_FoldChange) of BMP4 or CNTF astrocytes and different cell types from Siletti et al, 2023 relative to undifferentiated progenitors at different BMP4- and CNTF-differentiated astrocyte DEG adjusted p-value thresholds.

**Figure 13 (supplementary)-.**
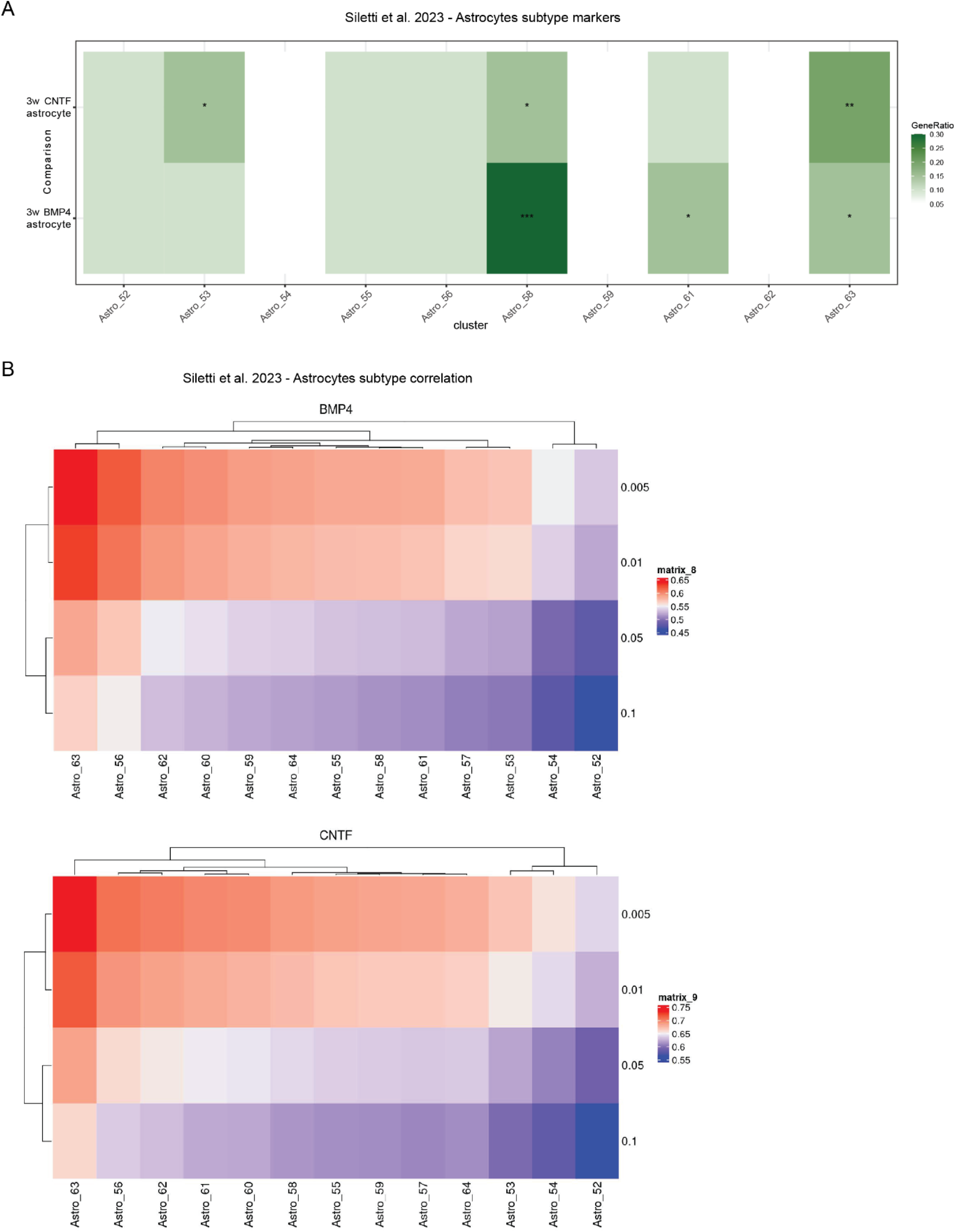
A) Comparison between the expression patterns of differentiated BMP4 and CNTF astrocytes and astrocyte subtype markers from Silletti et al, 2023. B) Correlation matrices showing the amount of correlation between the expression patterns of BMP4 or CNTF astrocytes and different astrocyte subtypes from Silletti et al, 2023

**Figure 14 (supplementary)-.**
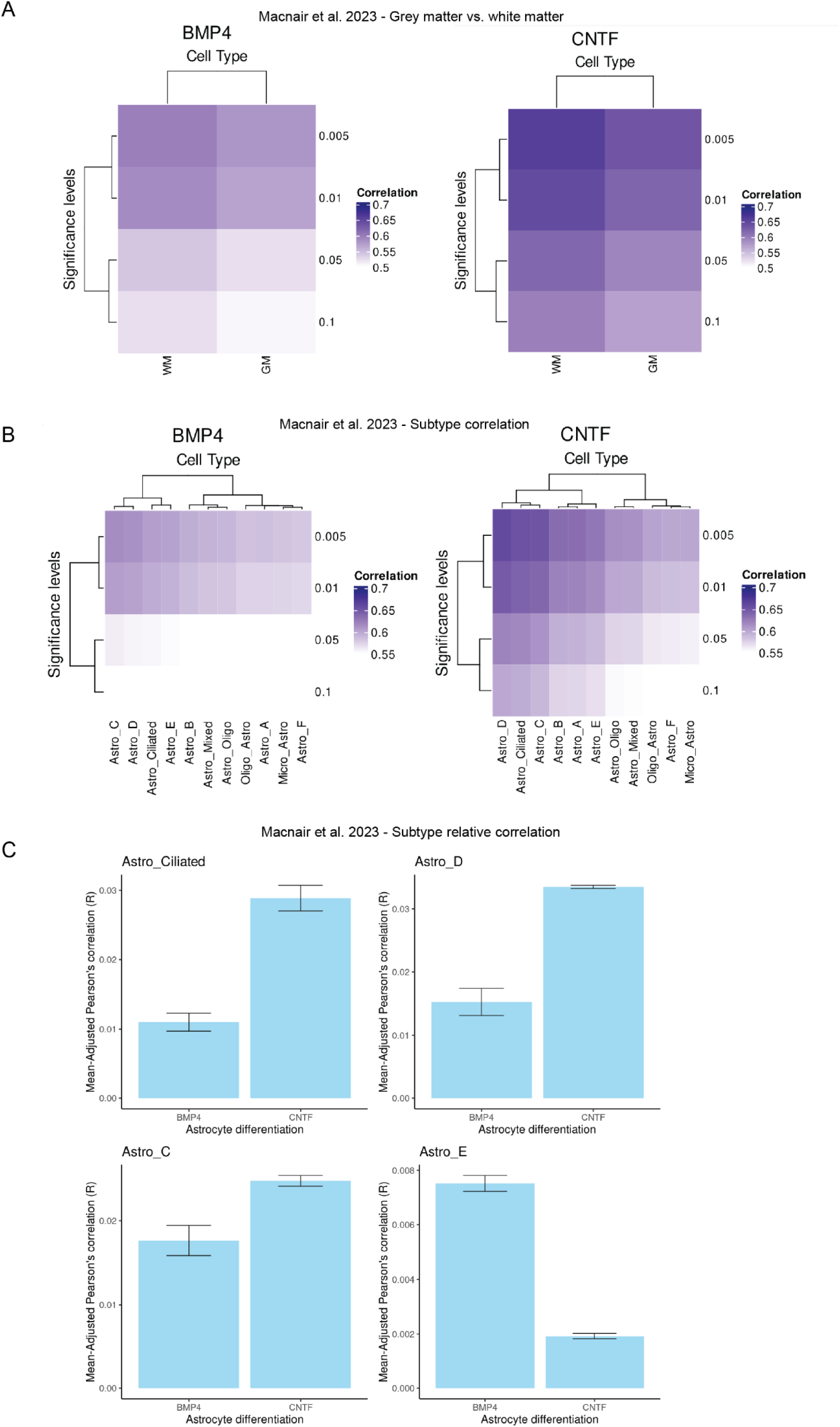
A) Comparison between the expression patterns of differentiated BMP4 and CNTF astrocytes and those of white and grey matter astrocytes from Macnair et al, 2023. B) Correlation matrices showing the amount of correlation between the expression patterns of BMP4 or CNTF astrocytes and different astrocyte subtypes from Macnair et al, 2023. C) Bar blots showing the relative correlation between BMP4 and CNTF astrocytes and the top subtypes from Macnair et al, 2023.

**Figure 15 (supplementary)-.**
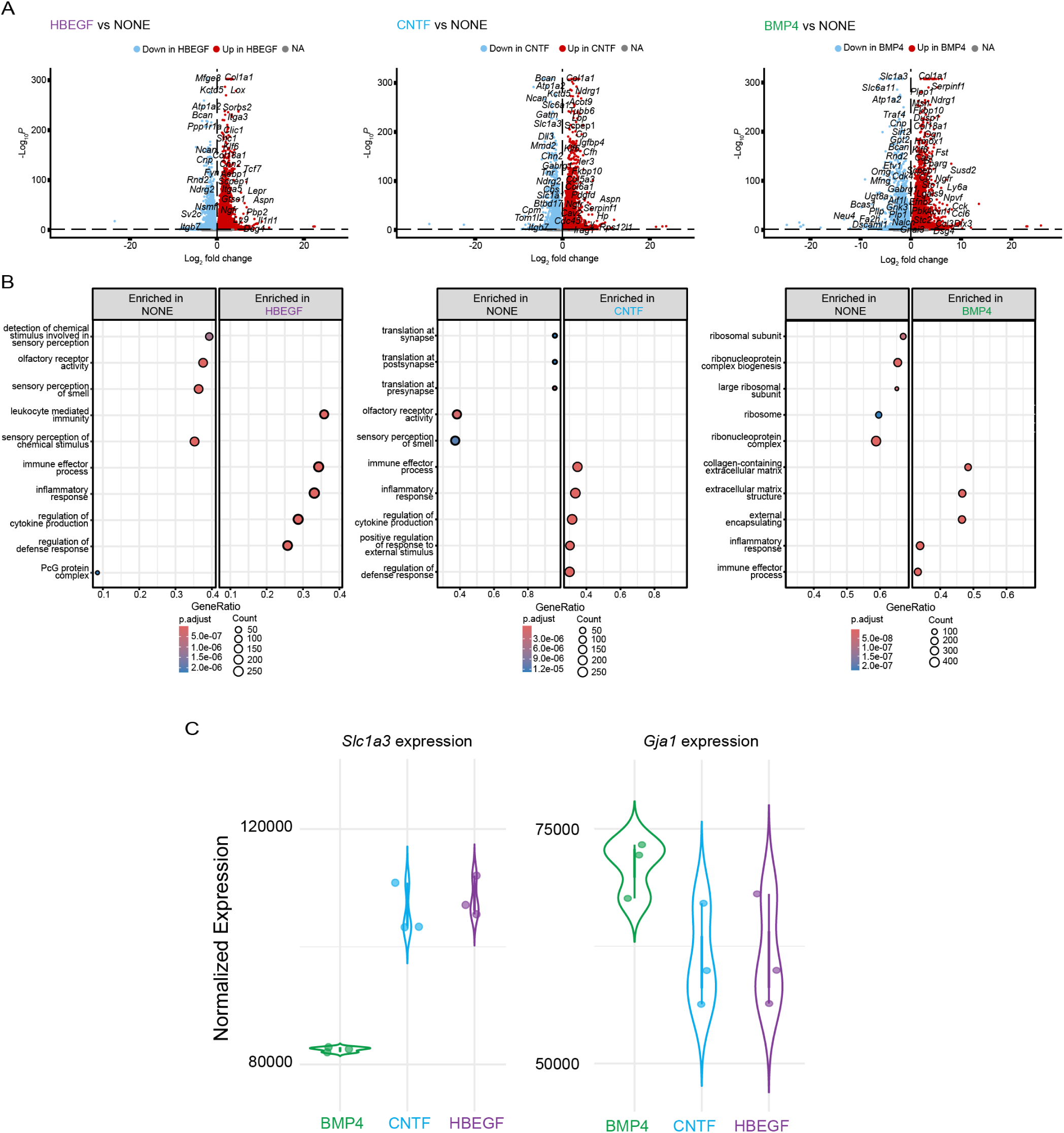
A) Volcano plots of differentially expressed genes between respective treatment conditions. Positive log_2_ fold change is representative of enrichment while negative log_2_ fold change is representative of de-enrichment. B) Dot plotting of GO biological process gene set enrichment. Gene ratio describes the ratio of significantly changed genes to the total genes in each pathway set. Dot size represents the total number of enriched or de-enriched genes while color is representative of adjusted p-value. C) Violin plotting of individual gene expression across n=3 technical replicates split by treatment condition. Counts normalized by DESeq2 median of ratios.

**Figure 16 (supplementary)-.**
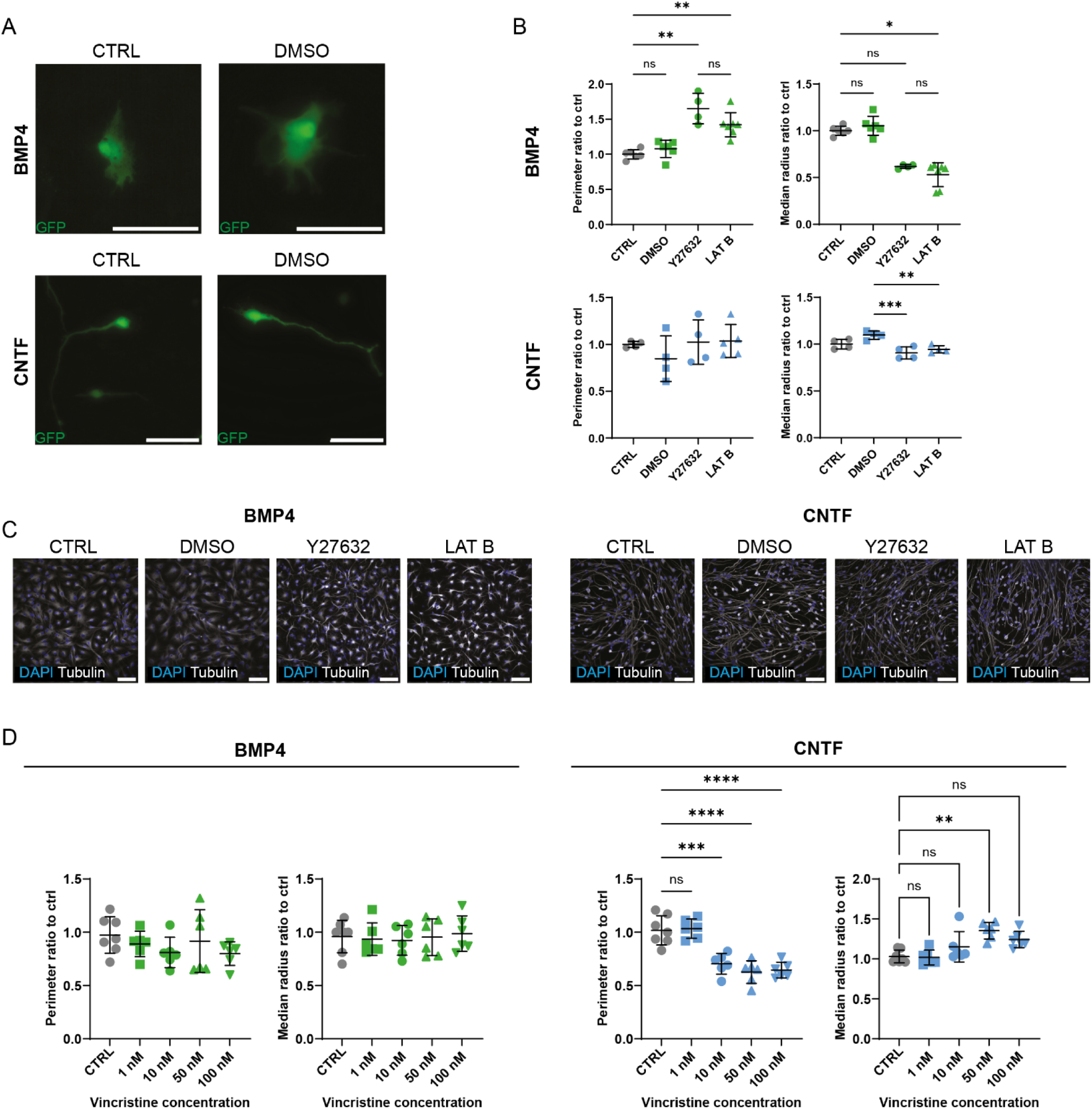
A) Representative images of GFP^+^ BMP4 and CNTF astrocytes, either untreated (CTRL) or treated with vehicle control DMSO. Scale bar: 100 μm. *B)* Quantification of shape changes in the presence of actin inhibitors analysing perimeter and median radius expressed as a ratio compared to control (ctrl, black). Data represents well averages of technical replicates of two control lines from two experimental blocks (n=4-5 wells total), mean ± SD. Ordinary one-way ANOVAs with Tukey’s post-hoc was used to test for significance. *p<0.05, **p<0.01, ****p<0.0001 ns or no comparisons shown: non-significant. *C)* Representative images of BMP4 and CNTF astrocytes, either untreated (ctrl) or treated with actin inhibitors Y27632 dihydrochloride (Y27632; 25 μM) or latrunculin B (LAT B; 1 μM) for 2 hours, stained with nuclear (DAPI, blue) and microtubule cytoskeletal markers (Tubulin, white). Scale bar: 100 μm. D) Quantification of shape changes in the presence of vincristine analysing perimeter and median radius, expressed as a ratio compared to control (ctrl, black). Data represents well averages of technical replicates of two control lines from two experimental blocks (n=6-7 wells total), mean ± SD. Ordinary one-way ANOVAs with Tukey’s post-hoc was used to test for significance.

**Figure 17 (supplementary)-.**
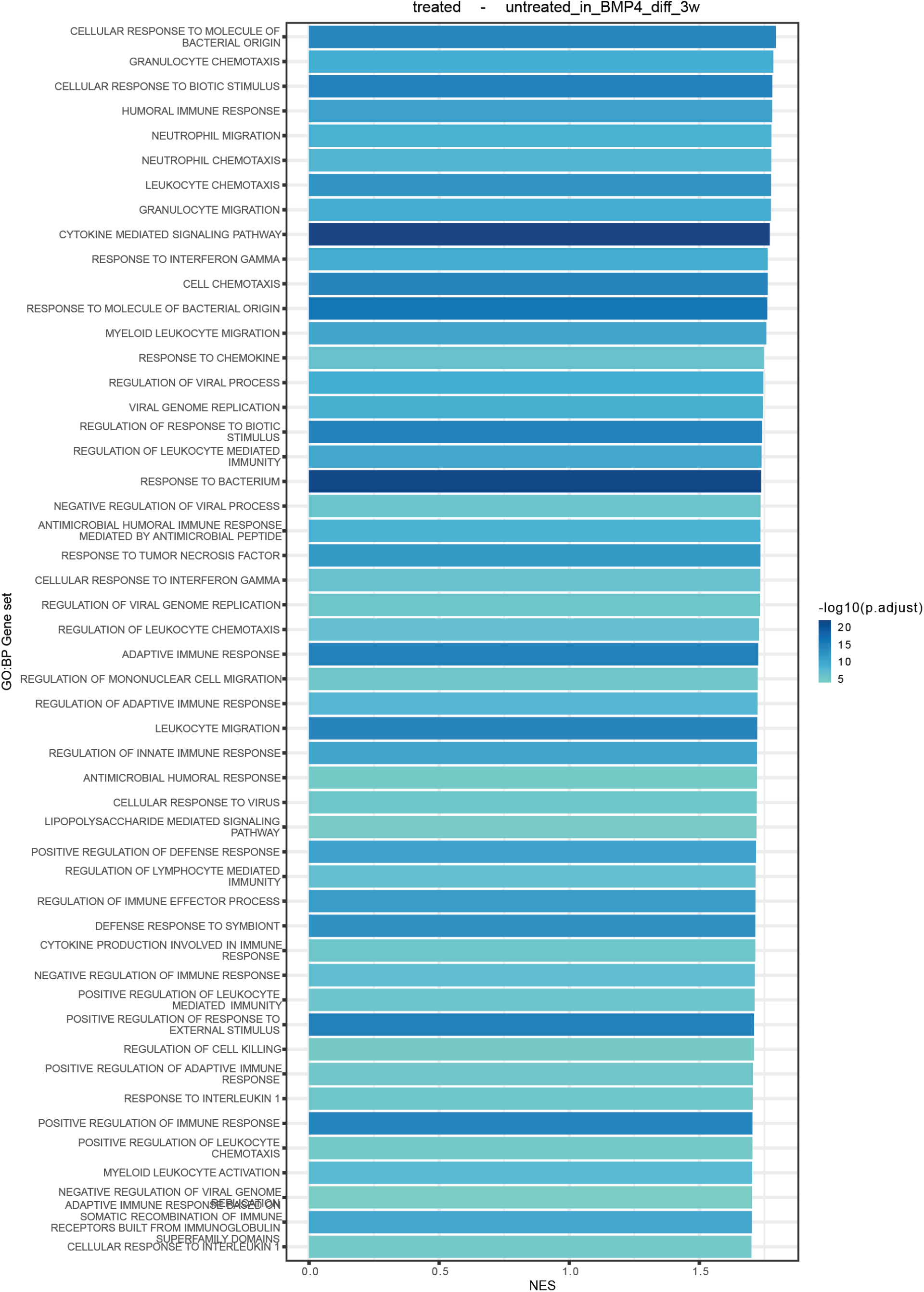
Bar plot showing the gene sets upregulated in treated BMP4 astrocytes versus untreated BMP4 astrocytes. Gene sets were derived from the Biological Process (BP) ontology. Gene sets are ordered according to the absolute effect size (NES) and colored by significance (−log_10_ p-value adjusted).

**Figure 18 (supplementary)-.**
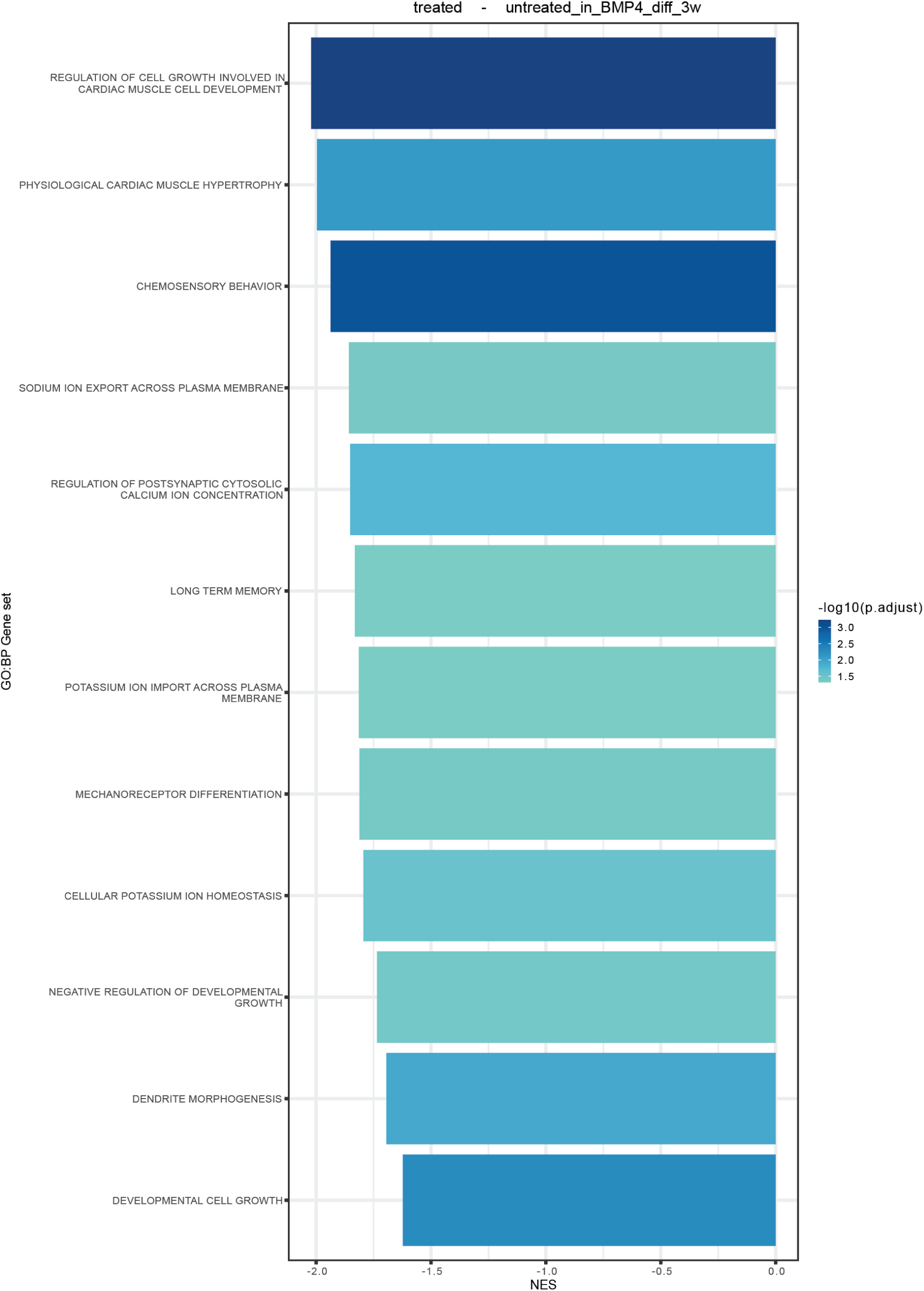
Bar plot showing the gene sets downregulated in treated BMP4 astrocytes versus untreated BMP4 astrocytes. Gene sets were derived from the Biological Process (BP) ontology. Gene sets are ordered according to the absolute effect size (NES) and colored by significance (−log_10_ p-value adjusted).

**Figure 19 (supplementary)-.**
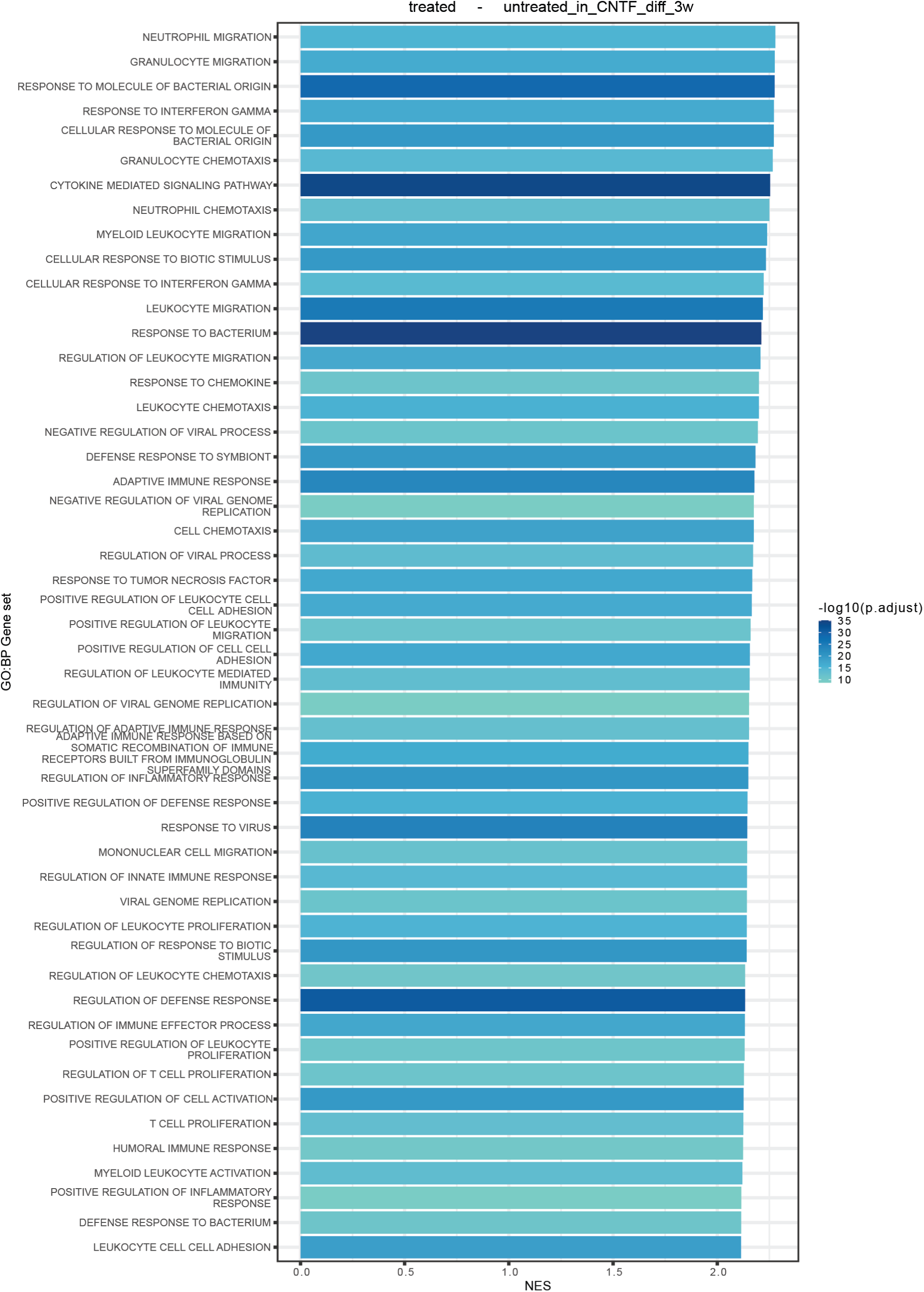
Bar plot showing the gene sets upregulated in treated CNTF astrocytes versus untreated CNTF astrocytes. Gene sets were derived from the Biological Process (BP) ontology. Gene sets are ordered according to the absolute effect size (NES) and colored by significance (−log_10_ p-value adjusted).

**Figure 20 (supplementary)-.**
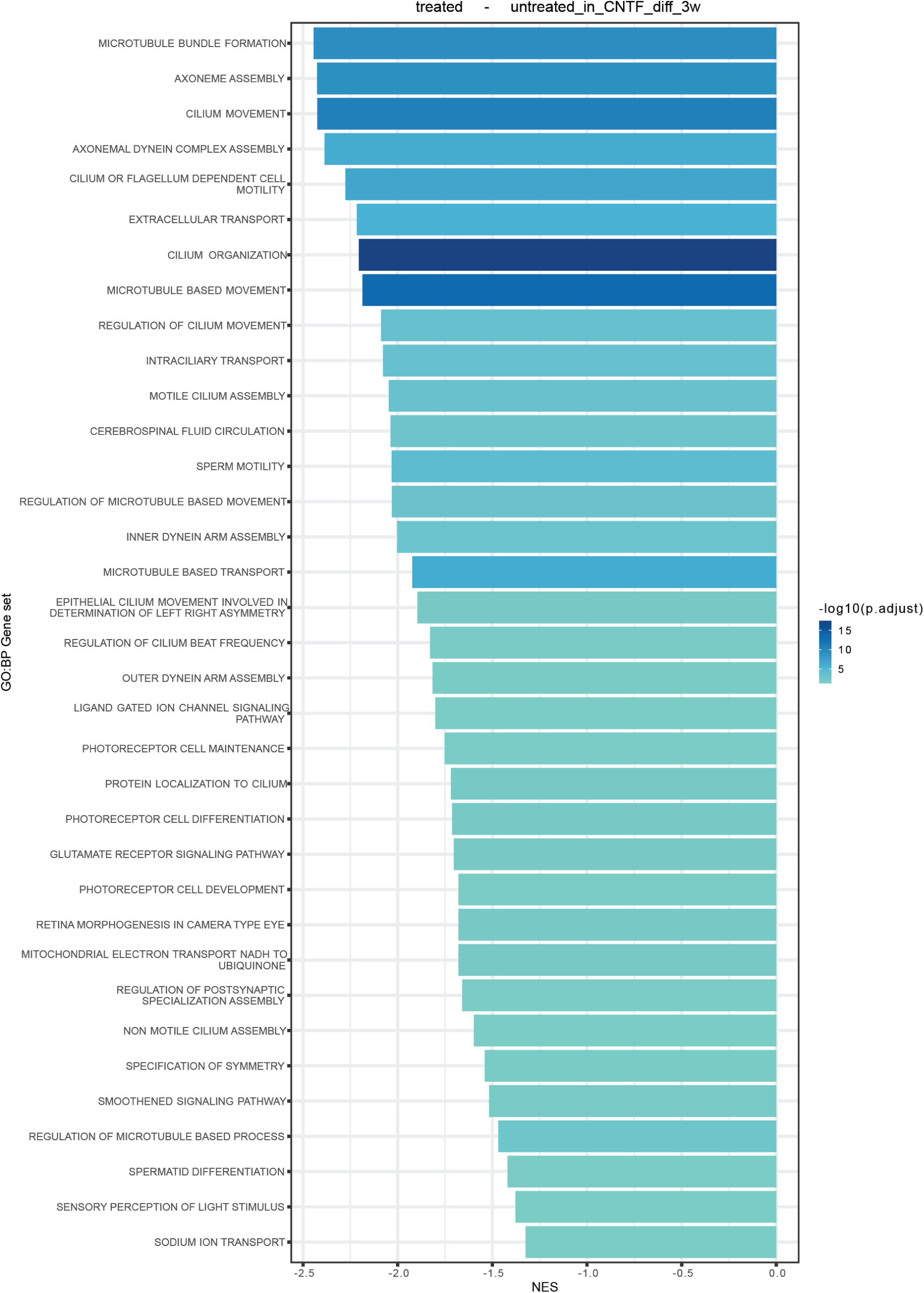
Bar plot showing the gene sets downregulated in treated CNTF astrocytes versus untreated CNTF astrocytes. Gene sets were derived from the Biological Process (BP) ontology. Gene sets are ordered according to the absolute effect size (NES) and colored by significance (−log_10_ p-value adjusted).

**Figure 21 (supplementary)-.**
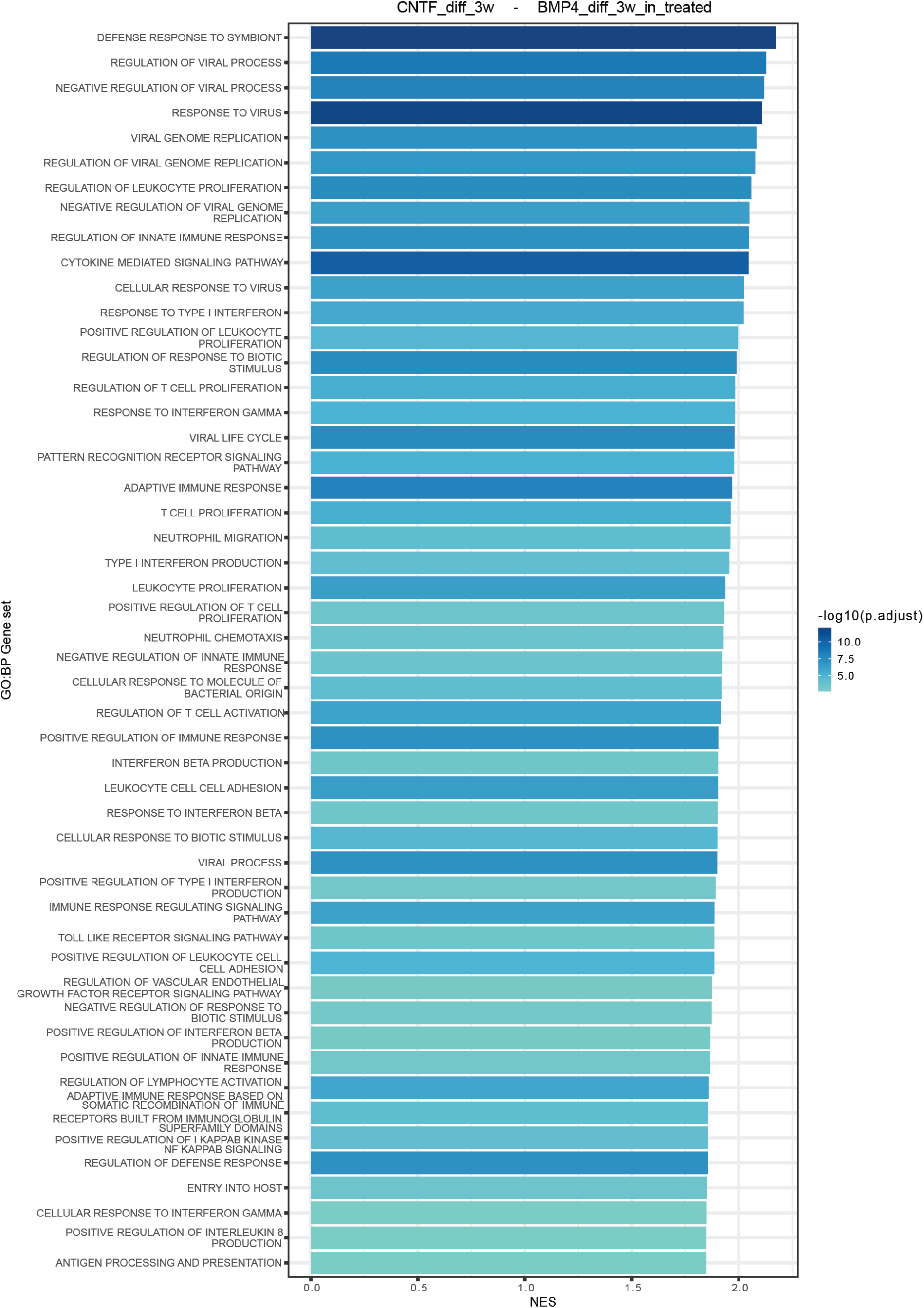
Bar plot showing the gene sets upregulated in treated CNTF astrocytes versus treated BMP4 astrocytes. Gene sets were derived from the Biological Process (BP) ontology. Gene sets are ordered according to the absolute effect size (NES) and colored by significance (−log_10_ p-value adjusted).

**Figure 22 (supplementary)-.**
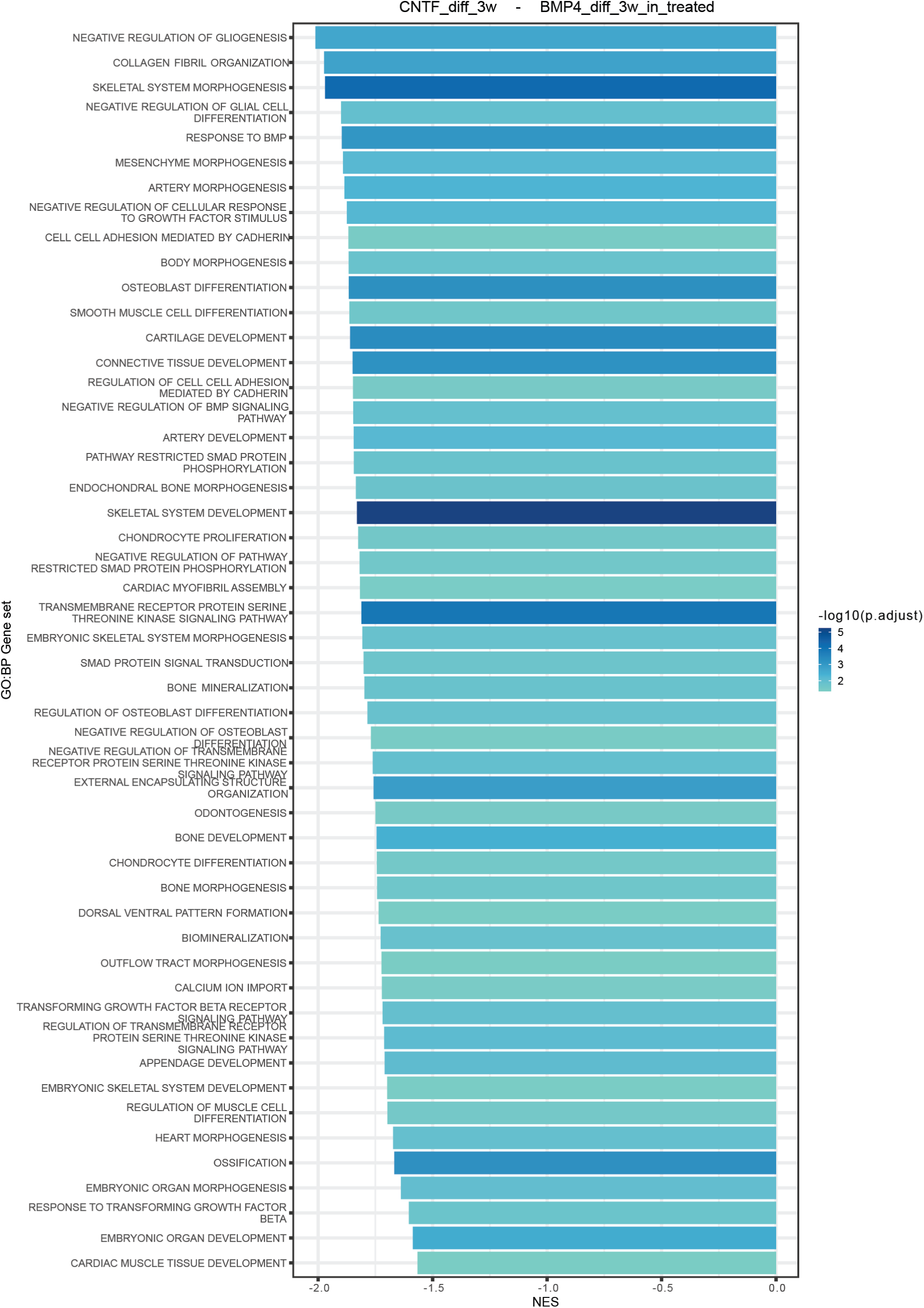
Bar plot showing the gene sets downregulated in treated BMP4 astrocytes versus untreated BMP4 astrocytes. Gene sets were derived from the Biological Process (BP) ontology. Gene sets are ordered according to the absolute effect size (NES) and colored by significance (−log_10_ p-value adjusted).

## Acknowledgements

A.S. wishes to acknowledge the UK-Dementia Research Institute and the Medical Research Council, which supported a part of this study with a programme grant funding.S.J.M. is supported by the UK Dementia Research Institute (award number UKDRI-6009) through UK DRI Ltd, principally funded by the UK Medical Research Council and a Medical Research Council grant (award number MR/W004984/1).

## Ethics declaration

The authors declare no competing interests.

## Materials and Methods

### Cell culture

All cells were kept in humidified atmosphere at 37°C with 5% CO_2_. Human iPSCs were cultured on plates coated with Matrigel (Corning; 1:100), fed with Essential 8 Flex medium (Gibco) and passaged with ethylenediaminetetraacetic acid with phosphate-buffered saline (EDTA-PBS) (Gibco, 0.5 mM).

Motor neuron progenitors were generated and patterned to ventral spinal cord identity following a previously published protocol^20^.

Cortical neuron progenitors were generated according to a previously published protocol^75^.

Both these progenitors were cultured in a base medium containing 50% advanced Dulbecco’s modified eagle’s medium (DMEM) (Gibco) and 50% NeuroBasal (Gibco) with B27 (Gibco) and N2 (Gibco) supplements, Penicillin-Streptomycin antibiotic mix (Gibco, 100 μg/ml) and GlutaMAX (Gibco, 1X). Base medium was supplemented with FGF-2 (Gibco, 20 ng/ml) (F20 medium).

Astrocyte progenitors were obtained following a published protocol^19^. Briefly, neurospheres were formed starting from overconfluent motor neuron progenitors and cultured on an orbital shaker (60 rpm) in F20 medium for 2 weeks. They were then mechanically chopped at the start of the glial enrichment phase and cultured for 2-4 weeks in base medium supplemented with EGF (Gibco, 20 ng/ml) and leukaemia inhibiting factor (LIF) (Sigma-Aldrich, 20 ng/ml). After the enrichment, spheres were propagated in base medium containing EGF (20 ng/ml) and FGF-2 (20 ng/ml) (EF20 medium). After 1-2 weeks the spheres were landed in Matrigel-coated plates (1:100) to allow a population of monolayer astrocytes to migrate out of them.

Single cell suspensions of astrocyte progenitor cells were isolated and then propagated in EF20 medium and passaged with Accutase (StemPro) when they reached confluency.

For neuronal differentiation of motor neuron progenitors and cortical neuron progenitors, base medium supplemented with compound E (Enzo, 0.1 μM) was used.

For differentiation of astrocyte progenitors into astrocytes, NeuroBasal medium was used. This was supplemented with B27 (Gibco), Non-Essential Amino Acids (Gibco, 1X), Penicillin-Streptomycin antibiotic mix (Gibco, 100 μg/ml), GlutaMAX (Gibco, 1X) and either BMP4 (Gibco, 10 ng/ml) or CNTF (Gibco, 20 ng/ml) or a combination of both (respectively, NB27+BMP4, NB27+CNTF, NB27+BOTH). Throughout the differentiation process (the duration is specified for each experiment) cultures were fed every other day.

**Table 1-.**
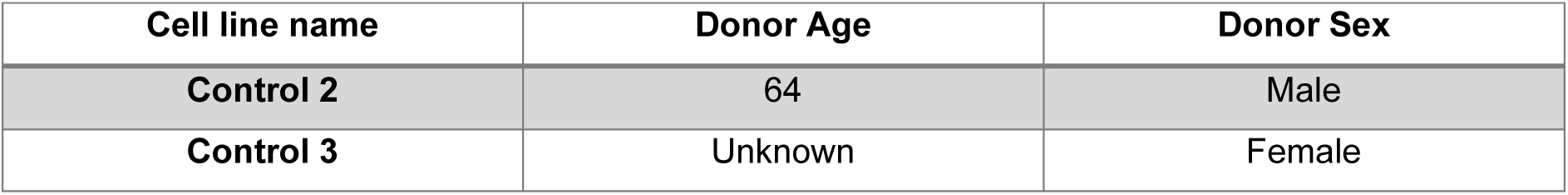
iPSC lines.

#### Constructs

The mito-dendra2 construct was derived from the Addgene plasmid #55796 (http://n2t.net/addgene:55796; RRID: Addgene_55796), contributed from David Chan. Cloning was performed using the GeneArt assembly kit (Invitrogen) to insert the mito-dendra2 sequence into a PiggyBac (PB) backbone construct with a T2A Neomycin selection cassette. Other PiggyBac constructs were cloned in the laboratory, such as PB-EF1α-GFP-PGK-Puro and PB-CAG-RFP-Hygro for stable transfection of GFP and RFP. When stable transfections were required, these vectors were co-transfected with a construct encoding the PB transposase.

#### Chemical transfection of astrocyte progenitors

Stable lines of astrocyte progenitors expressing PB-Ef1α-GFP-PGK-Puro were established to perform single cell shape analysis aiming for 5-10% transfection efficiency. Stable co-transfection of PB-CAG-RFP-Hygro and PB-Ef1α-mito-dendra2-Neo was carried out to allow visualisation of mitochondrial distribution in single cells. All transfections were performed via lipofection using TransIt-LT1 transfection reagent (Mirus). Astrocyte progenitors were passaged into 24 well plates one day before the transfection. A total of 1μg of DNA per well of a 24 well plate was used in 200 µL of pen-strep free EF20 medium.

This solution was well mixed and added to the wells which also contained 200 µL of penicillin-streptomycin (Pen-Strep) free medium. The medium containing the transfection reagent was exchanged with growth medium after 24 h. Cells were cultured to confluency and then pooled into a 6-well plate and there further expanded.

### Treatment with reactivity factors

To induce reactivity, 2-weeks differentiated astrocytes were cultured for one additional week in the same medium supplemented with reactivity factors TNF (Cell Signalling Technology, 30 ng/ml), IL-1α (Sigma, 1 μg/ml) and C1q (Abcam, 400 μg/ml) (TIC).

After treatment cells were either fixed, stained and processed for morphological analysis and protein quantification or scratched and live images (following the methods documented in relevant sections).

### Treatment with cytoskeleton disrupting drugs

For the disruption of actin cytoskeleton, 2 weeks differentiated BMP4 and CNTF astrocytes were treated with ROCK inhibitor Y27632 (Tocris, 25 μM) or LAT B (Sigma, 1 μM) for 2 hours. A vehicle control for LAT B was carried out using 0.1% (v/v) dimethyl sulfoxide (DMSO, Sigma). For the disruption of microtubules, 2 weeks differentiated BMP4 and CNTF astrocytes were treated with a range of concentrations (1 nM, 10 nM, 50 nM, 100 nM) of Vincristine (ThermoFisher) for 24 hours. After treatment, cells were fixed and imaged at 20x magnification on a Nikon Tie Eclipse microscope.

For all experiments, an untreated control was carried out.

#### Immunostaining and live dyes

For immunostaining, cells were fixed using 4% paraformaldehyde (PFA) for a duration of 15 minutes. This was followed by 3 washes with PBS. Cell permeabilization was achieved through 10 minutes incubation with 0.1% Triton-X (Sigma) in PBS. This was then replaced with the blocking solution, consisting of 3% goat serum (GS, Sigma) in PBS, which was incubated for 30 minutes. Primary antibodies were diluted in a solution of 0.05% Triton 1.5% GS in PBS and incubated for 1 hour at room temperature, followed by 3 x PBS washes. Details of primary antibodies and their concentrations can be found in Table 2. Secondary antibodies were diluted in the same solution as the primary and incubated for 30 minutes, followed by 3 x PBS washes. All secondary antibodies were employed at a final concentration of 2 μg/ml. Details of the secondary antibodies used can be found in table 3. DAPI (Sigma) was used in combination with the secondary antibodies at a final concentration of 1 μg/ml.

**Table 2-.** Primary antibodies.

| Antibody | Host species | Isotype | Manufacturer | Catalogue code | Dilution |
| --- | --- | --- | --- | --- | --- |
| **GFAP-Cy3** | Mouse | IgG1 | Sigma | C9205 | 1:500 |
| **GFAP** | Chicken | Polyclonal | Abcam | Ab4674 | 1:1250 |
| **S100β** | Mouse | IgG1 | Sigma | S2532 | 1:1000 |
| **Ki67** | Rabbit | IgG1 | Abcam | Ab92742 | 1:500 |
| **Cx43** | Mouse | IgG1 | ThermoFisher | 13-8300 | 1:500 |
| **GFP** | Rabbit | Polyclonal | Abcam | Ab6556 | 1:500 |
| **β3-tubulin** | Mouse | IgG2b | Sigma | T5076 | 1:1000 |
| **MAP2** | Mouse | IgG1 | Sigma | 13-1500 | 1:750 |
| **Synaptophysin** | Rabbit | Polyclonal | ThermoFisher | PA1-1043 | 1:500 |
| **α-tubulin** | Mouse | IgG2a | ThermoFisher | MA5-31466 | 1:500 |
| **C3d** | Rabbit | Polyclonal | Dako | A0063 | 1:500 |
| **PSD95** | Mouse | IgG2a | ThermoFisher | MA1-045 | 1:500 |
| **EAAT1** | Rabbit | Polyclonal | Novus Biologicals | NBP1-84940 | 1:500 |
| **EAAT2** | Rabbit | Polyclonal | Abcam | Ab41621 | 1:500 |

**Table 3-.**
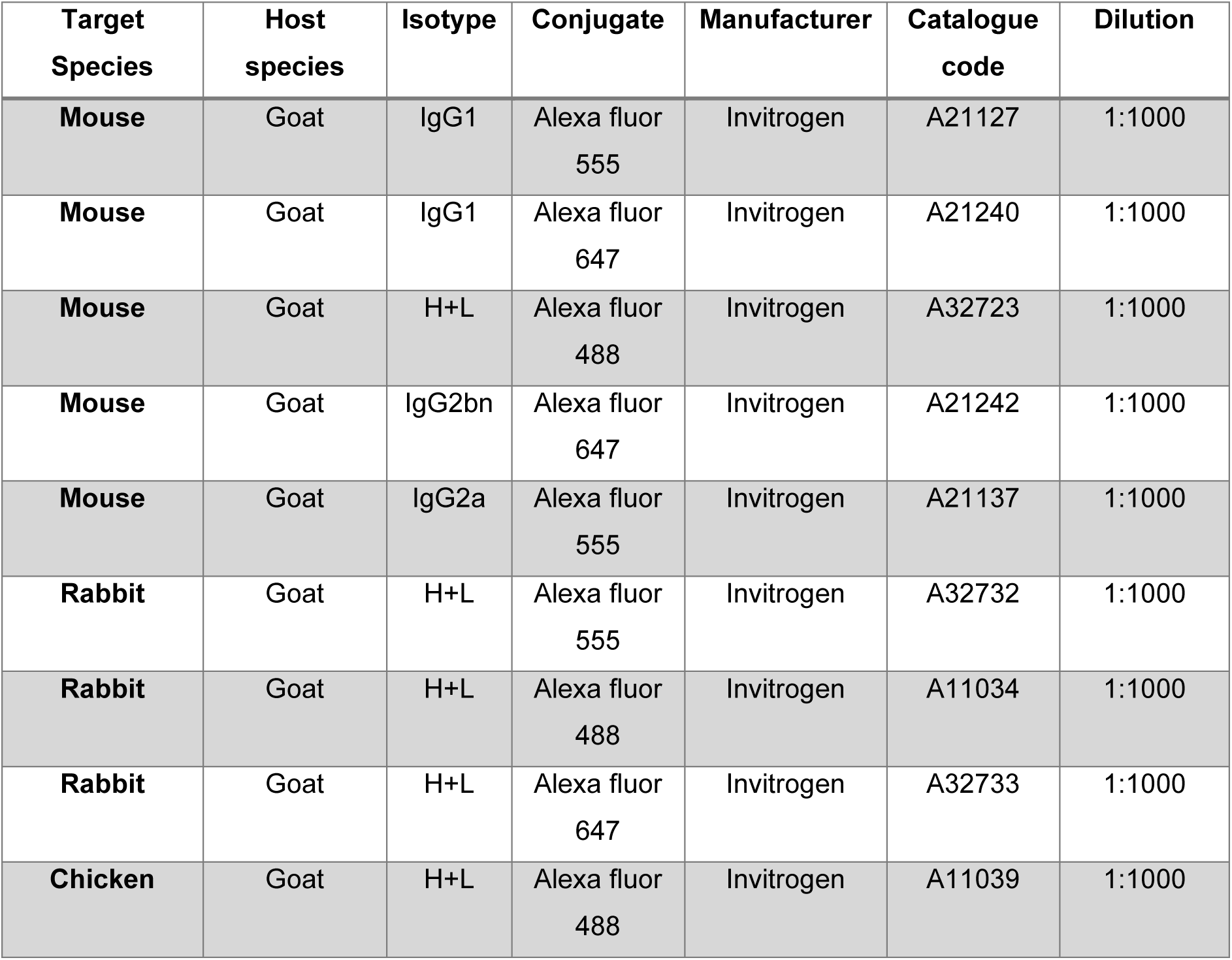
Secondary antibodies.

For Connexin-43, EAAT1, EAAT2 staining and synaptogenesis analysis, cells were fixed with 100% methanol for 10 minutes at −20°C to maintain membrane integrity. This was followed by 3 x PBS washes. In addition, the Triton-X permeabilization step was omitted. Triton-X was replaced by PBS as a diluent in the antibody staining solutions.

All samples were mounted with FluorSave (Millipore) for a minimum of 1 hour at room temperature before imaging.

Silicon rhodamine-tubulin (Spirochrome) was used at a concentration of 100 nM and incubated with the cells for 1 hour in the incubator, followed by 3 x PBS washes to remove dye residues.

Fluo-4AM (ThermoFisher) was used at a concentration of 5 μM and incubated with cells for 1 hour, followed by 2 x PBS washes.

Cell permeable Hoescht 33342 (Sigma) was used at a concentration of 1 μg/ml and incubated for 15 minutes in live cells, followed by 3 x PBS washes.

#### Astrocyte morphological analysis over differentiation

Astrocyte progenitors sparsely transfected with GFP were differentiated for 3 weeks in either NB27+BMP4, NB27+CNTF, NB27+BOTH or NB27 alone. Live imaging was performed on the Nikon Ti2 Eclipse microscope on days 0, 1, 7, 14 and 21 during the differentiation process. For each imaging timepoint imaging medium was used. 8×8 grids with 10% overlap were acquired in each well at each timepoint, using a 40x extra-long working distance objective. An imaging control was performed in parallel.

#### Manual process and cell body analysis

To assess subtle differences in shape between GFP^+^ astrocytes cultured in CNTF or NB27 alone, they were fixed with PFA, stained and imaged on the SoRa microscope at 60x magnification. Z-stacks of 35 slices with a step size of 0.3 μm were taken and images tiled to capture whole GFP^+^ cells.

Manual analysis of process length and cell body shape in GFP^+^ astrocytes was performed by a blinded operator using ImageJ after stitching the tiles. The segmented line tool was used to trace the primary processes starting from the cell body and ending at the process tip. Process number and length were recorded and averaged per cell. The cell soma was traced using the freehand selection tool, excluding the cell processes.

#### Cytokine switching experiments

Astrocytes were differentiated with BMP4 or CNTF for either 1 or 2 weeks; at the time of switching, a number of wells were changed to the alternate cytokine (i.e. BMP4 to CNTF and CNTF to BMP4) while a number of control wells were maintained in the same differentiation media. At the end of the differentiation, 2 or 3 weeks of differentiation respectively, cells were fixed with PFA and imaged on the Nikon Ti2 Eclipse microscope at 20x magnification, taking a 5×5 grid with 10% overlap in each well.

#### Glutamate uptake assay

To analyze glutamate uptake kinetics, a fluorometric glutamate uptake assay kit (Abcam) was used. 2-weeks differentiated BMP4 and CNTF astrocytes were incubated with 100 μM L-glut in Hanks’ Balanced Salt Solution (HBSS, Gibco) for different intervals of times (0, 60, 180 minutes). EAAT inhibitor PDC was used as a control and was incubated with the cells for 180 minutes (180 + PDC). Media was collected and enzyme mixes prepared according to the manufacturer’s instructions, before transferring to a clear bottom 96-well plate and incubating for 1 hour at RT. Glutamate-concentration standards were prepared in dilution buffer and used as a reference for quantification. A Tecan spark microplate reader was used to quantify fluorescence which correlated to L-glut concentration in the media. A blank well containing only enzyme mix was used as a control to subtract background fluorescence from all experimental wells. All experimental wells were then interpolated to the standard curve.

During the enzyme incubation period, the astrocytes were incubated with Hoechst 33342 and then PFA fixed; nuclei images were taken on a Nikon Ti2 widefield microscope and the number of nuclei used to calculate average number of cells per well.

Fluorescence intensity was normalised to the number of cells to standardise differences in confluency. All wells were also normalised to the average of the 0-minute timepoint of the corresponding experiment to produce a % to the t0 baseline.

#### Calcium imaging

To analyse calcium wave dynamics, 3 weeks differentiated BMP4 and CNTF astrocytes stained with Fluo-4AM were live imaged in a custom-built widefield microscope. A 5 mM ATP (Sigma) solution was prepared in Ca^2+^ free HBSS.

Timelapses were taken with 20x magnification, acquiring 400 images with a 400 ms interval. ATP was added at a final concentration of 0.5 mM at frame 110-115. An imaging control and a vehicle control were also performed.

To quantify calcium kinetics, pre-processing of the time-lapse videos was performed, including bleach correction with exponential fit and image registration to account for plate movements during the imaging, both on ImageJ. Approximately 30 circular, 5-pixel radius ROIs were drawn in each well, centering on the cell bodies. Fluo-4AM intensity in these ROIs was measured across every frame and the data input into PeakCaller software to perform peak analysis. To visualize intensity changes, a temporal color code with Fire LUT was applied to the timelapses on ImageJ.

### Scratch assay

To test wound healing capacity, astrocytes were differentiated with BMP4 or CNTF for 2 weeks, followed by either another week of standard differentiation or, in a subset of wells, one week of treatment with TICs. Cells were stained with silicon rhodamine-tubulin (SiR-Tubulin) and, in a subset of wells for each condition, a scratch was made with a pipette tip in the center of the well.

Following two PBS washes, cells were put in imaging medium and then imaged live on a custom built widefield microscope at 4x magnification. Imaging was then performed again, in the same positions, at 24 hours, 48 hours and 1 week after the scratch. Before each imaging session, cells were re-stained with SiR-Tubulin. At the end of the week, they were fixed with 4% PFA and stained using immunocytochemistry

The ImageJ Wound Healing plugin was used to quantify scratch areas at each time point and in each condition from the live images. The percentage of scratch area filled over the original total area was calculated, and every condition was normalized to untreated BMP4.

To assess nuclei presence in the scratch area, nuclei were identified from DAPI images using a pipeline including pixel classification using ilastik and object recognition using CellProfiler. At the same time, ilastik was used to produce a segmentation of the SiR-Tubulin image from the Day 0 timepoint. This was used to mask the nuclei and identify for each condition the nuclei that, at the one-week timepoint, had migrated or proliferated to the point of occupying the original scratch area.

#### Mitochondrial dynamics analysis

For the analysis of mitochondrial distribution and motility, astrocytes progenitors were co-transfected with mito-dendra2 and RFP, seeded on glass bottom chambers (Ibidi) and differentiated for 2-weeks with either BMP4 or CNTF. After PFA fixation, they were stained with DAPI.

For mitochondrial distribution, images were acquired on a CSU-W1 SoRa microscope with 60x magnification. Z-stacks composed of 35 slices with 0.3 μm step size (for a total of 10.5 μm) were taken and tiling was performed on ImageJ to capture whole cell morphology.

For the analysis of mitochondrial distribution, pixel classification was carried out with ilastik on the DAPI and RFP channels to generate binary segmentations of the nuclei and whole cells. These were uploaded onto CellProfiler together with the mito-dendra2 images. Nuclei with less than 95% overlap with the RFP cell objects were excluded, as well as RFP whole cells containing more than one nucleus (due to overlapping of non-fluorescent cells), to obtain one nucleus per RFP cell. The distribution of mito-dendra2 intensity was measured dividing the cell objects in 40 radial 25 μm bins (for a total of 1000 μm) starting from the edge of the nucleus. The fraction of total mito-dendra2 intensity in each bin was converted to a percentage and plotted.

For mitochondrial motility, 2-weeks differentiated BMP4 and CNTF astrocytes were changed to imaging medium and imaging performed on the CSU-W1 SoRa with 40x magnification. For each cell, an initial image was taken of both RFP and mito-dendra2, followed by a timelapse of only mito-dendra2 signal. For this, an image was taken every 10 seconds for 2 minutes with Z-stacks of 6 slices with a 2 μm step size (for a total of 12 μm). Low laser power was used to reduce toxicity. After the timelapse, another picture was taken with the RFP, mito-dendra2 and Hoechst channels to be used to locate the nucleus (without Z-stack).

For CNTF cells, an additional video was acquired at the end of processes (at a minimum distance of 150 μm from the nucleus).

The analysis of mitochondrial motility was carried out on the mito-dendra2 timelapse videos; 25×25 μm ROIs were established in the perinuclear (within 25 μm from the nucleus) and non-perinuclear (50-100 μm from the nucleus) areas, as well as at the process end for CNTF differentiated astrocytes (within 50 μm from the tip of an RFP^+^ process and at least 150 μm away from the nucleus). These ROIs were then bleach corrected, thresholded, contrast enhanced and binarized employing ImageJ. The ImageJ Total Motility plugin was used to quantify the percentage of pixels changing their intensity (0 to 1 or 1 to 0) between each frame. These percentages were averaged per compartment for each cell and plotted. In addition, temporal colour coding (Spectrum LUT) was used to visualise moving and stationary mitochondria on the timelapse.

#### Neuronal co-culture and synaptogenesis analysis

BMP4 and CNTF astrocytes were differentiated in their usual media for 2 weeks before plating cortical neuron progenitors on top of them and switching the culture to 50/50 medium with compound E for 2 additional weeks. At the end, they were fixed with methanol and stained for MAP2, PSD95 and synaptophysin.

Imaging was performed on the SoRa microscope at 60x magnification, acquiring 10.5 μm Z-stacks with 0.3 μm step size.

To quantify synaptic proteins and their co-localization, 150×150 μm ROIs were made. Computer software ilastik was used to produce binary segmentations for each of the three channels. For PSD95 and synaptophysin the training for pixel classification was carried out on the Z-stacks, followed by the creation of maximum intensity projections (MIP); for MAP2, training was performed directly on 2D MIPs as this enables more accurate dendrite segmentation. A CellProfiler pipeline was used to count the number of PSD95^+^, Synaptophysin^+^ and co-localised PSD95^+^/Synaptophysin^+^ puncta located within the MAP2 signal. The data is represented as number of puncta per 100 μm^2^ MAP2.

#### Imaging

Images for the scratch assay, calcium signaling and cell cycle assay were taken on a custom-built widefield microscope, at 4x magnification and 20x magnification, respectively.

Images for the live astrocyte shape characterisation during differentiation, cytokine switching experiments, cytoskeletal assays and reactive morphological analysis were taken on a Nikon Yi2 Eclipse widefield microscope, all at 20x magnification except for the live characterisation for which a 40x ELWD objective was used.

Images for the following experiments were acquired using the Olympus SpinSR10 SoRa spinning disk confocal microscope: manual analysis of processes, connexin-43 expression analysis, EAAT1 expression analysis, neuronal co-culture experiments, mitochondrial distribution and motility assays. Magnification ranging from 20x to 60x was used.

### Astrocyte morphological analysis

For all experiments which required single-cell morphology analysis of astrocytes transfected with GFP, a semi-automated image analysis pipeline was established which relies on open-source software.

GFP images were first pre-processed using a custom ImageJ macro which carried out contrast enhancement and gaussian blur to improve boundary detection, in addition to stitching tiles when required.

ilastik was then used to perform pixel classification. A subset of randomly selected images (around 10%) was used to train the algorithm in an operator-dependent manner by assigning two labels, corresponding to foreground pixels (GFP^+^ cells) and background pixels (GFP^−^).

After sufficient training, the software was able to perform accurate unsupervised pixel classification, and all images were batch processed in an automated manner. The results were binary segmentations of GFP^+^ cells which were batch thresholded on ImageJ and, if necessary, split into smaller tiles to allow faster downstream analysis. These were then input into CellProfiler where the GFP^+^ cells were identified as objects (after size-based filtering to exclude debris). Any objects touching the image border were excluded to limit the analysis to entire cells. Morphometric parameters were obtained for all cell objects and exported for further analysis.

#### Protein expression analysis

In all cases where protein expression was quantified from immunostaining images, the images from the channel corresponding to the protein of interest were pre-processed to subtract background and, in the case of Z-stacks, made into maximum intensity projections using ImageJ.

These pre-processed images were input into CellProfiler together with the DAPI channel images. Nuclei were identified as objects thanks to both size- and intensity-based thresholding and their number counted for each field of view. The whole image intensity of the staining of interest was measured and the data exported to excel. For each image, the lower quartile intensity was subtracted from the total intensity, and the result was then divided by the number of identified nuclei, providing a normalized intensity value per image (arbitrary units, a.u). The fields of view average for each well were plotted.

### Library prep and bulk RNA sequencing

The sequencing library was prepared with NEBNext UltraTM II Directional PolyA mRNA kit (NEB) and sequencing was performed on Illumina HiSeq 4000 yielding around 33 million strand specific 75 bp paired-ends reads per sample. Information on the number of reads and other statistics can be found in Table 4 and Table 5.

**Table 4-.** RNA sequencing statistics.

| Sample | Tot Sequences | Reads Mapped | %reads_mapped |
| --- | --- | --- | --- |
| CTRL2_BMP4_2_weeks_untreated_1 | 63318020 | 62742108 | 99.09044534241596 |
| CTRL2_BMP4_2_weeks_untreated_2 | 60343938 | 59872215 | 99.21827607604926 |
| CTRL2_BMP4_2_weeks_untreated_3 | 68280126 | 67632955 | 99.05218247546877 |
| CTRL2_BMP4_3_weeks_treated_1 | 72523052 | 71896908 | 99.13662762013932 |
| CTRL2_BMP4_3_weeks_treated_2 | 68719910 | 68097505 | 99.09428723058572 |
| CTRL2_BMP4_3_weeks_treated_3 | 68118536 | 67530234 | 99.13635548479786 |
| CTRL2_BMP4_3_weeks_untreated_1 | 66628114 | 66104610 | 99.21428963155104 |
| CTRL2_BMP4_3_weeks_untreated_2 | 62246314 | 61673077 | 99.07908281926541 |
| CTRL2_BMP4_3_weeks_untreated_3 | 75548322 | 74816579 | 99.03142388787933 |
| CTRL2_CNTF_2_weeks_untreated_1 | 67261926 | 66585345 | 98.99410998132882 |
| CTRL2_CNTF_2_weeks_untreated_2 | 71659442 | 71061694 | 99.16584893306873 |
| CTRL2_CNTF_2_weeks_untreated_3 | 71068294 | 70510948 | 99.21575998433282 |
| CTRL2_CNTF_3_weeks_treated_1 | 68734158 | 68098991 | 99.07590778954476 |
| CTRL2_CNTF_3_weeks_treated_2 | 59399876 | 58853905 | 99.08085498360299 |
| CTRL2_CNTF_3_weeks_treated_3 | 71788402 | 71182205 | 99.15557808349043 |
| CTRL2_CNTF_3_weeks_untreated_1 | 58906132 | 58378701 | 99.10462463 |
| CTRL2_CNTF_3_weeks_untreated_2 | 68983766 | 68423294 | 99.18753058509448 |
| CTRL2_CNTF_3_weeks_untreated_3 | 53404294 | 52966947 | 99.18106397961182 |
| CTRL2_progenitor_0_weeks_untreated_1 | 67729414 | 67172903 | 99.17833188398765 |
| CTRL2_progenitor_0_weeks_untreated_2 | 55756870 | 55285879 | 99.15527718826398 |
| CTRL2_progenitor_0_weeks_untreated_3 | 66857320 | 66328786 | 99.20945978690142 |
| CTRL3_BMP4_2_weeks_untreated_1 | 60670484 | 60124120 | 99.09945666495754 |
| CTRL3_BMP4_2_weeks_untreated_2 | 60575578 | 60062557 | 99.15308939 |
| CTRL3_BMP4_2_weeks_untreated_3 | 72381544 | 71709248 | 99.07117759190105 |
| CTRL3_BMP4_3_weeks_treated_1 | 59569862 | 58937206 | 98.93795960111508 |
| CTRL3_BMP4_3_weeks_treated_2 | 60456496 | 59835814 | 98.97334109472703 |
| CTRL3_BMP4_3_weeks_treated_3 | 63511056 | 62774236 | 98.83985553633372 |
| CTRL3_BMP4_3_weeks_untreated_1 | 63483212 | 62906589 | 99.09169214689389 |
| CTRL3_BMP4_3_weeks_untreated_2 | 63907994 | 63137193 | 98.79388954064183 |
| CTRL3_BMP4_3_weeks_untreated_3 | 70417194 | 69720828 | 99.01108527556495 |
| CTRL3_CNTF_2_weeks_untreated_1 | 67219916 | 66494396 | 98.92067702078057 |
| CTRL3_CNTF_2_weeks_untreated_2 | 57342148 | 56836702 | 99.11854365832266 |
| CTRL3_CNTF_2_weeks_untreated_3 | 68479538 | 67892401 | 99.14260957776906 |
| CTRL3_CNTF_3_weeks_treated_1 | 83891758 | 82793982 | 98.69143760224932 |
| CTRL3_CNTF_3_weeks_treated_2 | 62189738 | 61368150 | 98.67890101096744 |
| CTRL3_CNTF_3_weeks_treated_3 | 70108650 | 69162549 | 98.65052172592112 |
| CTRL3_CNTF_3_weeks_untreated_1 | 61146520 | 60582830 | 99.07813232870815 |
| CTRL3_CNTF_3_weeks_untreated_2 | 61434970 | 60895515 | 99.12190890627927 |
| CTRL3_CNTF_3_weeks_untreated_3 | 66441722 | 65868380 | 99.13707534551858 |
| CTRL3_progenitor_0_weeks_untreated_1 | 71789704 | 71124727 | 99.07371536174603 |
| CTRL3_progenitor_0_weeks_untreated_2 | 77861818 | 77130398 | 99.06061787563193 |
| CTRL3_progenitor_0_weeks_untreated_3 | 58143092 | 57661737 | 99.17212005168216 |

**Table 5-.** RNA sequencing statistics.

| Sample | %rRNA | %Exonic | %Sense | %Antisense |
| --- | --- | --- | --- | --- |
| CTRL2_BMP4_2_weeks_untreated_1 | 1.0702831391233224e-05 | 0,9083504 | 1,3 | 91,07 |
| CTRL2_BMP4_2_weeks_untreated_2 | 7.498254031548753e-06 | 0,90971877 | 1,27 | 90,16 |
| CTRL2_BMP4_2_weeks_untreated_3 | 3.602798417991563e-05 | 0,92046098 | 1,06 | 91,74 |
| CTRL2_BMP4_3_weeks_treated_1 | 3.412575402404687e-05 | 0,91035656 | 1,15 | 91,08 |
| CTRL2_BMP4_3_weeks_treated_2 | 2.9365221788994567e-05 | 0,9182632 | 0,88 | 91,7 |
| CTRL2_BMP4_3_weeks_treated_3 | 8.864763598916725e-05 | 0,92032158 | 0,98 | 93,42 |
| CTRL2_BMP4_3_weeks_untreated_1 | 2.3645756836494996e-05 | 0,91036341 | 1,34 | 90,51 |
| CTRL2_BMP4_3_weeks_untreated_2 | 6.841568114778179e-05 | 0,91692811 | 0,93 | 90,77 |
| CTRL2_BMP4_3_weeks_untreated_3 | 5.652362846828613e-05 | 0,914929 | 1,26 | 91,6 |
| CTRL2_CNTF_2_weeks_untreated_1 | 1.3677717281994113e-05 | 0,89461464 | 1,09 | 90,23 |
| CTRL2_CNTF_2_weeks_untreated_2 | 9.502154566037075e-06 | 0,91487257 | 1,14 | 90,76 |
| CTRL2_CNTF_2_weeks_untreated_3 | 9.525412817103375e-06 | 0,91915607 | 1,07 | 90,97 |
| CTRL2_CNTF_3_weeks_treated_1 | 1.6254286214639087e-05 | 0,92243262 | 0,69 | 93,29 |
| CTRL2_CNTF_3_weeks_treated_2 | 3.0267865692853577e-05 | 0,92125743 | 0,84 | 92,09 |
| CTRL2_CNTF_3_weeks_treated_3 | 1.864833452501371e-05 | 0,92318015 | 0,76 | 94,32 |
| CTRL2_CNTF_3_weeks_untreated_1 | 7.788432526550377e-06 | 0,89587303 | 1,18 | 88,65 |
| CTRL2_CNTF_3_weeks_untreated_2 | 1.964225559877943e-05 | 0,91908482 | 0,9 | 92,15 |
| CTRL2_CNTF_3_weeks_untreated_3 | 5.8648220516638876e-05 | 0,92558214 | 0,81 | 91,47 |
| CTRL2_progenitor_0_weeks_untreated_1 | 3.3703698785460256e-06 | 0,90843109 | 1,02 | 91,65 |
| CTRL2_progenitor_0_weeks_untreated_2 | 1.2248234478241135e-05 | 0,91232669 | 0,94 | 90,34 |
| CTRL2_progenitor_0_weeks_untreated_3 | 2.026570911641441e-05 | 0,91766338 | 0,85 | 93,12 |
| CTRL3_BMP4_2_weeks_untreated_1 | 1.8582752210409086e-05 | 0,91282913 | 1,98 | 90,41 |
| CTRL3_BMP4_2_weeks_untreated_2 | 0 | 0,89503774 | 2,42 | 90,45 |
| CTRL3_BMP4_2_weeks_untreated_3 | 6.267368641202202e-06 | 0,8993392 | 2,48 | 91,69 |
| CTRL3_BMP4_3_weeks_treated_1 | 0 | 0,90702329 | 1,9 | 91,73 |
| CTRL3_BMP4_3_weeks_treated_2 | 0 | 0,90361078 | 1,96 | 91,42 |
| CTRL3_BMP4_3_weeks_treated_3 | 3.571096586985146e-06 | 0,90903192 | 1,93 | 91,42 |
| CTRL3_BMP4_3_weeks_untreated_1 | 1.0904201465153934e-05 | 0,88758165 | 2,29 | 88,74 |
| CTRL3_BMP4_3_weeks_untreated_2 | 0 | 0,89868903 | 2,19 | 90,51 |
| CTRL3_BMP4_3_weeks_untreated_3 | 6.627272831501655e-06 | 0,87752893 | 2,32 | 89,03 |
| CTRL3_CNTF_2_weeks_untreated_1 | 1.358954702914156e-05 | 0,90195388 | 1,85 | 92,09 |
| CTRL3_CNTF_2_weeks_untreated_2 | 4.003537044908435e-06 | 0,89327581 | 1,78 | 91,18 |
| CTRL3_CNTF_2_weeks_untreated_3 | 3.315715719950804e-06 | 0,90856993 | 1,92 | 91,91 |
| CTRL3_CNTF_3_weeks_treated_1 | 8.006105135532151e-06 | 0,92146936 | 1,03 | 95,63 |
| CTRL3_CNTF_3_weeks_treated_2 | 3.6379577232932977e-06 | 0,91029995 | 1,17 | 94,74 |
| CTRL3_CNTF_3_weeks_treated_3 | 0 | 0,92149999 | 1,04 | 94,6 |
| CTRL3_CNTF_3_weeks_untreated_1 | 3.7903913579077043e-06 | 0,88722491 | 2,12 | 89,74 |
| CTRL3_CNTF_3_weeks_untreated_2 | 0 | 0,89913839 | 1,89 | 90,48 |
| CTRL3_CNTF_3_weeks_untreated_3 | 1.0311228625510474e-05 | 0,90097132 | 2,05 | 91,39 |
| CTRL3_progenitor_0_weeks_untreated_1 | 9.432035815326398e-06 | 0,91558161 | 1,53 | 93,48 |
| CTRL3_progenitor_0_weeks_untreated_2 | 1.1648714153775956e-05 | 0,90272594 | 2,1 | 91,8 |
| CTRL3_progenitor_0_weeks_untreated_3 | 0 | 0,90587083 | 1,85 | 93,26 |

### Bioinformatic analysis of bulk RNA-seq

#### Bulk RNA-seq data processing

Primary data analysis was done using the nf-core/rnaseq pipeline [https://github.com/nf-core/rnaseq] using default parameters (version 3.10.1) written in the Nextflow domain specific language (version 22.10.3) in conjunction with Singularity (version 3.6.4). Alignment of sequencing reads was carried out using the reference human genome (Ensembl GRCh38 release 95). Gene counts per gene per sample were obtained using RSEM-STAR (RSEM v1.3.1, STAR v2.7.10a)^76^ option of the pipeline and they were imported on DESeq^77^ (v1.38.3) within R environment (v4.2.2) for differential expression analysis. For further quality control (QC), PCA and heatmap plots (data not shown) with hierarchical clustering of top 500 variable genes post variable stabilizing transformation were studied for any batch effects.

#### Differential expression analysis

Differential gene expression analysis was carried out in R environment v4.2.2 with the DESeq2 (v1.38.3) package. The gene expression response was modelled as: ∼ cell_line_batch + target + treatment + target:treatment, where “target” refers to cell type and “cell_line_batch” to the 3 individual batch of experiments per sample group. The differential expression using DESeq2 was achieved by use of negative binomial generalized linear models along with Wald test *p* value for significance, IHW^78^ (v1.26.0) for multiple-testing correction, and ashr^79^ (v2.2-63) for effect-size shrinkage to ensure reported fold-changes are more robust.

Additional PCA plots were produced using data after regressing out the effect of all experimental factors from the above model except the one of interest thus masking variance contribution from other factors. The log2 fold changes for differential comparison of interest were visualized as volcano plots to represent log fold-change versus p-value between two experimental factors. Gene signatures were considered significant if FDR q-value <0.05.

#### Functional enrichment and pathway analysis

Gene Set Enrichment^80^ was performed using clusterProfiler package^81^ (version 4.6.2) for each pairwise comparison using gene lists ranked by the regularized log2 fold changes. Gene sets enrichments were quantified from MSigDB (version 7.5.1) for gene ontology (GO) Biological Process. Additionally, Venn diagrams were generated using the ggVennDiagram^82^ (version 1.2.2) package across selected differential comparisons to illustrate set overlaps using significant gene signatures showing either upregulated or downregulated characteristics separately.

#### Single-cell RNA-seq reference data

The Human Brain Cell Atlas v1.0 single-cell RNA sequencing (scRNA-seq) dataset from three pathology-free post-mortem donors across 10 brain regions was downloaded from the Chan Zuckerberg CELLxGENE explorer (https://cellxgene.cziscience.com/collections/283d65eb-dd53-496d-adb7-7570c7caa443)^41^. For the ∼2.5 million neuronal and 888k non-neuronal nuclei, the integrated Seurat object was randomly down-sampled to 50k cells and gene-cell expression counts were log normalized with Seurat *NormaliseData*^83^.

Data for cell type markers in the scRNA-seq dataset enriched for vascular cell types, containing glial and neuronal cell populations, in entorhinal and somatosensory cortex samples from 6 Alzheimer’s disease (AD) and 6 non-disease control (NDC) cases, and prefrontal cortex samples 10 AD and 10 NDC cases were accessed from GEO^42^.

The snRNA-seq dataset containing grey and white matter astrocytes was downloaded from zenodo (<u>10.5281/zenodo.8338962</u>), and processed into a Seurat object using a script provided by the authors of the publication. The dataset was subset to 9054 astrocyte cells from 26 control samples before downstream analysis^43^.

#### Comparative analysis

Gene expression levels in the differentiated astrocytes from bulk RNA-seq data were correlated with Human Brain Cell Atlas^41^ pseudo-bulk counts from scRNA-seq data in the form of gene effect sizes, which were derived by comparison with progenitor cells. The down-sampled raw gene-cell count matrix used to derive pseudo-bulk counts by donor and cell type (neuronal and non-neuronal, annotation level:’cell_type’) or subtype (astrocyte, annotation level: ‘cluster_id’) with Seurat *AggregateExpression*^83^, which were compared against progenitor cell gene expression with DESeq2 v1.38.3) differential analysis. The degree of similarity was quantified by Pearson’s correlation coefficient (*R*; p<0.05) of the gene log_2_(FoldChange) effect sizes. This was repeated at several adjusted *p*-value thresholds of differentiated astrocyte differentially expressed genes (n=8817 and 9005, 7472 and 7713, 5338 and 5538, 4654 and 4852 for CNTF and BMP4 astrocytes respectively, at p.adj<0.1, 0.05, 0.01, 0.005) to ensure the robustness of the associations. Hierarchical clustering of the cell type correlation coefficients was carried out with the ComplexHeatmap R package using Euclidean distance (v2.14.0). These steps were replicated for the grey/white matter astrocyte comparisons, with pseudo-bulks derived for each individual and matter-type (annotation level:’matter’) or subtype (annotation level:’fine_type’) at the same adjusted p-value thresholds of differentiated astrocyte differentially expressed genes.

As a complimentary approach, we tested the enrichment of cell type marker genes in the single cell human reference datasets. The derive the marker gene lists for the down-sampled Human Brain Cell Atlas dataset^41^, we used Seurat *FindAllMarkers*^83^ with log_2_ FoldChange threshold=0.1, minimum proportion of cells expressing gene=0.2, and considering only upregulated genes. This was repeated using the full astrocyte dataset for astrocyte subtype comparisons. Cell type marker gene list for a human post-mortem snRNA-seq dataset enriched in vascular cell types were downloaded from a previous publication^42^. The intersection between the top reference cell type markers and differentiated astrocyte upregulated genes (Human Brain Cell Atlas cell type Nmarker= 50 and astroDEG=1000; Human Brain Cell Atlas astrocyte cluster Nmarker=20 and astroDEG=500; Vascular-enriched cell type Nmarker=100 and astroDEG=1000) was quantified and tested for enrichment with a hypergeometric test using *phyper* in R.

### Mouse astrocyte isolation and bulk RNA sequencing

Primary mouse cortical astrocytes were isolated from C57BL/6N mice according to previously established protocol^44^. Briefly, cortices were isolated from postnatal day 2-3 mouse pups and dissociated with Papain (Worthington) in DMEM (Gibco) with DNase (Worthington, 12.5U/ml) and filtered through a 70μm filter. Following dissociation, cells were plated in 10 cm dishes coated with poly-D-lysine (Sigma-Aldrich, 2μg/ml) and laminin (Gibco, 6μg/ml) in survival medium containing 50% DMEM/F-12 (Gibco) and 50% neurobasal (Gibco) with B27 (Gibco, 1X), N2 (Gibco, 1X), L-glutamine (VWR, 200μg/ml), non-essential amino acids (Gibco, 1X), penicillin-streptomycin (VWR, 1X), glucose (Alfa Aesar, 3mg/ml), and N-acetyl-l-cysteine (Sigma-Aldrich, 5μg/ml). Survival medium was supplemented with Fgf2 (Stemcell Technologies, 20ng/ml), CNTF (Stemcell Technologies, 10ng/ml), BMP4 (Stemcell Technologies, 10ng/ml) and hbEGF (Sigma-Aldrich, 5ng/ml). After 24 hours in survival medium, B-27 was removed from the medium. From this point, media was changed every 48 hours while the cells grew, and once they were confluent (5 days), they were cryopreserved in Cryostor media (Stemcell Technologies) and stored at −80C.

Cells were thawed on day 0 of an experiment, and they were plated on poly-D-lysine (Sigma-Aldrich, 2μg/ml) and laminin (Gibco, 6μg/ml) coated plates in growth media containing 50% DMEM/F-12 (Gibco) and 50% neurobasal (Gibco) with N2 (Gibco, 1X), L-glutamine (VWR, 200μg/ml), non-essential amino acids (Gibco, 1X), Pen-Strep (VWR, 1X), glucose (Alfa Aesar, 3mg/ml), and N-acetyl-l-cysteine (Sigma-Aldrich, 5μg/ml). Growth media was supplemented with either BMP4 (Stemcell Technologies, 10ng/ml) or CNTF (Stemcell technologies, 20ng/ml). Media was changed every 48 hours. On Day 7, cells were imaged on the Olympus CKX53 microscope with the Olympus DP23M camera. Post-processing was done in FIJI. Then, cells were lysed to isolate RNA according to manufacturer instructions with the RNeasy Plus Mini Kit (Qiagen).

Cell lysates were then sequenced on the NovaSeq X+. Raw sequencing FastQ data quality was assessed using *fastqc*. Adapter and quality trimming was performed using *trimgalore.* Files were then aligned to the mouse genome (GRCm38 - mm10) using *salmon* (V1.10.0). Differential gene expression testing between conditions was then generated using DESeq2(V1.42.1)^77^. Volcano plots were generated using *EnhancedVolcano* (V1.20.0) and Gene set enrichment go term analysis was performed using clusterProfiler(V4.10.1)^81^.

#### Multivariate analysis of astrocyte morphometric parameters

We performed singular value decomposition (SVD) of matrix of 52 morphological measurements across the 5,376 cells at distinct stages of astrocyte differentiation in four different culture conditions (basal, CNTF, BMP4, CNTF+BMP4). We next quantified the association between individual left singular vectors and culture conditions using linear mixed modelling, accounting for the idiosyncratic differences that may arise from the different cell lines and experiments using R and lme4 37. As fixed effects, we entered the culture condition variable into the model. As random effects, we had intercepts for either cell lines and experiments. Visual inspection of residual plots did not reveal any obvious deviations from homoscedasticity or normality. P-values were obtained by likelihood ratio tests of the full model with the effect in question against the model without the effect in question. We iteratively fitted a linear mixed model to the each left singular vectors.

To visualize the right singular vectors, we plotted the average singular vectors per group on the vertical axis as a function of the time corresponding to each sample on the horizontal axis and coloring all samples corresponding to culture conditions. Next, we identified morphological features whose profiles correlated (Pearson correlation between individual morphological measurements profile and right singular vectors) and contributed (projection of each individual morphological measurements profile onto right singular vectors) most strongly (either positively or negatively) with the profile of the singular vectors. In order to identify representative measurements for each singular vector, features were ranked according to both projection and correlation scores. The highest (most positive scores in both projection and correlation) and lowest (most negative scores in both correlation and projection) motifs were selected for each singular vector using K-mean clustering.

#### Statistical analysis and plotting

For every experiment, Gaussian distribution of the data was tested using Shapiro-Wilks or Kolmogorov-Smirnov tests, and parametric or non-parametric statistical test used in accordance.

Statistical tests are specified in figure legends but, briefly, unpaired T-test was used for comparison between two groups with normal distribution, while Mann-Whitney U-test was used in case of non-normal distribution. For comparison of 3 or more groups, one-way ANOVA, Kruskal-Wallis or two-way ANOVA were used based on the normality of the data. Tukey, Dunn’s or Šídák’s tests were used for multiple comparisons.

Plotting was performed either on GraphPad Prism or using custom R scripts. Statistical significance was represented as *, **, ***, **** for p-values of <0.05, <0.01, <0.001, <0.0001, respectively.

Figures were assembled using Adobe Illustration software.

## Data availability

For RNA sequencing, raw and transformed gene expression counts, differential analysis, and gene set enrichment analysis output tables, as well as Human Brain Cell Atlas reference dataset cell type marker gene lists can be found on the GitHub repository: https://github.com/Marzi-lab/AstroShape_RNA.

Raw RNA sequencing data can be accessed on GEO, under accession number GSE279716.

R scripts used to perform comparative analysis and reproduce the figures in this manuscript are available on the GitHub repository: https://github.com/Marzi-lab/AstroShape_RNA.

For multivariate analysis of astrocyte morphometric parameters, we provide the raw images and complete source code (which is not a software but rather a compilation of R to readily reproduce figures, tables, and other results that involve computation to facilitate the development and evaluation of additional profiling methods. We also provide the >60 measurements of each of the ∼500 cells whose origins are annotated. The raw images, metadata and single-cell measurements provided as comma-delimited files have been deposited on Github together with the statistical analysis in the following repository: https://github.com/RLuisier/AstroShape.

## CRediT (Contributor Roles Taxonomy) Statement

Kelly O’Toole: conceptualization, methodology, validation, formal analysis, investigation, visualization, writing – review & editing.

Ludovica Guetta: conceptualization, methodology, validation, formal analysis, investigation, writing-original draft, writing – review & editing, visualization.

Paulina Urbanaviciute: software, formal analysis, data curation, visualization, writing – review & editing.

Pacharaporn Suklai: validation, investigation, formal analysis, visualization.

Chandler Wright: validation, formal analysis, investigation, writing – review & editing.

Theodore Fisher: validation, formal analysis, investigation, writing – review & editing.

Shane Liddelow: conceptualization, supervision, funding acquisition, validation, writing – review & editing.

Raphaelle Luisier: software, formal analysis, data curation, visualization, writing – review & editing.

Sarah J Marzi: software, formal analysis, data curation, visualization, writing – review & editing.

Andrea Serio: conceptualization, writing-original draft, writing – review & editing, supervision, project administration, visualization, funding acquisition.

## Notes

### Competing Interest Statement

The authors have declared no competing interest.

### Summary of Updates

References 12-13 adjusted and formatted, removed an editorial comment left by accident

